# GCAT|Panel, a comprehensive structural variant haplotype map of the Iberian population from high-coverage whole-genome sequencing

**DOI:** 10.1101/2021.07.20.453041

**Authors:** Jordi Valls-Margarit, Iván Galván-Femenía, Daniel Matías-Sánchez, Natalia Blay, Montserrat Puiggròs, Anna Carreras, Cecilia Salvoro, Beatriz Cortés, Ramon Amela, Xavier Farre, Jon Lerga-Jaso, Marta Puig, Jose Francisco Sánchez-Herrero, Victor Moreno, Manuel Perucho, Lauro Sumoy, Lluís Armengol, Olivier Delaneau, Mario Cáceres, Rafael de Cid, David Torrents

**Affiliations:** Life sciences dept, Barcelona Supercomputing Center (BSC), Barcelona, 08034, Spain.; Genomes for Life-GCAT lab Group, Institute for Health Science Research Germans Trias i Pujol (IGTP), Badalona, 08916, Spain.; Institute for Research in Biomedicine (IRB Barcelona), The Barcelona Institute of Science and Technology, 08028, Barcelona, Spain (current affiliation).; Institut de Biotecnologia i de Biomedicina, Universitat Autònoma de Barcelona, Bellaterra, Barcelona, 08193, Spain.; High Content Genomics and Bioinformatics Unit, Institute for Health Science Research Germans Trias i Pujol (IGTP), 08916, Badalona, Spain.; Catalan Institute of Oncology, Hospitalet del Llobregat, 08908, Spain.; Bellvitge Biomedical Research Institute (IDIBELL), Hospitalet del Llobregat, 08908, Spain.; CIBER Epidemiología y Salud Pública (CIBERESP), Madrid, 28029, Spain.; Universitat de Barcelona (UB), Barcelona, 08007, Spain.; Sanford Burnham Prebys Medical Discovery Institute (SBP), La Jolla, CA 92037, USA.; Cancer Genetics and Epigenetics, Program of Predictive and Personalized Medicine of Cancer (PMPPC), Health Science Research Institute Germans Trias i Pujol (IGTP), Badalona, 08916, Spain.; Quantitative Genomic Medicine Laboratories (qGenomics), Esplugues del Llobregat, 08950, Spain.; Department of Computational Biology, University of Lausanne, Génopode, 1015 Lausanne, Switzerland.; Swiss Institute of Bioinformatics (SIB), University of Lausanne, Quartier Sorge – Batiment Amphipole, 1015 Lausanne, Switzerland.; ICREA, Barcelona, 08010, Spain.

**Author notes:** These authors contributed equally to this work (Jordi Valls-Margarit, Iván Galván-Femenía, Daniel Matías-Sánchez). Lead Contact for Data Access (Rafael de Cid).

**Keywords:** Complex Diseases, Reference Haplotype panel, Germline Structural Variation, GWAS, Variant calling, Genotype calling, Phasing, Imputation, Personalised medicine.

## Abstract

The combined analysis of haplotype panels with phenotype clinical cohorts is a common approach to explore the genetic architecture of human diseases. However, genetic studies are mainly based on single nucleotide variants (SNVs) and small insertions and deletions (indels). Here, we contribute to fill this gap by generating a dense haplotype map focused on the identification, characterization and phasing of structural variants (SVs). By integrating multiple variant identification methods and Logistic Regression models, we present a catalogue of 35,431,441 variants, including 89,178 SVs (≥50bp), 30,325,064 SNVs and 5,017,199 indels, across 785 Illumina high coverage (30X) whole-genomes from the Iberian GCAT Cohort, containing 3.52M SNVs, 606,336 indels and 6,393 SVs in median per individual. The haplotype panel is able to impute up to 14,360,728 SNVs/indels and 23,179 SVs, showing a 2.7-fold increase for SVs compared with available genetic variation panels. The value of this panel for SVs analysis is shown through an imputed rare Alu element located in a new locus associated with mononeuritis of lower limb, a rare neuromuscular disease. This study represents the first deep characterization of genetic variation within the Iberian population and the first operational haplotype panel to systematically include the SVs into genome-wide genetic studies.

## INTRODUCTION

One of the central aims of biology and biomedicine has been the characterization of genetic variation across humans to answer evolutionary questions, and to explain phenotypic variability in relation to disease. From the first genotyping and sequencing efforts, scientists have been gradually identifying specific genomic regions that vary within and across different populations, elaborating the first maps of human genetic variation (e.g. the HapMap Phase I^1^). Next-generation sequencing (NGS) technologies are now allowing to systematically evaluate the genetic variability across the entire genome of hundreds and thousands of individuals. This has increased more than 200-fold the number of known genomic variants over the past 10 years, resulting in much richer reference catalogues of genetic variability. One example is HRC^2^ or Trans-Omics for Precision Medicine (TOPMed)^3^, listing more than 39.2M and 410M polymorphic positions, respectively, from several human populations. The extensive genetic and phenotypic characterization of cohorts using rich variability reference panels is now fuelling up Genome-Wide Association Studies (GWAS). A total of 151,703 unique genetic variants are already reported to be associated across 5,193 unique traits (GWAS catalog, version1.0.2 release 05/05/2021, https://www.ebi.ac.uk/gwas/). Despite these advances, a large fraction of the genetic variability underlying complex diseases still remains unexplored, as studies have been mostly restricted to single nucleotide variants (SNVs) and small insertions and deletions (indels) (<50bp). Large structural variants (SVs) are known to play an important role in disease^4-7^ and could actually explain part of the well-known missing heritability paradox^8, 9^. However, the technical and methodological challenges associated with the identification and classification of this type of variation from whole-genome sequences (WGS) have left this type of variation out from GWASs.

Large-scale efforts combining improved sequencing methodologies are now identifying a much larger and richer spectrum of structural variation in humans. For example, by increasing the sequencing coverage and sample size across different populations, the gnomAD-SV project^10^ detected a median of 7,439 SVs per individual, generating one of the most extensive catalogues of structural variation so far. Other whole-genome studies have gone a step further by phasing the variants and constructing haplotype panels, as the 1000 Genomes project (1000G)^11^, becoming a reference within the GWAS community. However, the SVs are less represented in the current 1000G phase3, including a median of 3,441 SVs per individual^11^. The use of costly family trios and an increase in the sequencing coverage, allowed the Genome of the Netherlands consortium (GoNL) to increase a median of 7,006 SVs per individual^12^. In parallel, the recent inclusion of long-read sequencing technologies has made possible to uncover many new SVs, reaching >20,000 per individual^13-16^, including repeat-rich regions, where short-read sequencing has traditionally shown low call rates.

Genome-wide imputation from SNP-genotyping array data is still the most practical and powerful strategy to predict SVs, and test them for association with particular phenotypes. Current haplotype reference panels allow a high-quality imputation (info score ≥ 0.7) of approximately 9,000 to 14,000 SVs (≥50 bp), but considering the ranges of SVs that the community is now reporting across individuals, this is still incomplete. Therefore, it is necessary to generate improved variability reference panels of controlled populations by including SVs in the discovery and functional interpretation of associated variants to power-up current genetic studies.

In this study, we contribute to fill this gap by generating a new SV-enriched haplotype reference panel of human variation, through the analysis of whole-genome sequences (30X) of Iberian individuals from the GCAT|Genomes for Life Cohort (www.genomesforlife.com)^17,18^. We developed and applied a comprehensive genomic analysis pipeline based on the weighted integration and orthogonal validation of the results of multiple variant callers to generate a robust catalogue of genetic variability that covers from SNVs to large SVs. These variants were further phased and converted into haplotypes that can be incorporated into GWAS. This study represents an important step towards the completion of the annotation and characterization of the human genome and provides a unique resource for the incorporation of SVs into genetic studies.

## MATERIAL AND METHODS

### Benchmarking samples

An *in-silico* genome with synthetic next-generation sequencing reads was generated with the ART software (ART-Illumina version 2.5.8)^36^, by inserting a controlled set of 5,334,669 changes, covering from single nucleotide variants (SNVs) to large structural variation (SVs), to the hg19 reference genome. The majority of these variants were extracted from real samples of the 1000G^11^ and the PanCancer projects^37^. In addition, to benchmark a wider and more complex range of variants, we designed and randomly inserted an additional set of 3,925 Structural Variants (SVs) (Supplementary Table. 2). These additional variants were inserted randomly in the hg19 reference genome (Supplementary Fig. 1), excluding telomeres and centromeres. Simulated FASTQ files were aligned to the hs37d5 reference genome using BWA^38^ (version 0.7.15-r1140) and Samtools^39^ (version 1.5). Best Practices of GATK^40^ were followed for marking duplicates (PICARD version 1.108) and for recalibrating Base Quality Scores of the BAM file with the VariantRecalibrator and ApplyVQSR modules of GATK4 (version 4.0.11).

The sample NA12878 from the genome in a Bottle (GIAB) Consortium^20^ and the *in-silico* were used to validate SNVs and indels detection. BAM files were reconstructed using the hs37d5 reference genome and following the GATK Best Practices guidelines.

### Variant calling

We originally selected 17 candidate programs for variant identification and classification, representing different calling algorithms and strategies: Split Read, Discordant Read, *de novo* Assembly and Read-depth. Variant callers were Haplotype Caller^41^ (version 4.0.2.0), Deepvariant^42^ (version 0.6.1), Strelka2^43^ (version 2.9.2), Platypus^44^ (version 0.8.1), and Varscan2^45^ (version 2.4.3) for SNVs and indels, and Delly2^46^ (version 0.7.7), Manta^47^ (version 1.2), Pindel^48^ (version 0.2.5b9), Lumpy^49^ (version 0.2.13), Whamg^50^ (version v1.7.0-311-g4e8c), SvABA^51^ (version 7.0.2), CNVnator^52^ (version v0.3.3), PopIns^53^ (version damp v1-151-g4010f61), Genome Strip^54^ (Version 2.0), Pamir^55^ (version 1.2.2), AsmVar^56^ (version 2.0) and MELT^57^ (version 2.1.4) (Supplementary information section 3) for SVs.

Recall, precision and F-score metrics were calculated to evaluate the performance of each variant caller for each variant type. The NA12878 sample was used as a gold standard to calculate performance metrics for SNVs and indels, and the *in-silico* was used to benchmark SVs. For SNVs and indels, a variant was considered a true positive when the calling matched with the exact position and alternative allele shown on the benchmarking set. The criteria to classify SVs as true positives were: (i) the chromosome and the breakpoint position ± the breakpoint-error of the variant caller overlaps with the gold standard (Supplementary Table 4), (ii) the SV type matched with the gold standard, and (iii) the length of the variant reported by the caller has a ≥80% reciprocal overlap with the length of the variant in the gold standard sample. In addition, from the benchmarking results of SVs, we also captured information from the callers regarding breakpoint resolution, the effect of the variant size on variant calling, and the genotyping accuracy. Platypus, Varscan, Genome Strip, Pamir and AsmVar (Supplementary Information section 4.2) were finally discarded due to technical incompatibilities with our computing environment, leaving 12 final variant callers to be applied to the GCAT-WG samples.

### Logistic regression modelling

Logistic Regression Modelling (LRM) was used on indels and SVs to merge and filter the results from all the different callers, generating a final set of high-quality variants with the highest recall and precision values. This method is proposed as an improved alternative to other simple strategies based on the number of coincident callers, which were also calculated for comparison and evaluation purposes. As discriminative variables, LRM used variant and calling related parameters, like size, reciprocal overlap and breakpoint resolution (Supplementary Table 5).

#### Logistic Regression Model for indels

LRM was trained using indels of the NA12878 sample and tested using the *in-silico* sample. Input for LRM was a merged dataset of the VCF outputs from the callers. We used as dependent variable the presence or absence of the variant in the training set and as independent variables the genotype reported by each variant caller. The prediction of the LRM was converted in a binary variable (PASS, if predicted probability ≥0.5; NO PASS otherwise), indicating if the variant is considered as true or false positive. The genotype considered in the LRM is a consensus genotype reported by Haplotype caller, Deepvariant and Strelka2 (Supplementary information section 5.1). The LRM was developed using R software (version 3.3.1) and the *ISLR* package.

#### Logistic Regression Model for SVs

For SVs, the LRM was trained using 10-fold cross-validations in a random subset of variants from the *in-silico* (70%) and tested in the remaining subset of the *in-silico* (30%). Similar to the LRM for indels, the input of the LRM for SVs is a merged dataset of the VCF outputs from the callers, where the dependent variable is the presence or absence of the variant in the training set and the prediction is a binary variable depending on the predicted probability (PASS, if predicted probability ≥0.5; NO PASS otherwise). The independent variables are the genotype reported by the variant callers, the size, number of callers/strategies and reciprocal overlap (Supplementary Table 5). Using stepwise backward criteria, we fitted a LRM for each SV type using the *caret* (version 6.0-85) and *e1071* (version 1.7-3) R packages.

The strategy to determine the position of a variant in the LRM was different for each SV type. First, variant callers were ranked according to the accuracy in resolving the breakpoint (with an interval of error of ±10 bp; Supplementary Table 6) and the number of variants detected. This was used to select unique variants according to the position of the caller for that particular variant. In the case that a variant was not detected by the best-ranked algorithms (Supplementary Table 6), the final position of the variant was considered as the median of the position and the length reported by the rest of the callers.

The strategy to determine the genotype of a variant in the LRM was adapted to each SV type. For Deletions and Insertions, we selected the final genotype based on the highest recurrency across callers that identified a particular variant. For Inversions, we directly reported the genotype obtained from the caller with the smallest genotyping error in the benchmarking analysis. For Duplications and Translocations, which show lowest genotyping accuracy in the benchmarking, we applied a customized genotyping method strategy. This is based on the information of the *in-silico* sample of the proportion of altered reads around the breakpoint. If the proportion of altered reads was < 0.20, genotype was 0/0, if the proportion was between 0.20 and 0.80, the genotype was 0/1, and if the proportion was >0.80, the genotype was 1/1 (Supplementary information section 5.2.3).

### Quality control

The GCAT Cohort is a prospective cohort study that includes 19,267 volunteers from Catalonia, in the North-east of Spain (http://www.genomesforlife.org/). The participants were recruited from the general population (2014-2017) with the only restriction to live at least five years in Catalonia and aged between 40-65 years. All participants who agree to be part of the study provided informed consent and are asked to sign a consent agreement. Whole-genome sequencing data from 808 individuals using HiSeq 4000 sequencer (Illumina, 30X coverage, read length 150 bp, insert size 600 bp) was obtained in FASTQ format. BAM files were built using the hs37d5 reference genome and following the GATK Best Practices. FASTQ and BAM files are deposited to the European Genome-Phenome Archive (EGA, EGAS00001003018).

Quality control was applied by assessing the quality alignment of the BAM files, the presence of contamination traces, possible swapped samples, population structure and relatedness. Alignment quality was analysed using PICARD (version 2.18.11), Biobambam^58^ (version 2-2.0.65) and Alfred^59^ (version 0.1.16). Contamination or swapped ID samples was determined by VerifyBamID^60^ (Supplementary information section 6.3.1, Supplementary Table 9). Population structure was assessed using reference ancestries genotypes. Identity by descent (IBD) estimates were used to remove up to third-degree relatives.

The GCAT sample was characterized by Principal Component Analysis (PC). Firstly, we ran the Haplotype Caller tool and only PASS variants from the VCF file were retained. Then, SNVs with minor allele frequency (MAF) > 0.01 and independent variants (LD, r² < 0.2, were selected) with PLINK (version 1.90b6.7 64-bit). Finally, on retained variants (∼1M) we ran PCs together with reference samples of known ancestry (i.e. 1000G project sample and the Population Reference Sample^19^ (POPRES)). Genetic homogeneity of the GCAT sample was confirmed by PCA in the retained cohort samples (Supplementary Fig. 7).

### Variant calling, filtering and merging

Each of the 12 selected variant callers was first executed independently with different parameters (Supplementary information section 7, Supplementary Fig. 8) on all samples. The results of all these callers were then merged according to the different criteria obtained through our benchmarking strategy. For SNVs and indels, we merged calls that agreed with (i) the chromosome, (ii) position at base-pair resolution, and (iii) the same REF/ALT allele. For SVs, we applied benchmarking rules and merged different calls into a single merged variant when they matched in (i) variant type, (ii) chromosome, (iii) position, considering the breakpoint error estimated for each variant caller (Supplementary Table 4) and (iv) reciprocal overlap ≥ 80% between callers (Supplementary information section 8.2). Then, we applied the LRM to analyse the credibility of each of the variants, considering a variant as true positive if the LRM prediction probability was ≥0.5. Given the consistently high accuracy in detecting SNVs for most callers, we considered one of these variants as a true positive if it was detected by at least two callers.

Variant VCF files were generated with these results applying different criteria. For SNVs and indels (<30bp), final variants were reported by merging calls on different individuals that coincided in chromosome and position at base-pair resolution, and in REF/ALT alleles. For SVs, we merged individuals by (i) variant type, (ii) chromosome, (iii) position, considering the maximum breakpoint-error obtained for that merged variant (iv) reciprocal overlap (≥80%) between individuals (Supplementary information section 8.3). We calculated the true positive proportion for each variant determined by the LRM prediction in all GCAT samples. We referred to this proportion as the quality score of the merged variant. Then, we considered a variant as PASS if the quality score was ≥0.5. We reported the length and position of each SV as the median length and median position of all the samples that have that SV (Supplementary methods). Finally, monomorphic variants, variants out of Hardy-Weinberg equilibrium (Bonferroni correction p-value < 5×10^-8^), and variants with ≥10% of missingness were excluded from subsequent analysis.

### Variant validation

#### Comparison with public datasets

SNVs and indels from the GCAT dataset were compared with the NCBI dbSNP database (Build version 153) (https://ftp.ncbi.nlm.nih.gov) to determine the number of unique/shared variants between them. GCAT SVs were compared with the following public databases: (i) The Genome Aggregation Database (gnomAD.v.2) (https://gnomad.broadinstitute.org/downloads), (ii) the Database of Genomic Variants (DGV) (http://dgv.tcag.ca/dgv/app/downloads?ref=GRCh37/hg19), (iii) the Human Genome Structural Variation Consortium set (HGSVC) (ftp://ftp.1000genomes.ebi.ac.uk/vol1/ftp/data_collections/hgsv_sv_discovery/) (iv) the Ira M Hall dataset (https://github.com/hall-lab/sv_paper_042020), (v) the 1000G project (Phase3) (ftp://ftp.1000genomes.ebi.ac.uk/vol1/ftp/phase3/) and (vi) GoNL (release 6.2) (on request). Finally, we determine the number of shared variants between the GCAT and at least one other dataset and the number of unique variants in the GCAT (Supplementary Information section 9.1.2).

#### Experimental validation

The validation of SNV and indels was performed using genotypes calls generated by WGS and genotyping data from SNP-array techniques in 570 GCAT samples. We include QCed genotypes generated in the GCAT cohort with the Infinium Expanded Multi-Ethnic Genotyping Array (MEGAEx) (ILLUMINA, San Diego, California, USA) as described elsewhere^18^ (i.e. 732,978 SNPs and 1,168 indels). Genotypes from both strategies were compared by (i) chromosome and position at base-pair resolution and (ii) REF/ALT alleles; the recall and genotype concordance for each individual sample was calculated.

Inversions were validated using the most recent benchmark set of validated human polymorphic inversions from the InvFEST porject^24^. We considered 59 experimentally genotyped inversions generated by non-homologous mechanisms^24^. Allele frequency (using CEU and TSI European populations) and length concordance was determined using an overlapping window of ±1Kb around the inversion breakpoints. Accuracy of inversion genotyping was compared for the 785 WGS samples using the available reference panel of experimentally-resolved genotypes^24^. Genotypes in GCAT samples were imputed with IMPUTE2^30^ with a posterior probability ≥ 0.8 and were classified as missing otherwise. Missing genotypes were recovered if they had a perfect tag SNP in the reference panel (r^2^ = 1).

### Phasing and imputation performance

In order to analyse the performance of the phasing and imputation processes, the 785 GCAT individuals were divided into two subsets, (i) a subset including 690 individuals used to construct a pilot reference panel; and (ii) the remaining 95 individuals, with WGS and SNP-genotyping array data available, which was used as a test sample in the different analyses.

The evaluation of phasing strategies was carried out by determining the imputation accuracy of SVs, using the genotypes independently generated by WGS and imputation techniques across the 95 test GCAT samples, and with the pilot reference panel of 690 individuals (Supplementary Information section 10.1). Accuracy was determined only for chromosome 22 to reduce computational efforts, and the quality score of imputed variants was considered as a validation proxy of the best phasing strategy. Each phasing strategy was evaluated by counting the number of variants with an info score ≥0.7, and by calculating the genotype concordance between imputed data and WGS calling. The phasing algorithms evaluated were ShapeIt2^61^ (version v2.r904), MVNcall^62^ (version 1.0), ShapeIt4^27^ (version 4.1.3) and WhatsHap^28^ (version 0.18). We used IMPUTE2^30^ (version 2.3.2) for imputation analysis (Supplementary methods).

In order to evaluate the imputation performance of the GCAT|Panel for distant ethnicities, we used the 1000G SNP-genotyping array data covering 2,318 samples from 19 populations^63^. First, quality control was applied to the 1000G SNP-genotyping array per population by removing variants (i) with ≥10% of missingness, (ii) in A-T, C-G sites and (iii) in Hardy-Weinberg Disequilibrium (p-value < 0.05); and by discarding samples with (i) ≥ 10% of missing, (ii) Kinship coefficient ≥ 0.05, and (iii) an excess of heterozygosity ± 2SD, obtaining finally 1,880 individuals from 19 populations. Each population was pre-phased with Shapeit4 and imputed separately using IMPUTE2. Then, we compared the allele frequency, type of variant distribution and the quality of the imputed SVs across populations.

To evaluate the imputation of SVs, we used as reference the SV characterisation performed by Audano et al.^15^. The imputed SVs with an info score ≥ 0.7 were compared considering a window of ±50bp. Furthermore, we evaluated the concordance of SV type and SV length error reported by WGS calling. On other hand, the genotype characterisation of the same samples reported by Hickey et al.^32^, was used to evaluate the genotype accuracy of imputed SVs.

### Benchmarking different panels of genetic variability

QCed genotypes generated in the GCAT cohort with the Infinium Expanded Multi-Ethnic Genotyping Array (MEGAEx) (i.e. 756,773 SNVs) were used to impute 4,448 individuals (e.g. excluding those 785 with WGS) using the GCAT|Panel and the public available 1000G phase3^11^, GoNL-SV^12^, UK10K^29^ and HRC^2^ reference panels. Multiple reference panel imputation was conducted using GUIDANCE^64^. For comparative purposes, we considered imputed variants with info imputation score ≥ 0.7 and MAF > 0.001. Variants were considered identical if they coincide at the same position and ALT allele in the case of SNVs and indels and overlap in a ±1kb window and type in the case of SV. Since allele frequency impacts imputation, we calculated the average of the info imputation score (r²) by frequency; rare (MAF < 0.01), low frequency (0.01 ≤ MAF < 0.05) and common (MAF ≥ 0.05).

### Functional impact of Structural Variants

#### Structural Variant annotation

Functional, regulatory, and clinical annotations of SVs were predicted using AnnotSV^25^. The functional impact of SVs was evaluated by considering (i) the level of overlap with known genes, (ii) the level of overlap with regulatory regions^65^, (iii) the predicted loss of function intolerance (pLI) effect and (iv) the reported disease association studies.

#### Comparison with the GWAS catalog

GWAS catalog version 1.0 (e98 r2020-03-08) was downloaded from https://www.ebi.ac.uk/gwas/docs/file-downloads. First, we selected 106,906 variant-phenotype associations of 72,849 unique autosomal entries identified in European ancestry. Second, we intersected with PLINK2.0^66^ 68,323 unique variant-phenotype associations (MAF > 0.01) with the GCAT dataset (∼30M variants) by chromosome and breakpoint. Finally, we identified 1,374 unique SVs (MAF > 0.01) in strong linkage disequilibrium (r^2^ > 0.80) with variant-phenotype associations in 1Mb window (Supplementary Fig. 27). From these 1,374 SVs, we evaluated the type of SVs and the overlapping with genes and regulatory regions.

#### Genome-Wide Association Analysis

Association analysis was performed for 70 multiples independent GWAS in selected phenotypes. Phenome analysis included only chronic conditions defined by linked Electronic Health Records from the cohort registry (2012-2017). Chronic conditions were selected according to the indicator of the chronic condition^67, 68^ and grouping the resulting conditions by similarity, considering ICD-9 codes and descriptions. Conditions with more than 50 cases were retained for the GWAS analysis (i.e. 70). Each genomic wide association tests were performed as independent logistic regression for each cohort under the assumption of an additive model for allelic effects, with adjustments made for age, sex, and the first five principal components. Gender-specific conditions were analysed only for a specific gender. Analysis was performed using PLINK2.0^66^ for autosomal chromosomes. Locus Zoom was derived for specific regions, and suggestive tower profiles were analysed, based on LD patterns and gene centred impact. Bonferroni correction was considered for 10 independent body systems.

#### Experimental validation of the Alu element

PCR amplicon analysis was designed using Primer 3.0 software using the hg19_dna range=chr3:49,492,813-49,496,062 sequence, including the Alu element. Sequence primers are for F-primer, (5’CATTGACTCATTCAGCAAGCA3’) and for R-primer (5’AAATTAAGCCCCACCCTAG3’). Using standard conditions (35X, Tm=60°C) in a Veriti™ 96-Well Thermal Cycler (Thermo Fisher Scientific), we obtain a 515 bp fragment corresponding to the control-allele and a 848 bp for the Alu- allele. Fragments were resolved by e-agarose gel, in a TapeStation (Agilent). Further, the amplicon of a non-ALU allele carrier was analysed by Sanger Sequence Method to verify the insertion point (i.e. at hg19 Chr3:49,494,280) and the ALU sequence insertion (324 bp).

### Statistical analyses

R software was used for data visualization and statistical analyses. 95% confidence intervals (CI) for recall, precision and genotype error metrics were assessed as point estimation ±1.96*s.d. Risk ratios with 95% CI and two-tailed p-values from the functional enrichment of common and rare SVs were calculated using the riskratio() function from the epitools R package. Pearson correlation coefficient with 95% CI and two-tailed p-value were estimated using the cor.test() function implemented in R.

## RESULTS

### Evaluation of cohort data quality and consistency

808 individuals were randomly selected (gender-balanced) for Illumina whole-genome sequencing (30X) from the GCAT cohort^17^. 23 samples were excluded based on sequence quality, ethnicity and relatedness parameters (see Methods, Supplementary Table 10). Principal component (PC) analysis on the remaining 785 individuals identified a unique and separated cluster compared with geographically neighbouring populations (Fig. 1a, Supplementary Fig. 7), in agreement with their geographic origin^19^.

**Fig. 1.**
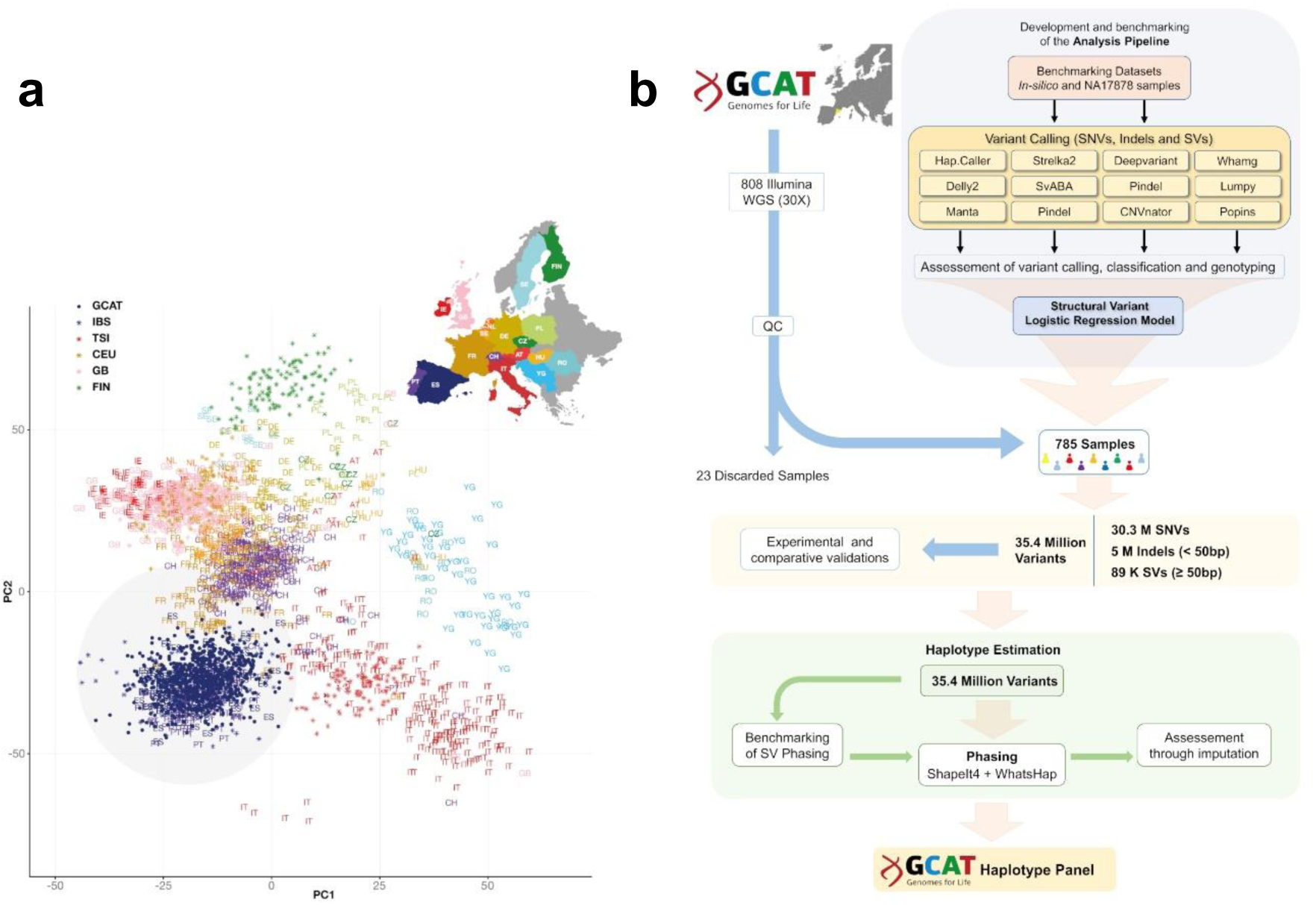
Overview of data and overall strategy. **a**, Distribution of genetic data (SNVs) based on principal component (PC) analysis (adapted from Novembre et al^19^). The PC grouped by geographic localization (coloured in grey) the individuals of the GCAT cohort (blue dots) with Iberian samples from 1000G (asterisk) and POPRES (letters) projects in the context of other European samples. **b**, Flowchart of the overall strategy followed in this study, covering from the quality control of the initial data, to the final generation of the GCAT Haplotype Panel, with particular focus on SVs. Overall, the complete strategy consumed approximately 3.5 Million CPU/hour, which highlights part of the computational challenges associated to this type of analyses (See also Supplementary Fig. 7).

### Generation of a comprehensive variant identification strategy

We designed, benchmarked and implemented a comprehensive strategy for capturing, classifying and phasing a wide range of germline variants from short-read Illumina sequences, with particular efforts devoted to the identification and subclassification of larger structural variants (Fig. 1b). Using sequencing data from an *in-silico* genome (Supplementary information 1, Supplementary Table 2), and a real sample (NA17878, from the Genome In A Bottle (GIAB) project^20^), we assessed the performance (i.e. recall, precision and F-score metrics) of 17 variant callers covering SNVs, small indels (<50bp), and large SVs (≥50bp) (see Methods), and retained the best twelve (Supplementary Table 3). SNVs were first filtered based on a minimum constraint of having the support from at least two callers, which provided high recall (>95%) and precision (>96%) values. On the other hand, for the filtering of small indels and SVs, which show high levels of discrepancy across individual callers, and their combinations (Fig. 2a), we built a Logistic Regression Model (LRM), to prioritize caller results through a reliability score from the weighted combination of different calling parameters (Fig. 2b, Supplementary Fig. 2) (see Methods). This approach outperformed other typical curation strategies over the entire spectrum of SV sizes (Fig. 2c, Supplementary Fig. 4). Furthermore, because accurate genotype calling is also key for downstream analyses, on top of this LRM, we prioritized those callers that best-resolved the heterozygosity (i.e. genotypes), resulting in a lower rate of genotype error across all variant types (<6%) (Fig. 2d, Supplementary Fig. 3) (see Methods).

**Fig. 2.**
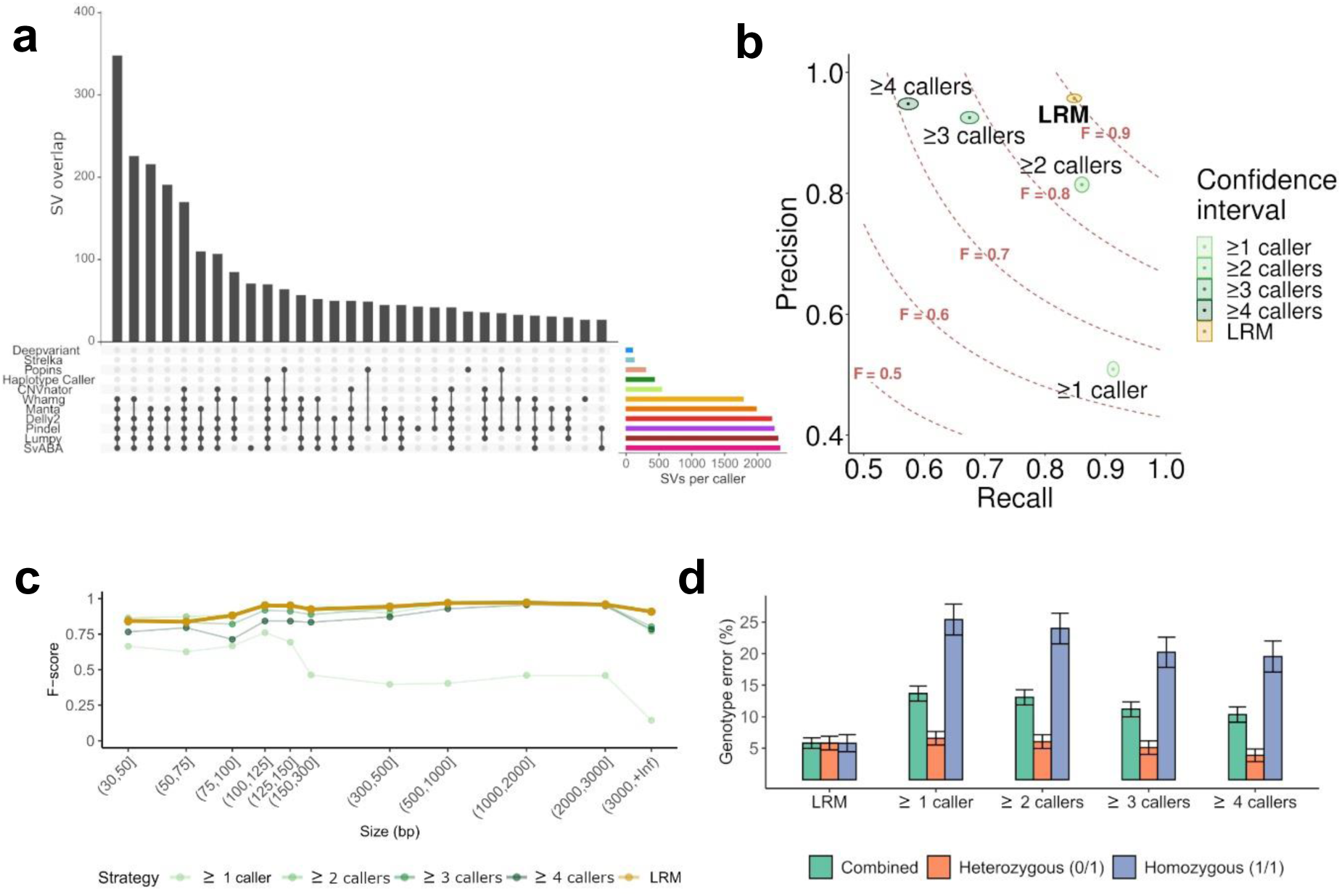
Benchmarking of the structural variant identification and classification pipeline. **a,** Structural variant (SV) detection patterns according to the programs used. Lines and dots indicate the programs used and bars the number of overlapping calls resulting from that combination. The first 30 patterns with more coincident SV calling are shown. Right coloured horizontal bars indicate the total number of SVs detected by caller. Variant callers that detect all SV types and sizes tend to recover more SVs than those that detect specific SV types (i.e. CNVnator) and smaller SVs (i.e. Strelka2). **b**, Overview of the detection performance of different strategies and filtering results from multiple variant callers. Each strategy is plotted according to the recall and precision ratios (F= F-score) using the benchmarking dataset. The Logistic Regression Model (LRM), with a F-score of 0.9, outperformed other commonly used strategies that are based on the number of coincident callers (logical rules). The confidence interval for each case is represented by coloured area of each strategy. **c**, Comparison of performances (F-score) of different merging and filtering strategies according to the size of the structural variant. **d**, Comparative overview of the genotype error, associated to each strategy for each allelic state. Error values and their intervals were inferred from the benchmarking dataset (See also Supplementary Fig. 2, Supplementary Fig. 3, Supplementary Fig. 4).

### Genome-wide variation analysis of the GCAT cohort

The application of this strategy to the selected 785 whole-genome Illumina sequences (30X), let us identify 35,431,441 unique variants across the cohort. Of these, 85.6% correspond to SNV, 14.1% to indels (<50bp) and 0.3% (n= 89,178) to SVs (≥50bp) (Fig. 3a). Median values of variants per individual were 3.52M SNVs (SD=24,983), 606,336 indels (SD=8,060) and 6,393 SVs (SD=222), showing good consistency across the cohort (Fig. 3b) and affecting a median of 7% of the entire genome. SV sizes ranged from 50 bp to 197 MB (duplication), with median values of 291 bp and a different distribution for each type of variation (Fig. 3c). Frequency ranges across all SVs were in agreement with other public WGS-based studies (Fig. 3d), with 31% of them being common or low-frequency (MAF ≥ 0.01), and 69% being rare (MAF < 0.01), including a large fraction (50%) present only in one or two individuals (i.e. MAF ≤ 0.0025).

**Fig. 3.**
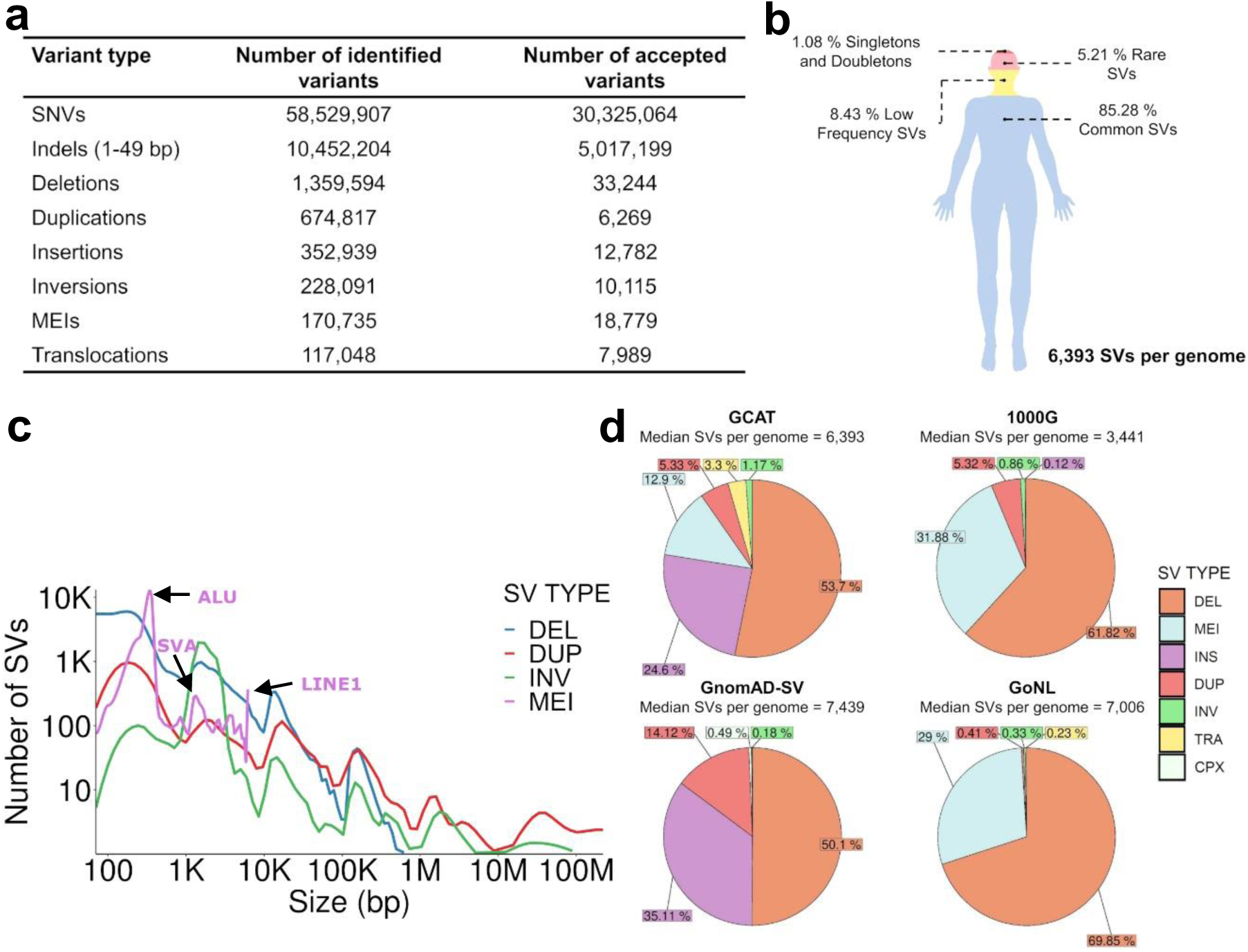
Overview of the GCAT variant catalogue. **a**, Table with the numbers of identified and accepted variants after applying the filters “at least 2 callers detecting the same variant” for SNVs, the LRM for indels and SVs, Hardy-Weinberg Equilibrium, and discard monomorfic variants and those with >10% missingness within the GCAT cohort, according to their class. **b**, Overview of the variant distribution within an average individual in the cohort, according to their observed Minor Allele Frequency (MAF). **c**, Distribution of SV type according to their genomic sizes. **d**, Comparative overview of the number and distribution of SV type across the GCAT, 1000G, GnomAD and GoNL catalogues. More inversions per genome were estimated with GCAT cohort samples.

The robustness of these results was evaluated using comparative and experimental approaches. A large fraction of SNVs and indels (i.e. >79% and >93% respectively) matched with dbSNP database^21^ (Build 153.v) entries (Supplementary Fig. 9a,b). Regarding SVs, the comparison against different public databases (i.e. gnomAD-SV^10^, 1000G^11^, GoNL^12^, HGSVC^13^, DGV^22^, dbVar^22^, Ira M.Hall Lab dataset^23^; see Methods) highlighted 49,333 novel SVs (i.e. 61% of all SVs), of which 27% were present in more than two individuals (Supplementary Fig. 9). As to the type, 26% of these novel variants correspond to deletions, 8% to duplications, 20% to insertions, 20% to inversions, 4% to LINEs, 1% to SVAs and 21% to Alu elements. Experimental validations were performed on a subset of the identified variants. The comparison of our results with array-based genotypes in a fraction of our cohort (n=570 individuals) validated 96% and 87% of SNVs and indels, respectively, with a genotype concordance of 97% and 96% (Supplementary Fig. 10). Furthermore, we also used a benchmarking set of 59 manually-curated and experimentally-genotyped inversions with MAF>0.01 from the InvFEST project^24^ to evaluate this type of variants within our catalogue. Of these 59 inversions, we detected 51 (86%), with concordant size and allele frequency values (Supplementary Fig. 11a, b; see Methods). This validates ∼38,000 of ∼40,000 independent inversion calls across the entire cohort, with an average genotype concordance of 95% (Supplementary Fig. 11c).

### Predicted functional impact of SVs

A first assessment of the potential functional impact and pathogenicity of our SVs was obtained using AnnotSV^25^. 46% of all SVs overlapped with genes, affecting a median of 2,868 per individual, whereas 18% overlapped with gene regulatory regions. While the majority (88%) of gene-overlapping SVs mapped within intronic regions (Supplementary Fig. 24a), 9% of them affected coding sequencing regions (CDS). In agreement with known variant fixation patterns within populations, we observed that rare SVs (MAF < 0.01) tend to be more disruptive, compared to common variants (MAF ≥ 0.05), as 13% of rare SVs are overlapping coding regions, compared to 5% of the common ones (RR = 0.13/0.054 = 2.4, 95% CI = [2.14,2.69], p-value = 2.6×10^-67^, Supplementary Tables 15a, 15b). Of the affected genes, 28% (10,600 SVs) are related to disease, as indicated by the predicted-loss-of-function intolerance parameter (pLI)^26^ (Supplementary Fig. 25a). In addition, 799 of the SVs overlapping genes or regulatory regions are in strong linkage disequilibrium (LD) (r^2^ ≥ 0.8) with variants associated with human traits from the GWAS Catalog.

### Iberian Haplotypes estimation

As a resource for the enrichment of SVs within genome-wide studies, we built a haplotype reference panel by phasing together all the variants identified within all GCAT samples. We first generated a cross-validation framework to identify the best available phasing strategy for SV (see Methods), using imputation as evaluation criteria (Supplementary Fig. 12, Supplementary Fig. 13). In our hands, the combination of ShapeIt4^27^ and WhatsHap^28^, which include phase informative reads (PIRs), provided the best results. With this protocol, the resulting haplotype panel allowed the imputation (info scores > 0.7) of 98%, 92%, and 90% of our common SNVs, indels and SVs, respectively, recovering a median of 5,120 SVs (SD=50), from a maximum of 6,393 SVs estimated per individual. While the best imputation results came from *de novo* insertions and deletions, with 96% and 95% recovery rates, respectively, duplications and translocations were imputed at lower rates, i.e. 48% and 19%, respectively (Supplementary Fig 15). Overall we imputed common SNVs, indels and SVs with a genotyping concordance of 99% (SD=0.4), 97% (SD=0.6) and 98% (SD=1.2) (Fig 4a), respectively. The lowest values were observed for duplications and translocations, with concordances of 84% (SD=9.2) and 73% (SD=27.6), respectively (Supplementary Fig. 16).

**Fig. 4.**
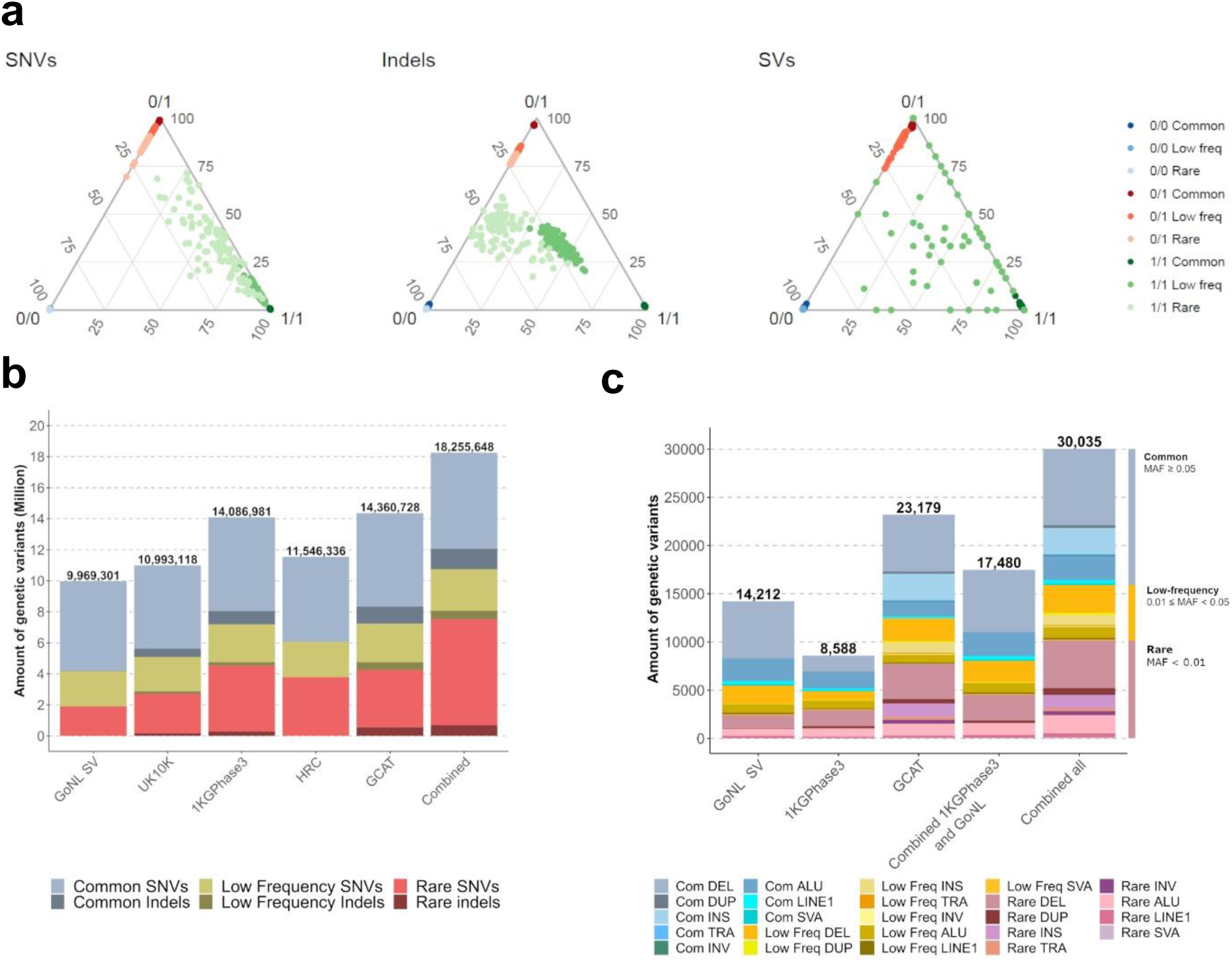
Phasing and Imputation performance of the GCAT|Panel. **a**, Ternary diagram of the genotype imputation accuracy by variant type and frequency, considering the genotype calling as reference. Three dots evaluate each genotype state per sample. The samples with high concordances between genotype imputation and genotype calling were located at ternary diagram vertices. **b**, Number of SNVs and indels imputed (info score ≥ 0.7) using different reference panels and combining their imputation results. More indels were recovered by GCAT|Panel. **c**, Number of SVs imputed (info score ≥ 0.7) using different panels, and combining the imputation results with and without GCAT|Panel. **e,** GCAT|Panel imputation results across all continents. European and Latin American populations recover more low frequency and rare variants at high infor socres (≥0.7) than African and Asian populations (See also Supplementary Fig. 15 and Supplementary Fig.16).

As the possibilities of accurately impute SVs are expected to correlate with the number of neighbouring SNVs and indels in LD, we next analysed the variation context of our SVs. Using one megabase window, we observed that the number of variants in strong LD (r^2^ ≥ 0.8) with common deletions, insertions, inversions and mobile element insertions (MEIs) was in the range of 39-42, in contrast to duplications and translocations, which showed mean values of 12 and 8 variants respectively (Supplementary Fig 17a). In fact, a positive significant correlation was observed between the number of variants in LD and the score of imputation for common SVs (Pearson’s r = 0.38, 95% CI = [0.37,0.40], p-value < 2×10^-16^) (Supplementary Fig. 17b), and for all SV types (except translocations) (Supplementary Fig. 18).

### Imputation performance of the haplotype panel

Following this strategy, we generated a complete and operational panel of Iberian haplotypes, with all the variants of our 785 individuals. The resulting GCAT|Panel was assessed through the imputation of the genotyping array data of 4,448 GCAT individuals, and compared the results with those of other reference panels, such as 1000G^11^, GoNL^12^, HRC^2^ and UK10K^29^. With IMPUTE2^30^, the GCAT|Panel was able to impute a total of 14,383,907 variants with MAF > 0.001 and high quality (info score ≥ 0.7). Across different reference panels, the overall imputation performance for SNVs and indels (<50 bp) was generally high (Fig 4b), with slight overperformances of the GCAT|Panel on indels, and of 1000G and HRC panels on SNVs. While HRC and 1000G recovered rarer SNVs, likely because of their larger sample sizes, the GCAT|Panel was able to recover rarer indels (Fig. 4b). At the structural variation level, the GCAT|Panel was able to impute a total of 23,179 SVs with info scores ≥ 0.7, resulting in a 1.6, 2.7 and 1.3-fold increase, compared with the 1000G, the GoNL and both panels combined, respectively (Fig 4c). For common SNVs/Indels (MAF > 0.05) the GCAT|Panel showed similar performance as HRC, 1000G, GoNL and UK10K reference panels (mean r^2^ > 0.96, Supplementary Fig. 13a). For common SVs, the GCAT|Panel outperformed (mean r^2^ = 0.91, SD=0.15) 1000G (mean r^2^= 0.80, SD=0.21) and GoNL-SV reference panels (mean r^2^ = 0.82, SD=0.21, Kruskal-Wallis p-value < 2.2×10^-16^, Supplementary Fig. 21b).

Informative structural variants imputed (info score ≥ 0.7) by the GCAT|Panel were tested together with SNV/indels for association across 70 chronic conditions grouped in 10 independent body systems (n cases ≥ 50) from the GCAT cohort (n = 4,988). 46 SV loci showed suggestive association (p-value ≤ 1×10^-6^ corrected by 10 body systems) with 26 conditions (Supplementary Fig. 28). Of all these associations, 63% could potentially be functionally explained through SVs, as they either lead the association (37%), or are in strong LD (r² ≥0.8) with the lead variant (26%). A notable example is a rare AluYa5-element in chr3 (g.49494276_49494600ins (hs37d5), MAF = 0.0013), located near the dystroglycan gene (*DAG1*) and associated (p-value = 9.84×10^-7^) with mononeuritis of lower limb (ICD-9 355) (Fig. 5a). This Alu element, imputed only with the GCAT|Panel (info score = 0.98), was experimentally confirmed in all carrier individuals (Fig. 5b, Supplementary Fig. 29).

**Fig. 5.**
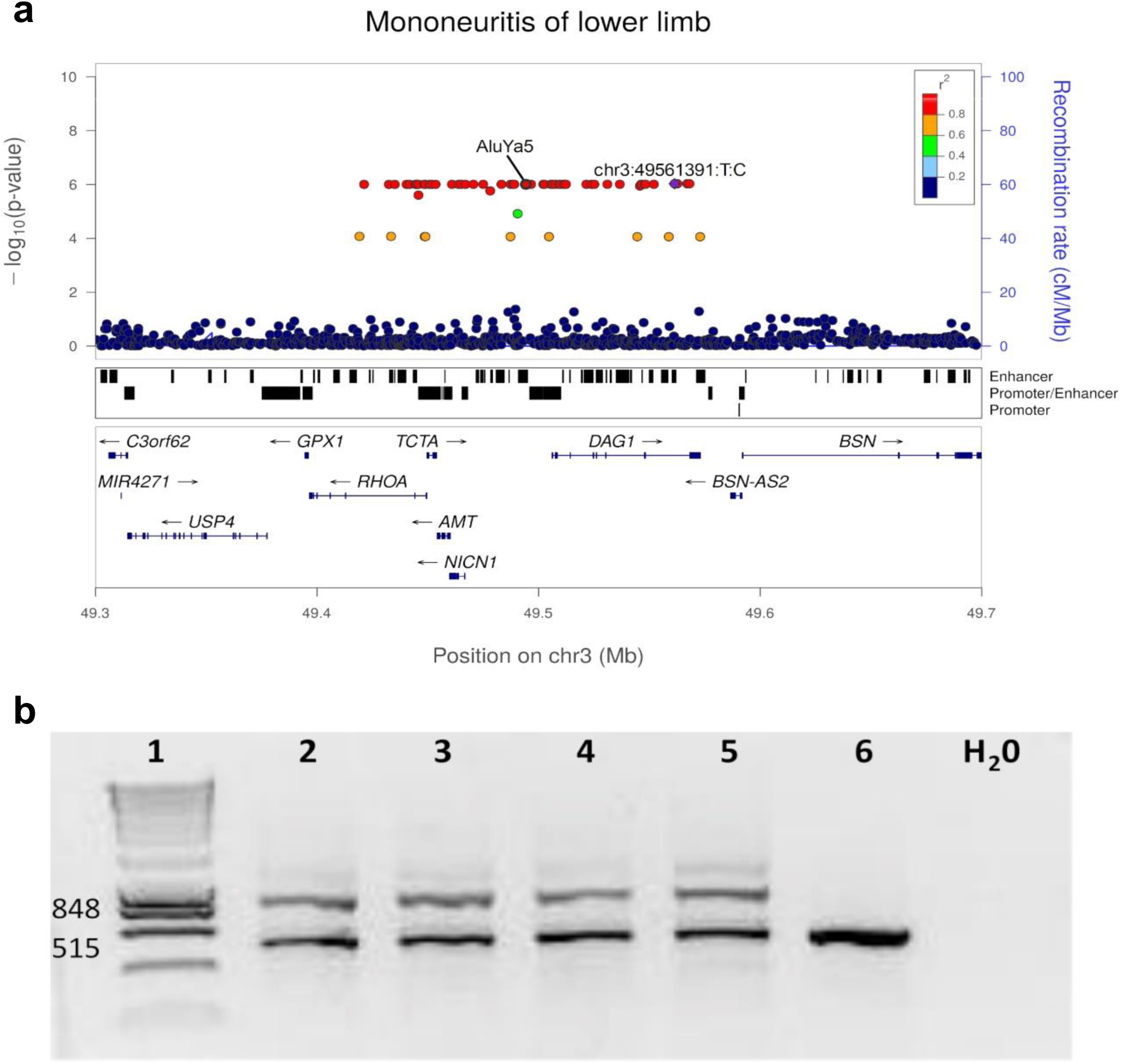
Genome-Wide Association Analysis using GCAT|Panel and Experimental validation of an AluYa5-element. **a,** Locus Zoom plot of the locus associated with mononeuritis of lower limb (ICD-9 355) (p-value = 9.84 x10^-7^), showing the lead variant in purpule. The AluYa5-element (g.49494276_49494600ins (hs37d5) maps in an enhancer element upstream of the *DAG1*. **b,** Experimental validation of an AluYa5- element, agarose e-gel electrophoresis of PCR products after amplification of Alu- insertion-specific DNA fragments from blood DNA Lanes: 1, 100 bp DNA ladder marker (Life Technologies), expected sizes of both states are shown to the left; 2-5 Alu carriers (EGA_04200, EGA_01901, EGA_13378, EGA_03940); 6 control individual (EGA_01399). The numbers to the left refer to the size (bp) of marker DNA fragments. Electrophoresis analysis of Alu carriers show two-band amplicons (515 bp and 848 bp) detected in Alu carriers (Lanes 2-5) and one-band amplicon (515 bp) in control non-Alu-allele individuals (Lane 6)(See also Supplementary Fig. 29).

Finally, we evaluated the portability of the GCAT|Panel to infer SVs across 19 different ethnic groups using 1,880 individuals from the 1000G project. While the imputation quality of SVs was higher within the European populations, the GCAT|Panel was also able to impute a large fraction of SVs across all other ethnicities (Fig 6, Supplementary Fig. 23a). Of nearly 50K unique SVs imputed across all groups, 25%, 35% and 40% of them were detected within the Asian, African and Latin American populations, respectively (Fig 6, Supplementary Fig. 23). In agreement with the mixed origin of Latin Americans, nearly half of all imputed variants within this group showed low-frequency values (MAF < 0.05), compared with other non-European groups, where the imputation covered predominantly common variants (Fig 6). In addition, 73% of all the structural variants identified and genotyped in previous studies, using long and short WGS^15, 32^ were also imputed by our panel on the same individuals, with 88% of matching genotypes (Supplementary Fig. 19a and Supplementary Fig. 20a).

**Fig. 6.**
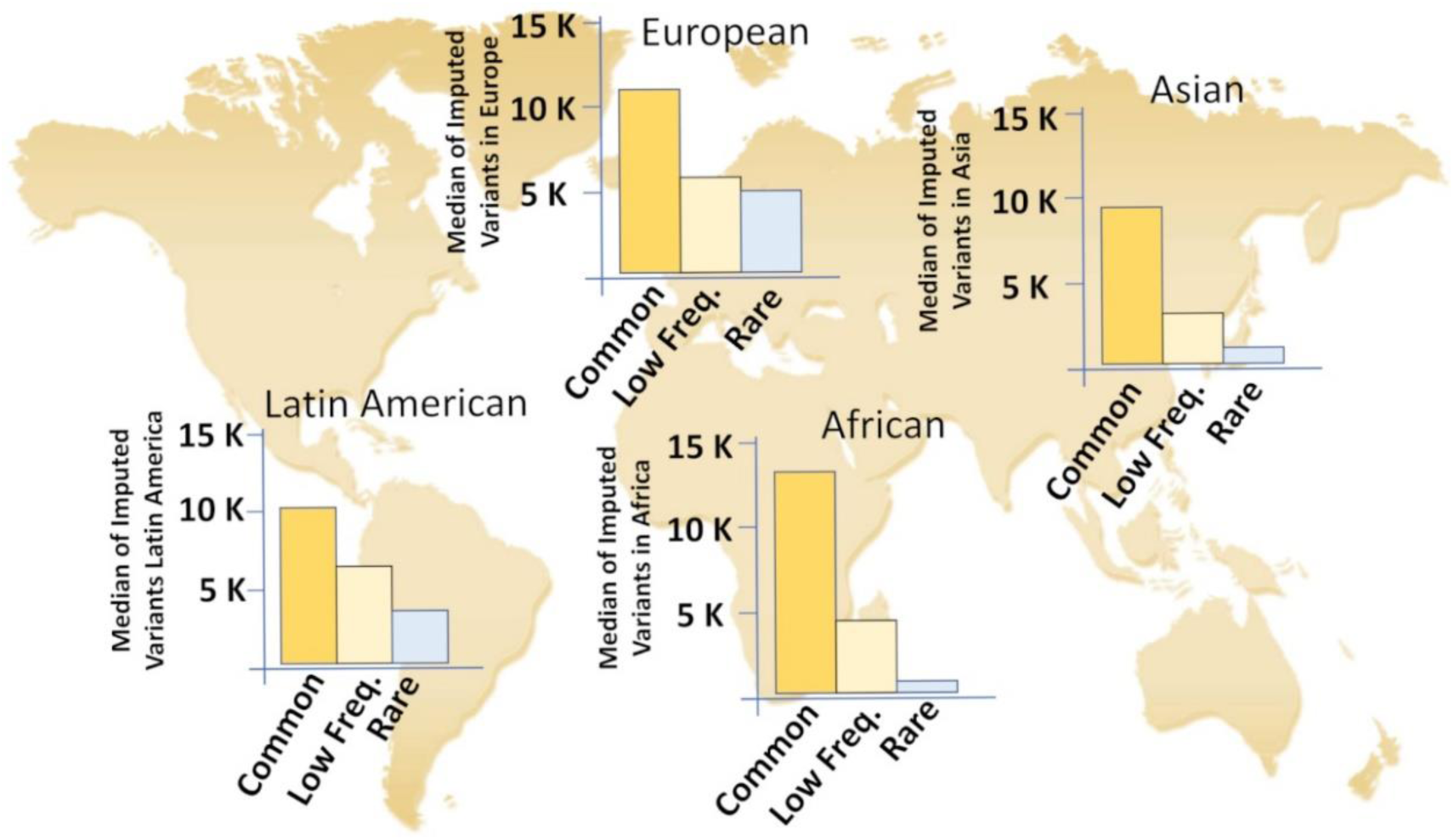
Structural Variant imputation performance using GCAT|Panel across all continents. European and Latin American populations recover more low frequency and rare SVs at high infor socres (≥0.7) than African and Asian populations (See also Supplementary Fig 23).

## DISCUSSION

Here we present the GCAT|Panel, the first Iberian Haplotype reference panel derived from high-coverage whole-genome sequencing. The strategy developed for variant identification, classification, and phasing, has provided a comprehensive and high-quality catalogue of genetic variants, with low rates of false-positive calls and genotyping errors for all variant types, including SVs. This is due to the combination of high sequencing coverage (30X) with a comprehensive analysis strategy that integrates multiple variant callers and a Logistic Regression Model for maximizing recall and precision for each SV type and size.

Increasing the sequencing coverage to 30X allowed us to resolve a large fraction of SVs and accurately define the genotypes that cannot be properly defined with lower sequencing depths. In addition, while previous projects inferred SVs into phased haplotype scaffolds^11, 12^, our sequencing coverage allows us, for the first time, to phase SVs together with biallelic SNVs and indels, and to use phase informative reads (PIRs), which are expected to improve the imputation of rare variants^31^. With this sequencing technology, we also expect a slight detection bias in our study against low complexity regions of the genome, where short-read sequencing tends to be less informative, in contrast to long-read sequencing technology^13-16^. This is further highlighted by the high portion (54%) of our SVs affecting genes or regulatory regions, which also tend to be within the non-repetitive portion of the genome.

Given the increasing incorporation of whole-genome sequencing into genetic studies, it is crucial to highlight the importance of accurately identifying and resolving SVs with the correct genotype, to then obtain robust and meaningful results during the imputation in a different cohort. Here, we found a positive correlation between the number of neighbouring variants in LD with SVs and their quality of imputation, suggesting that variants with a high genotyping error show a lower number of variants in LD, which translates into a lower imputation accuracy for those variants (Supplementary Fig. 17). On the other hand, software limitations (PLINK or ShapeIt4), can translate into poor estimations of haplotypes and LD, directly hampering the association test, which relies on accurate counts of variant allele frequencies and states. Improved variant calling strategies that can accurately identify and define complex structural variation events are still needed, together with new and dedicated analysis frames (e.g. phasing and LD) for SVs, where the actual size and type of the variant is considered, in contrast to the current scenario where SVs are taken as SNVs.

In our cohort, the GCAT|Panel led to the identification of potential risk SV, including those within the rare spectrum. Here, we highlight the identification of a rare polymorphic 324 bp-long AluYa5 element in chromosome 3 (g.49494276_49494600, MAF = 0.0013) associated with mononeuritis of the lower limb (ICD-9 355). This SV is located within a multi enhancer-elite element (GeneCards)^33^, proximal to *DAG1*, a gene involved in pathways responsible for neuromuscular diseases, and already associated with severe limb-girdle muscular dystrophy type 2P (LGMD2P) through missense point mutations^34^. Further studies are now needed to validate the resulting hypothesis, in which this Alu element could be affecting the expression of the *DAG1* gene in this disease.

Finally, this study also provides detailed guidance for the comprehensive analysis of whole-genome sequences, including the identification, classification, and phasing of SVs. We expect that this type of analysis will soon become the standard within large genetics studies that are already incorporating whole-genome Illumina sequences and combining them with existing genotyping array information. In agreement with other studies^35^, our analysis evidences the potential of using population-matched reference panels and their clear benefit, in particular, for the identification of rare variants and to complete our understanding of the underlying genomic architecture of genetic diseases.

## RESOURCE AVAILABILITY

Below we attatch the information of the data and code availability used in this study.

### Contact for resource sharing

Further information and requests for resources should be directed to and will be fulfilled by the Lead Contact, David Torrents.

### Data and code availability

808 FASTQ and BAM files data, the VCF files with genotype information, GCAT- genotyping array, and GCAT|Panel are deposited at European Genome-phenome Archive (EGA) under the accession number EGAS00001003018. The EGAS00001003018 supporting the current study have not been deposited in a public repository because it includes genetic information of volunteers but are available from the corresponding author on request. The variants of the GCAT catalogue accepted after variant filtering, the SV annotation files (both resources under petition) (Fig. 3a) and the *in-silico* information (FASTQ, BAM files, catalogue of variants inserted) are available in http://cg.bsc.es/GCAT_BSC_iberianpanel.

All original code has been deposited at (https://github.com/gcatbiobank/GCAT_panel) and is publicly available as of the date of publication. DOIs are listed in the key resources table.

## Supplementary information

Refer to Web Version on PubMed Central for supplementary material

### FUNDING AND ACKNOWLEDGEMENTS

This study makes use of data generated by the GCAT|Genomes for Life, a cohort study of the Genomes of Catalonia, Fundacio Institut Germans Trias i Pujol (IGTP). IGTP is part of the CERCA Program / Generalitat de Catalunya. GCAT is supported by Acción de Dinamización del ISCIII-MINECO and the Ministry of Health of the Generalitat of Catalunya (ADE 10/00026); the Agència de Gestió d’Ajuts Universitaris i de Recerca (AGAUR) (2017-SGR 529). B.C is supported by national grants PI18/01512. X.F is supported by VEIS project (001- P-001647) (co-funded by European Regional Development Fund (ERDF), “A way to build Europe”). Full list of the investigators who contributed to the generation of the GCAT data is available from www.genomesforlife.com/. We thank Dr. Lluís Puig and Vanessa Plegezuelos on behalf of Blood and Tissue Bank from Catalonia (BST), who collaborated in GCAT recruitment, and all the GCAT volunteers that participate in the study. We also thank the Centro Nacional de Análisis Genómico (CNAG-CRG), who collaborated in the sequence analysis of the study, the other members of the Comparative and Functional Genomics Group at the UAB for help with inversion validatioņ Dr. Francesc Calafell for his comments on the manuscript, Marta Morell from qGenomics for the technical assistance in Alu validation, the collaboration of all the Computational Genomics Group at the BSC, specially Ignasi Morán and Lorena Alonso, for their helpful discussions and valuable comments on the manuscript, and the technical support group from the Barcelona Supercomputing Center Center is gratefully acknowledged. This work has been sponsored by the SEV-2011-00067 and SEV2015-0493 grants of the Severo Ochoa Program, awarded by the Spanish Government, by the grant TIN2015- 65316-P, awarded by the Spanish Ministry of Science and Innovation and by the Generalitat de Catalunya (contract 2014-SGR-1051) to D.T, and research grants BFU2016- 77244-R and PID2019-107836RB-I00 funded by the Agencia Estatal de Investigación (AEI, Spain) and the European Regional Development Fund (FEDER, EU) to M.C. J.V.M was supported by FPI (BES-2016-0077344) grant from the Spanish Ministry of Science and Innovation. C.S received funding from the European Union’s Horizon 2020 research and innovation program under the Marie Skłodowska-Curie grant agreement H2020-MSCA-COFUND-2016-754433. This study made use of data generated by the UK10K Consortium from UK10K COHORT IMPUTATION (EGAS00001000713), by formal agreement with the Barcelona Supercomputing Center (BSC). Funding for UK10K was provided by the Wellcome Trust under award WT091310. This study made use of data generated by the Genome of the Netherlands’ project, which is funded by the Netherlands Organization for Scientific Research (grant no. 184021007), allowing us to use the GoNL reference panel containing SVs, upon request (GoNL Data Access request 2019203). This study also used data generated by The Haplotype Reference Consortium (HRC) accessed through The European Genome-phenome Archive with the accession numbers EGAD00001002729, by formal agreement of the Barcelona Supercomputing Center (BSC) with WTSI. This study made use of data generated by the 1000 Genomes (1000G), accessed through the FTP portal (http://ftp.1000genomes.ebi.ac.uk/vol1/ftp/release/20130502/). This study used the GeneHancer-for-AnnotSV dump for GeneCards Suite Version 4.14, by formal agreement by the BSC and The Weizmann Institute of Science. We acknowledge Red Española de Supercomputación (RES) for awarding us access to MareNostrum4 supercomputer from Barcelona Supercomputing Center (proposal numbers BCV-2018-3-0010 and BCV-2019-1-0015).

## AUTHOR CONTRIBUTIONS

J.V.M, I.G.F, D.M.S, D.T and R.dC designed and planned the whole study. J.V.M, I.G.F, D.M.S, D.T and R.dC contributed to the writing and editing of the manuscript. A.C, prepared and QCed samples for NGS. J.V.M created the *in-silico,* and D.M.S prepared the NA12878 sample. J.V.M, I.G.F and D.M.S performed the variant calling and designed the Logistic Regression Models. J.V.M, I.G.F, D.M.S contributed to the creation of BAM files, quality control, variant merging, filtering and genotyping, with the collaboration of M.P. J.V.M, I.G.F and D.M.S performed a comprehensive comparative analysis of the Iberian catalogue with other repositories. J.V.M, I.G.F and D.M.S conducted the SNV, indel, and large deletion and duplication validations, collaborating with L.A on behalf of qGenomics conducting CGH validation analysis. The inversion validation and genotype data analysis was provided by J.L.J, M.Pu and M.C. J.V.M. I.G.F, and D.M.S performed the SV annotation, the creation of the GCAT|Panel and their benchmarking, in collaboration with C.S. R.A executed and adapted GUIDANCE with ShapeIt4 and GCAT|Panel. I.G.F, N.B, X.F, B.C conducted and analyzed the Phenome analysis, from phenotype extraction to GWAS analysis. N.B, R.dC, X.F and L.A conducted the AluY5a validation. L.S, JF.S.H conducted SNP-Array validation analysis of SV. V.M and M.Pe contributed to the editing of the manuscript. D.T and R.dC supervised the study. All authors reviewed and approved the manuscript. J.V.M, I.G.F and D.M.S contributed equally to this study.

## DECLARATION OF INTERESTS

The authors declare no competing interests

## Notes

### Competing Interest Statement

The authors have declared no competing interest.

