## Supplementary Information for "GCAT|Panel, a comprehensive structural variant haplotype map of the Iberian population from high-coverage whole-genome sequencing"

4.2.2.3 Genome Strip (version 2.0) 13

**5 Increasing accuracy detection using a machine learning algorithm 14**

**5.1 Logistic Regression Model for Indels 14**

5.1.1 Train and Test the model 14

5.1.2 Genotype reported by LRM for SNVs and small indels 14

**5.2 Logistic Regression Model for SVs 14**

5.2.1 Train and Test the LRM for SVs 14

5.2.2 Strategy to report the position and length of variants 15

5.2.3 Genotype reported by LRM for SVs 15

5.2.4 Filtering out the SVs variants following the GoNL strategy 16

**6 GCAT project 17**

**6.1 Genomic data features 17**

**6.2 FASTQ files and BAM file generation 17**

**6.3 Quality control (QC) 17**

6.3.1 Alignment quality 18

6.3.2 Contamination analysis 18

6.3.3 Population structure using reference ancestries 18

6.3.5 Population structure without reference ancestries 19

**7 Variant calling 19**

**7.1 SNVs and Indels calling 19**

7.1.1 Haplotype Caller 19

7.1.2 Strelka2 19

7.1.3 Deepvariant 19

**7.2 Mid-size and Structural Variant calling 20**

7.2.1 Delly2 20

7.2.2 Manta 20

7.2.3 Pindel 20

7.2.4 Lumpy 20

7.2.5 SvABA 20

7.2.6 Whamg 20

7.2.7 CNVnator 21

7.2.8 Popins 21

7.2.9 Melt 21

**8 Variant Calling integration 21**

**8.1 VCF pre-processing 21**

8.1.1 SNV and indels set 21

8.1.2 Structural Variants set 21

**8.2 Merging all VCFs per-sample and variant type ... 22**

**8.3 Combining all samples in one VCF ... 22**

**8.4 Variant Quality Control ... 22**

### 9 Validation of variants ... 23

**9.1 Public datasets comparative ... 23**

9.1.1 SNVs and Indels ... 23

9.1.2 Structural Variants ... 23

**9.2 Experimental validation ... 23**

9.2.1 SNV and Indel validation using SNP-array genotyping ... 23

9.2.2 Inversions validation using a verified dataset ... 24

### 10 Creation and integration set of haplotypes ... 24

**10.1 Benchmarking of different phasing strategies ... 24**

10.1.1 Sample pre-processing ... 24

10.1.2 Phasing strategies ... 25

10.1.2.1 ShapeIt2 (with SVs) ... 25

10.1.2.2 ShapeIt2 and MVNcall ... 25

10.1.2.3 ShapeIt2, MVNcall with PIRs ... 25

10.1.2.4 ShapeIt4 ... 26

10.1.2.5 ShapeIt4 and MVNcall ... 26

10.1.2.6 ShapeIt4 and WhatsHap ... 26

10.1.3 Imputation using different phasing strategies ... 26

10.1.4 Imputation accuracy ... 27

**10.2 Pipeline to construct the GCAT haplotype panel ... 27**

### 11 Imputation using the GCAT reference panel ... 27

**11.1 Imputation analysis using the GCAT SNP-array ... 27**

**11.2 Imputation quality using the array of 1000G ... 28**

11.2.1 Sample filtering and Quality Control of the 1000G array ... 28

11.2.2 Imputation study of GCAT panel with non-Iberian samples ... 28

**11.3 Preliminary discoveries using the GCAT|Panel ... 29**

11.3.1 Genome-Wide Association Analysis ... 29

11.3.2 Experimental validation of the Alu element ... 30

### 12 Benchmarking different panels of genetic variability ... 30

### 13 Biological impact of Structural variants ... 30

**13.1 Structural Variant distribution in worldwide populations ... 30**

**13.2 Functional implications of Structural Variant in humans ... 31**

13.2.1 Structural variant annotation using AnnotSV ... 31

### 1 *In-silico* sample

An artificial sample (hereafter *in-silico*) was created by ART software (ART-Illumina) to calibrate and benchmark all variant callers used in this project. The *in-silico* FASTQ files were derived from Illumina's technology, covering the wide spectrum of genome rearrangements (Supplementary Table 1).

In order to generate a sample mimicking the real human genome variation, we inserted in the reference genome (hg19) sequence 5,330,762 human variants described in the 1000 Genomes (1000G) and the PanCancer project. Additionally, we generated and inserted using scripts in-house 3,925 Structural Variants (artificial SVs). In brief, the *in-silico* includes 4,871,660 Single Nucleotide Variants (SNVs), 723 Point Mutations, 458,047 indels, 2,181 Deletions (DEL), 901 Insertions (INS), 386 Inversions (INV), 772 Duplications (DUP), 62 Copy Number Variants (CNVs), 755 Translocations (TRA), 100 Transposons (TRP), 89 Viruses (VIR) and 9 Pseudogenes (PSG). Supplementary Table2 summarises the number and origin of all variants (The list of *in-silico* variants is deposited at http://cg.bsc.es/GCAT_BSC_iberianpanel/).

Selected variants were inserted in FASTA files of hg19 genome by chromosome (http://hgdownload.cse.ucsc.edu/goldenpath/hg19/chromosomes/). For each chromosome, we generated two haplotypes with their respective variants as follows. Variants were randomly distributed for each artificial chromosome in the whole sequence, excluding telomere and centromere regions. Initial heterozygotes were used to generate homozygotes genotypes by duplicating these variants (flag "Homozigotic"). We used ART (version 2.5.8) to simulate FASTQ files sequencing data, then we derived the BAM file using the reference genome hs37d5 (decoy version) and BWA (version 0.7.15-r1140) and Samtools (version 1.5). This method creates synthetic NGS sequencing reads. The command used to run ART-Illumina is the following:

```
art_illumina -ss HS20 -i input.fa -rs 4 -qs 1 -qs2 1 -d _idallele_ -p   -l
                   100 -f 15 --mflen 500 --sdev 20 -o out
```

In order to correct (i) technical biases, (ii) make the data suitable for analysis and (iii) reduce the false positives in the variant calling step, we applied the Best Practices of GATK. We marked the duplicated reads using PICARD (version 1.108) and we recalibrated the Base Quality Scores (BQSR) of the BAM file using two modules (VariantRecalibrator, ApplyVQSR) of GATK4 package (version 4.0.11) (*in-silico* FASTQ and BAM files are deposited at http://cg.bsc.es/GCAT_BSC_iberianpanel/).

#### 2 Genome in a Bottle sample

To validate the calling of SNVs and indels, we used the sample NA1287831 from the genome in a Bottle (GIAB) Consortium as follows. Firstly, we downloaded the down-sampling 30X BAM file of NA12878 (RMNISTHS_30xdownsample.bam) from ftp://ftp-trace.ncbi.nlm.nih.gov/giab/ftp/ data/NA12878/NIST_NA12878_ HG001_HiSeq_300x/. Secondly, we converted the BAM file to FASTQ file using Biobambam2 version 2-2.0-65, and we reconstructed it using the reference genome hs37d5 following the GATK Best Practices.

The file "HG001_GRCh37_GIAB_highconf_CG-IllFB-IllGATKHC-Ion-10X-SOLID_CHROM1X_ v.3.3.2_hi_ghconf_PGandRTGphasetransfer.vcf.gz" of NA12878 sample, downloaded from ftp://ftp-trace.ncbi.nlm.nih.gov/giab/ftp/release/NA12878_HG001/latest/GRCh37/,which contained all variants validated by the GIAB consortium, was used as a gold standard.

#### 3 Variant callers selected

Selected variant callers for the benchmarking using the *in-silico* are summarised in this section.  To better understanding, callers are presented separately into SNPs/Indels and SVs (An overview of variant callers described below can be found in Supplementary Fig. 8).

Variant Callers were executed in the Marenostrum 4 supercomputer machine at Barcelona Supercomputing Center (BSC), Barcelona. The general-purpose block of this machine consists of 48 racks housing 3,456 nodes with a total of 165,888 processor cores and 390 Terabytes of main memory. Each compute node is equipped with 48 cores (96 GB of main memory 1,880 GB/core, 12x 8GB 2667Mhz DIMM (216 special nodes with high memory, 10,368 cores with 7,928 GB/core)). The processors support well-known vectorisation instructions such as SSE, AVX up to AVX-512.

##### 3.1 Single Point and Indels Variants detection

###### 3.1.1 Haplotype Caller (GATK4 version 4.0.2.0)

Haplotype caller (HC) detects SNVs, indels (1 to 30 bp), mid-size deletions (31 to 150 bp), and INS. The strategies used to identify the variants are Split Read (SR) and Assembly Based (AS). We use java version 8u131 to run HC following three modules: 1) Run the Haplotype caller in mode gvcf; 2) Run the GenotypeGVCF module and 3) apply a variant filtration with VariantFiltration module. For each step, we used the default parameters and 16 CPUs.

###### 3.1.2 Deepvariant (version 0.6.1)

Deepvariant detects SNVs, indels, and mid-size deletions. This program uses a Machine Learning (ML) algorithm based on a deep neural network. To run this tool, we executed three steps by default parameters: 1) Run make_examples.zip, 2) Run call_variants.zip, and 3) Run the postprocess_variants.zip. To improve performance, we parallelised the first step as recommended by the developers, using 48 cores by the node.

###### 3.1.3 Strelka2 (version 2.9.2)

Strelka2 detects SNVs, indels, mid-size deletions, and INS. This caller uses the de novo assembly (AS) strategy. This tool requires two main steps: 1) Run the configureStrelkaGermlineWorkflow.py by default parameters, and 2) Run the runWorkflow.py with –m local flag. Strelka2 required 48 cores to obtain all memory of the node for each execution.

###### 3.1.4 Platypus (version 0.8.1)

Platypus detects SNVs, indels, and DELs up to 300 bp. This tool uses the strategy of local de novo assembly (AS). Before running Platypus, we imported Bcftools (version 1.6), python (version 2.7.13) and htslib (version 1.5). Then, we ran it by default parameters, including the following flags: --assemble=1 --assembleBrokenPairs=1 –mergeClustered Variants=1 –nCPU 16. These flags allowed DEL detection with lengths between 50bp to 2kb and INS between 50-500 bp. We used 16 CPUs to improve performance by parallelisation.

###### 3.1.5 VarScan2 (version 2.4.3)

VarScan2 detects SNPs and indels. This tool uses the SR strategy plus the information of map quality, coverage, and base quality. We ran Varscan2 in germline mode, as follows. 1) We executed the mpileup module from Samtools (version 1.5) by default, including the following parameters --no-BAQ --min-MQ 1, 2) with the file .mpileup, we ran VarScan2, using the mode mpileup2snp and mpileup2indel to detect SNVs and Indels respectively. Besides, we used the following flags (--min-coverage 10 --min-var-freq 0.20 --p-value 0.05) to depurate the harmful quality variants detected. For memory requirements, we used 16 CPUs.

##### 3.2 **Structural Variant (SVs)**

###### 3.2.1 Delly2 (version 0.7.7)

Delly2 detects DEL, DUP, INV, INS, and TRA. The recent version combines split-read (SR), Discordant-pairs (DP) and Read-depth (RD) strategies. This tool is executed by default parameters. We modified the –t parameter to DEL, DUP, INV, INS, and TRA, depending on the variant type to analyse. We excluded the telomere and centromere regions with –x flag because they were prone to provide false positives. We executed the steps busing 24 CPUs.

##### 3.2.2 Manta (version 1.2)

Manta detects DEL, DUP, INV, INS, and TRA. This tool combines the information of SR, DP, and de novo assembly (AS) strategies. To run Manta, we followed two steps: 1) We ran the configManta.py, which scans the genome to find SV associated regions. 2) We ran the runWorkflow.py obtained from the previous step. We executed the steps by default parameters using 24 CPUs.

##### 3.2.3 Pindel (version 0.2.5b9)

Pindel detects DEL, DUP, INV, INS, and TRA, following the SR and DP strategies. Some parameters were modified to improve SV detection. These include accuracy of the calling, detection of interchromosomal events and long insertion. Pindel was ran by chromosome using the next flags: (-a 3 -C -k -l -I -M 8 -T 6 -x 5 -v 10 -c 1 -R hs37d5 -d Feb2009). In the config file required, we included the insert size of 300 bp. We converted the BCF to VCF, using the pindel2vcf module by default parameters. Pindel was run using 8 CPUs.

We developed a custom script to genotype the Long Insertions. The genotyping was called using the reads from the BAM file, covering Pindel reported breakpoint. The reads with mapping quality $\geq 20$ were selected in a window of 10 bp from the breakpoint. Genotype was determined based on the fraction of the number of altered reads divided by the total coverage in the window. If this fraction is $\leq 0.20$, we genotyped 0/0. If the fraction was between 0.20 and 0.80, we genotyped 0/1, and if the fraction was $\geq 0.80$, we genotyped 1/1. (the script to genotype de novo insertions is deposited at https://github.com/gcatbiobank/GCATpanel/tree/main/genotyping/Pindel_genotype_insertions)

##### 3.2.4 Lumpy (version 0.2.13)

Lumpy detects DEL, DUP, INV, and BND (break-end orientation). This variant caller uses SR, DR, and a generic module (like RD). Before running Lumpy, we pre-processed the *in-silico* BAM file, following the recommendations of Lumpy developers. We extracted the split reads from the BAM file with extractSplitReads_BwaMem module provided by Lumpy and discordant reads with Samtools (version 1.5). Next, the BAMs were sorted using Samtools. After the pre-processing stage, we used Lumpyexpress by default parameters, including the –P label. Finally, we used SVTyper to genotype. We ran it as developers recommendations. After the genotyping step, we filtered out those variants which the Quality is $\leq 20$. Also, we discarded variants with SVTYPE = BND if the two chromosomes reported by the variant caller for that variant were the same. We used 12 CPUs to extract discordant and split reads and 24 CPUs and one CPU for Lumpyexpress and SVTyper tools, respectively.

###### 3.2.5 Wham (version v1.7.0-311-g4e8c)

Wham detects DEL, DUP, INS, INV. This caller uses the SR and DR strategies for variant detection and a Machine Learning (ML) algorithm to classify the SV by type. We used the Whamg version as recommended by the developers. The Whamg is executed by default parameters. Finally, we used SVType to genotype. To run Whamg, we parallelised the execution by 48 CPUs.

###### 3.2.6 SvABA (version 7.0.2)

SvABA classifies all SV types by breakpoint orientation. This variant caller uses the SR, DP and de Novo Assembly (AS) strategies. We ran SvABA in germline mode, as recommended by the developers. We did not consider the SV type reported by SvABA. To run SvABA, we parallelised the execution by 16 CPUs.

###### 3.2.7 CNVnator (version v0.3.3)

CNVnator classifies Copy Number Variations (CNV) such as DEL and DUP. This caller uses the RD strategy and detects CNVs larger than 200 bp. We ran the CNVnator following all the steps recommended by the developers, using a bin size of 100, which is the read length of the sequencing. Finally, we converted the .root file to VCF using a Perl script "cnvnator2VCF.pl" from the CNVnator toolkit. To run CNVnator, we used 12 CPUs.

###### 3.2.8 Popins (version damp_v1-151-g4010f61)

Popins detects de novo insertions (INS). This tool uses the AS strategy and detects INS ≥ 100 bp. To run this variant caller, we followed different steps: 1) assembly, 2) merge (skipped), 3) contigmap (generate the supercontigs file, 4) place-refalign, 5) place-splitalign, 6) place-finish and 7) genotype. All steps were done using the default parameters.

The NONANCHOR variants were discarded as recommended by the Popins developers. In all steps, we used 48 CPUs for each execution and to parallelise steps 1 and 3 by 48.

###### 3.2.9 MELT (version 2.1.4)

MELT detects Mobile Element Insertions (MEIs) such as Transposons (TRP). The strategy applied to identify the TRPs were SR and DP. This variant caller is executed in a SINGLE mode. The following flags are used to run MELT (–bamfile –c –e -t –h –r –k –w). We executed MELT for each transposon type: ALU, LINE1, and SVA. To run MELT, we used 24 special CPUs with high memory.

##### 3.2.10 Genome Strip (Version 2.0)

Genome Strip is designed to detect DEL, DUP, and multiple Copy Number Variants (mCNV) in cohorts. This tool combines the SR, DR and RD strategies. This variant caller requires different tools and a specific computational environment: a) Java version 1.7, b) R tool (version 3 or newer), c) Samtools and Htslib, d) and the LSF environment. We followed three steps to run the Genome Strip: 1) SVPreprocess, 2) SVDiscovery, and 3) SVGenotyper. Each step was executed by default parameters, as developers recommended, using 48 CPUs.

##### 3.2.11 Pamir (version 1.2.2)

Pamir detects INS. This tool combines the SP, DR, AS, and One End Anchored (OEA) strategies. We ran Pamir in default mode including the flag –p, to report the name of the sample which we process. Finally, the output is genotyped using a python script provided by Pamir developers. We used 48 CPUs for more than ten days to run the *in-silico* sample. This tool requires high computational resources.

##### 3.2.12 AsmVar (version 2.0)

AsmVar detects DEL, DUP, INS, INV, and TRA. This tool uses the AS strategy to detect all SVs. Before the variant detection, we realigned the *in-silico* using the LAST aligner (de novo aligner), as developers recommended. This software required the following steps: 1) Variant detection, 2) Altalignemt, 3) Genotyping, 4) RecalibrationVg, and 5) a Variant Quality Score Recalibration (VQSR).

### 4 Benchmarking of different variant callers

#### 4.1 Recall, Precision and F-Score

Both *in-silico* and GIAB samples were used as a gold standard to benchmark the variant calling step. The metrics used to compare the accuracy of the variant callers are Recall, Precision and F-score. These metrics are calculated using True-Positive (TP), False-Negative (FN), and False-Positive (FP) detections. These parameters are evaluated for each SV type, SNVs, and indels independently.

The Recall is the fraction of TP variants detected among all variants in the sample:

$$Recall = \frac{TP}{TP + FN}$$

The Precision is the proportion of TP that are positives:

$$Precision = \frac{TP}{TP + FP}$$

The F-score is the harmonic mean of Precision and Recall:

$$F - score = 2 * \frac{Recall * Precision}{Recall + Precision}$$

For each variant caller, we did a preliminary variant filtering by using the flag PASS in the VCF output. We also discarded all variants detected in 1) Chromosome Y, 2) mitochondrial chromosome, 3) decoy sequences and 4) variants with genotypes "0/0" or "./.".

###### 4.1.1 Evaluation of indels breakpoint-error

Indel detection is at base-pair resolution. However, as the size of the indel increases, breakpoint detection becomes less accurate. For this reason, we studied the breakpoint-error as follows. Firstly, we normalised the outputs obtained by each caller from the *in-silico*/GIAB samples following the recommendations of the Global Alliance for Genomics and Health (GA4GH) (detailed documentation in section 8.1). Secondly, we merged the outputs of the callers with the *in-silico* at base-pair resolution. Finally, we calculated the recall for each size (1-50). Since the recall decreases from 30bp, we labelled small indels those variants with length ≤ 30bp. For indels from 31bp to 150bp, we labelled these variants as mid-size indels and associated a breakpoint-error of ± 10.

###### 4.1.2 The categorisation of *in-silico* variants

We grouped by variant type all genome variability of the *in-silico* to calculate the Recall, Precision and F-score metrics. We filtered out from the *in-silico* files, the variants with the label "Homozygotic" and counted once the duplications with the flag "tandem" (overview of all variants used to benchmark the variant callers are in Supplementary Fig.1).

Of note in the *de novo* insertion (INS) group, small variants from other variant types were further included; these additional variants were 32 Duplications, 2 Inversions, 1 Pseudogene, 47 Translocations and 37 small insertions (indels) (all numbers are obtained after filtering homozygous and duplicate variants), which were misclassified by variant callers, and were therefore added to the INS group, obtaining a total of 727 variants for INS benchmarking. (The input files used to construct the logistic regression models are deposited at https://github.com/gcatbiobank/GCAT_panel/tree/main/1_LRM)

###### 4.1.3 Determination of SVs breakpoint-errors

Knowing that the larger the length of SVs, the more discrepancies between callers to report the SV breakpoint, we evaluated the breakpoint-error for each caller by comparing the F-score metric of different windows from the breakpoint reported. We used window breakpoint-error of 10, 20, 50, 100, 200 and 300. The maximum window breakpoint-error considered coincides with the insert size of the *in-silico* (300bp). The range of the window breakpoint-error is calculated as follows:

$$Range_{lower\_point} = Position\ reported\ by\ caller - window\ breakpoint\_error$$

$$Range_{upper\_point} = Position\ reported\ by\ caller + window\ breakpoint\_error$$

We considered a variant as TP if the position reported by the in-silico overlapped with the window breakpoint-error range of the caller. Otherwise, it is classified as FP. Supplementary Table 4 shows the breakpoint-errors considered for each SV type.

###### 4.1.4 Evaluation of variant caller metrics

We evaluated the recall, precision and F-score for each variant caller, and variant type. Here, we describe the criteria to calculate these metrics for SNVs, small Indels, and SVs.

###### 4.1.4.1 SNVs/ and small Indels metrics

The strategy to calculate the metrics for SNVs and indels is described as follows. We classified the variants as TP if (i) the chromosome and the position reported by the algorithms overlap with the *in-silico* at base-pair resolution and (ii) the REF/ALT alleles are the same. Otherwise, it is classified as FP. All the variants not detected by the callers and included in the *in-silico* were FN.

###### 4.1.4.2 SVs metrics

The strategy to calculate the metrics for SVs is described as follows. We classified the variants as TP if (i) the SV type coincides with the *in-silico* (ii) the *in-silico* chromosome and position overlaps with the breakpoint-error of the variant caller (section 4.1.3) and (iii) the length of the variant reported by the caller is > 80% reciprocal overlap with the length of the *in-silico*. Otherwise, it is classified as FP. All the variants not detected by the callers and included in *the in-silico* were FN.

###### 4.1.4.3 Recall and Precision to detect variants by length

The size is a determinant factor for detecting the SVs. Larger SV sizes, the less mapping quality of the reads in the region. This leads to misinterpretations and increases the False-Positive detections. Then, for each SV type, we calculated the recall and precision by intervals of length (30,50], (50,75], (75,100], (100,125], (125,150], (150,300], (300,500], (500-1000], (1000,2000], (2000,3000], (3000, +Inf) (Fig. 2c, Supplementary Fig. 4).

###### 4.1.5 The Genotype error evaluation

Variant callers provide a genotype for each variant. Good genotyping is essential when generating the reference panel since it allows a higher quality imputation. For this reason, genotype calls of each variant caller were matched with the gold standard. The genotype error has been evaluated for heterozygous and homozygous using the following formula:

$$\frac{100 - Matched\ genotype}{All\ Genotypes}$$

All variants with a genotype matching the *in-silico* sample were defined as "Matched Genotype". "All Genotypes" refers to all the genotypes of variants reported by the variant caller.

#### 4.2 Variant callers discarded from the study

Here, we show the variant callers discarded from the study, either for computational reasons or because of the low metrics observed during the benchmarking process (Supplementary Table 3).

##### 4.2.1 Variant callers discarded for SNVs and INDELs

###### 4.2.1.1 Varscan2 (version 2.4.3)

Varscan2 was discarded mainly due to computational problems. Varscan2 is executed in a multiple sample mode in order to improve the calling and genotyping. We performed a preliminary study by running four samples altogether, generating in the first step a file of 753Gb in eight hours. The size of the outputs generates an incompatibility problem due to disk space limitations.

###### 4.2.1.2 Platypus (version 0.8.1)

Platypus was discarded due to computational problems; the assembly process generates computer reading problems bugging the execution when we ran different samples simultaneously. Furthermore, this caller provides the lowest recall in the benchmarking.

##### 4.2.2 Variant callers discarded for Structural Variants

###### 4.2.2.1 Pamir (version 1.2.2)

Pamir needed 22 days for running one sample and generated a file of almost 900 GB. It was discarded due to space and time limitations.

###### 4.2.2.2 AsmVar (version 2.0)

AsmVar execution was not possible due to its high memory requirements.

###### 4.2.2.3 Genome Strip (version 2.0)

Genome strip execution was not possible due to the incompatibility between the program architecture (LSF need) and the Marenostrum4 environment.

### 5 Increasing accuracy detection using a machine learning algorithm

To increase the variant detection accuracy of single algorithms, we developed a Logistic Regression Model (LRM). This model works using detection patterns of the variant callers and discriminative variables such as size, number of callers/strategies or reciprocal overlap. The LRM is suitable since the small number of features and the large number of variants considered. Furthermore, the LRM allows to estimate parameters indicating the accuracy of each caller and to make predictions based on the sum of the estimates in the logistic regression equation. This model was specific for each type of variant as detailed below (The Logistic regression models are deposited at https://github.com/gcatbiobank/GCAT_panel/tree/main/1_LRM).

#### 5.1 Logistic Regression Model for Indels

##### 5.1.1 Train and Test the model

The LRM for indels was trained using the GIAB sample and tested using the *in-silico* sample. The input of the LRM is a merged dataset of the VCF outputs from the callers following the same criteria to calculate the TP, FP and FN (section 4.1.4.1) (The script to merge the variant callers and generate the model is deposited at https://github.com/gcatbiobank/GCAT_panel/tree/main/ 1_LRM). We developed a specific LRM for indels using the R software (version 3.3.1) and the ISLR package. The function to fit the LRM is:

$$glm(PASS \sim Deepvariant + Haplotype\ caller + Strelka2, data = database, family = \text{“}binomial\text{”})$$

The outcome of the LRM is a binary variable (PASS) indicating if the variant is present in the GIAB sample. The independent variables are the genotypes reported by the variant callers indicating their detection pattern.

##### 5.1.2 Genotype reported by LRM for SNVs and small indels

The most common genotype reported by Haplotype caller, Deepvariant and Strelka2 is considered as the genotype of the LRM.

#### 5.2 Logistic Regression Model for SVs

##### 5.2.1 Train and Test the LRM for SVs

For SVs, the LRM was trained using 10-fold cross-validations for a random subset of variants (70%) from the *in-silico* and was tested using the remaining subset of variants (30%) of the *in-silico*. The input of the LRM is a merged dataset of the VCF outputs from the callers following the same criteria to calculate the TP, FP and FN (section 4.1.4.2).

We developed a specific LRM for each SV type independently using the *caret* (version 6.0-85) and *e1071* (version 1.7-3) R packages (The script to merge the variant callers and generate the models for each SV type are deposited at https://github.com/gcatbiobank/GCAT_panel/tree/main/1_LRM). The function to train the LRM is:

```
train(PASS ~ independent variables, data=database70, method="glm",  family="binomial", trControl= ctrl)
```

The outcome of the LRM is a binary variable (PASS) indicating if the variant is present in the *in-silico*. The independent variables are the genotypes reported by the variant callers, the size, number of callers/strategies and reciprocal overlap. Using stepwise backward criteria, we fitted a final LRM for each SV type (Supplementary Table 5 shows the discriminative variables used to build the LRM for each SV type).

###### 5.2.2 Strategy to report the position and length of variants

The strategy to report the position of the LRM is different for each SV type. We ordered the callers depending on (i) the accuracy in reporting the position in a breakpoint-error of ±10 bp and (ii) the number of variants detected (Supplementary Table 6). On the other hand, the variant length was considered as the median of the length reported by the callers that detected the variant.

###### 5.2.3 Genotype reported by LRM for SVs

The strategy to report the genotype of the LRM is different for each SV type (Fig. 2d, Supplementary Fig. 3).

1. DEL

Due to the high precision in the genotype calling, we reported the most coincident genotype between the genotype calls. When the consensus genotype was not possible to report, we included a missing "./.".

2. INS

We reported the genotype most coincident between the genotype calls. The sizes between 30 and 50 bp of Manta and Whamg detections were not used, because they do not report any genotype. When the consensus genotype was not possible to report, we included a missing "./.".

3. DUP

Since the genotyping error of callers is high, we developed a genotyping method using the BAM information of the *in-silico*. To obtain the total coverage of each DUP, we reported the median from all reads that cover twice the length of variant, in both upstream, downstream directions of the breakpoint reported by the LRM. The altered reads were obtained from the breakpoint reported by the LRM as follows. In a window of 10 bp, we counted the split reads, discarding the Hard-clipped reads and those containing INS or DEL in the CIGAR. Finally, we calculated the proportion of altered reads dividing by the total coverage.

$$\frac{Altered\ reads\ at\ breakpoint}{Total\ Coverage\ region}$$

If the proportion of altered reads was ≤ 0.20, we genotyped 0/0. If the proportion was between 0.20 and 0.80 we genotyped 0/1. If the proportion was ≥ 0.80, we genotyped 1/1 (The script to genotype duplications is deposited at https://github.com/gcatbiobank/GCAT_panel/tree/main/genotyping/Duplication_genotyping).

4. INV

We reported the genotype depending on the accuracy of each caller, with the exception of SvABA, due to its error. We ranked the callers by decreasing genotype error and reported the genotype based on the following order:

1. Lumpy 2. Pindel 3. Whamg 4. Delly2 5. Manta

5. TRA

The genotype error reported by the callers in TRA is high. For this reason, we re-genotyped the variants using the BAM information of the *in-silico*. The total coverage was obtained by counting all reads that cover the breakpoint reported by LRM in a window of 4 bp by chromosome. The altered reads were obtained by (i) discarding Hard-clipped reads, (ii) counting all reads with map quality ≥ 20 and (iii) counting reads with label different to 151M in the CIGAR. Finally, to report the genotype, we applied the same formula described for DUP (The script to genotype translocations is deposited at https://github.com/gcatbiobank/GCAT_panel/tree/main/genotyping/Translocation_genotyping).

###### 5.2.4 Filtering out the SVs variants following the GoNL strategy

GoNL constructed a panel of genetic variability, including SVs. The GoNL strategy is based on retaining variants that were detected by at least two different algorithms. We extended this strategy to three and four different algorithms and calculated the recall and precision, considering all these criteria (Fig. 2b, Supplementary Fig. 2).

### 6 GCAT project

In this paper, we use genetic data from the GCAT biobank project based on 19,267 volunteers recruited from the general population of Catalonia, in the northeast of Spain. The participants aged between 40-65 years with 16% non-Caucasian ancestry, mostly from American-Hispanic origin.

#### 6.1 Genomic data features

Available genetic data included SNP array-data (N=5,459; 56% female) and whole genome sequencing (WGS) data (N=808, 50.61% female), from which 71% samples overlap both datasets.

The genomic data was obtained in FASTQ format. BAM files were obtained by alignment of the sequencing reads to hs37d5 reference genome, applying the GATK Best Practices to clean up the BAM files (Supplementary Fig. 5) (FASTQ and BAM files are deposited at https://ega-archive.org/studies/EGAS00001003018).

#### 6.2 FASTQ files and BAM file generation

The FASTQ files of the 808 GCAT samples were generated in batches. Four batches included 192 samples, one batch included 24 samples, and the remaining batch included 16 samples. Each sample contains multiple LANEs grouped by the Multiplex index. Furthermore, the paired-end was generated in separated files:

| FASTQ paried-end 1 | FASTQ paired-end 2 |
|---|---|
| H52MCDSXX_1_60idt_1.fastq.gz | H52MCDSXX_1_60idt_2.fastq.gz |
| H52MCDSXX_2_60idt_1.fastq.gz | H52MCDSXX_2_60idt_2.fastq.gz |

The BAM files were built using the hs37d5 reference genome and following the GATK Best Practices. Firstly, we aligned each pair-end FASTQ file. Then, for each sample, we merged the BAM file according to the LANE.

BAM files for male and female samples were generated separately, since the number of reads and variants increases in female samples by removing the Y chromosome from the reference genome (Supplementary Fig. 5).

#### 6.3 Quality control (QC)

QC of the genetic data was evaluated by analysing the alignment quality of the BAM files, contamination traces and population structure of the WGS sample (Supplementary Table 10 shows all samples discarded).

##### 6.3.1 Alignment quality

Alignment quality was analysed by Picard (version 2.18.11), Biobambam (version 2-2.0.65) and Alfred (version 0.1.16). In total, 75.4% of samples fulfilled the QC standard metrics of 1000G and GoNL projects (Supplementary Table 9, Supplementary Fig. 6). The remaining samples were inspected across all thresholds, and the variant callers were executed correctly, determining that they passed the QC. One sample (EGA_16988) was discarded because of the irregularities observed when executing variant callers.

##### 6.3.2 Contamination analysis

We used VerifyBamID to determine the contamination or swapped ID samples. We used this tool with both genotyping data (array-based) and array-free approaches. The contamination thresholds of samples sequenced and array genotyped was [CHIPMIX] >> 0.02 and [FREEMIX] >> 0.02. Otherwise, if the sample was WGS, the threshold was [FREEMIX] ≥ 0.03 and [FREELK1]-[FREELK0]. For this purpose, we selected 570 GCAT samples from the SNP array data that were also sequenced. Using this array data and all 807 WGS BAM files, we ran VerifyBamID with -–best --ignoreRG --maxDepth 30 -–precise arguments. All the metrics obtained from VerifyBamID shown that any GCAT sample was contaminated.

##### 6.3.3 Population structure using reference ancestries

We generated a panel of genetic variability, including individuals from the Iberian population, based on Principal Component Analysis (PCA). Firstly, we ran the Haplotype Caller tool and selected the PASS filter variants from the VCF file. Then, we used PLINK (version 1.90b6.7 64-bit) to keep ~1M of SNVs with minor allele frequency (MAF) > 0.01 (discarding rare and monomorphic variants) and linkage disequilibrium (LD) $r^2$ < 0.2 (obtaining independent variants). Finally, we applied two PCAs based on different populations of known ancestries: (i) Using worldwide populations from the 1000Genomes project, and we discarded 16 GCAT samples from non-European ancestry (Supplementary Fig. 7a and Supplementary Fig. 7b). (ii) Using the Population Reference Sample (POPRES) project available in the webserver LASER (https://laser.sph.umich.edu/) and the 1000G European samples, 2 GCAT samples were discarded from non-Iberian ancestry according to the mean ± 4sd criteria (Supplementary Fig. 7c).

##### 6.3.4 Identity by Descent analysis

We used PLINK (version v2.00a2LM) to estimate the Identity by Descent (IBD probabilities) of 789 samples of GCAT. We identified one full-sibling pair and one first-cousin relationship. These pairs of individuals shown probabilities of sharing 0, 1 and 2 IBD alleles equal to (0.3,0.48,0.22) and (0.78,0.22,0), which are close to the theoretical values (0.25,0.5,0.25) and (0.75,0.25,0) for full-siblings and first-cousins respectively. We discarded one sample per pair according to the maximum number of missing values in the genotype data (Supplementary Fig. 7d).

###### 6.3.5 Population structure without reference ancestries

To detect potential outliers from other populations not present in 1000G or POPRES projects, we applied an additional PCA without reference ancestries. We discarded from this analysis two additional GCAT samples (Supplementary Fig. 7e). After all the QC steps applied, we used 785 samples to construct the panel of genetic variability of the Iberian population.

#### 7 Variant calling

Here, we describe the variant calling process for the GCAT cohort in multi-sample mode to improve the detection and genotyping of the variants. The Y chromosome has been called in the subset of male samples (Supplementary Fig. 8 overview all steps to run each variant caller). Supplementary Table 11 shows the computational time consumed to perform the variant calling of 785 samples.

##### 7.1 SNVs and Indels calling

###### 7.1.1 Haplotype Caller

Haplotype Caller was run for each sample (section 3.1.1) adding the flags (--ERC GVCF, --dbsnp dbsnp_138.b37.vcf, --L chrm --G Standard Annotation, --G StandardHCAnnoation). Then, we combined all samples using the GenomicsDBimport module in batches of 1 MB. We re-genotyped the variants with the GenotypeGVCF module (flags -V gendb://merge, --G StandardAnnota tion, --new-qual). Next, we applied the Variant Quality Score Recalibration (VQSR) by running the VariantRecalibrator (flag –AS) and ApplyVQSR modules separately by SNVs and indels (flag –module). Finally, we executed the genotype refinement using the CalculateGenotyePosteriors module, and we filtered out variants with genome quality < 20 using the VariantFiltration module.

###### 7.1.2 Strelka2

Strelka2 was run by sample as described in section 3.1.3. Strelka2 is not able to run in multi-sample mode if the chromosome number is different between samples. Then, we could not run all samples together because female samples do not have the Y chromosome in the BAM file, and males include it.

###### 7.1.3 Deepvariant

Deepvariant was run as described in section 3.1.2. After applying the three Deepvariant modules, we genotyped all variants using the next two GATK modules. We combined the 785 samples by batches of 1 MB, with the CombineGVCFs module. Next, we genotyped all variants using the GenotypeGVCF module.

#### 7.2 **Mid-size and Structural Variant calling**

##### 7.2.1 Delly2

Delly2 was run as described in section 3.2.1. We could not apply all Delly2 pipeline to call TRA events because three samples provided time and computational limitations; for this reason, we only applied the first step to detect TRA events. For the remaining SV types, we run the MERGE and CALL BCF modules to combine and re-genotype all variants. Next, we used Bcftools to merge all BCFs. Finally, we used the FILTER module to delete the redundant variants and find the confident germline SV.

##### 7.2.2 Manta

Manta was run as described in section 3.2.2. Similar to Strelka2, we could not execute the multi-sample algorithm due to BAM construction issues (section 7.1.2).

##### 7.2.3 Pindel

Pindel was run by chromosome and sample as described in section 3.2.3. For each sample, we included the insert size obtained from Picard (section 0) in the config file. Due to computational requirements, we could not execute the pipeline for eight samples in chromosome 2. We converted the Pindel format files to VCF using the pindel2vcf module. This module cannot convert the translocation file to VCF format. To genotype the large insertions, we applied our custom genotyper, described in section 3.2.3.

##### 7.2.4 Lumpy

Lumpy and SVTYPER were run as described in section 3.2.4. We re-genotyped all variants in the multi-sample mode as follows. Firstly, we used the lsort and lmerge (flag: -f 50) SVTOOLS modules to combine the VCFs. Then, we re-genotyped all variants using SVTYPER in batches of 15K variants.

##### 7.2.5 SvABA

SvABA was run by sample as described in section 3.2.6.

##### 7.2.6 Whamg

Whamg was run by sample as described in section 3.2.5. This tool detects the SVs without genotypes. Then, we used SVTYPER as described in section 3.2.4 to genotype variants.

###### 7.2.7 CNVnator

CNVnator was run by sample as described in section 3.2.73.2.7. We used the read-length (150 bp) in the bin size parameter.

###### 7.2.8 Popins

Popins was run as described in section 3.2.8. with additional steps, allowing the execution in a multi-sample mode. First, we executed by sample the steps 1) assembly, 3) contigmap and 7) genotype. Then, we executed for all samples the steps 2) merge, 4) place-refaling, 5) place-splitalign, and 6) place-finish. To genotype variants, we used the -m RANDOM label.

###### 7.2.9 Melt

Melt was run in multi-sample mode using the MELT-SPLIT pipeline. This pipeline is composed of (i) Pre-processing BAM files, (ii) Individual analysis, (iii) Group analysis, (iv) Genotype and (v) Make VCF.

### 8 Variant Calling integration

#### 8.1 VCF pre-processing

###### 8.1.1 SNV and indels set

We normalised the VCF outputs from Deepvariant, Haplotype Caller and Strelka2, before merge them. For Haplotype Caller and Deepvariant, we used the SelectVariants GATK module (including the flags: --exclude-filtered, –-sm sample, --remove-unused-alternates) to split each multi-sample VCF before normalising. VCF normalisation was performed with the pre.py tool, designed by the Global Alliance for Genomics and Health (GA4GH). We run pre.py as recommendations by developers, including the options: --L, --decompose, --pass-only, –-threads. Finally, we created a single VCF per sample and chromosome, discarding multiple nucleotide variants (MNP) and variants that do not specify the ALT allele.

###### 8.1.2 Structural Variants set

We removed the low quality/NO_PASS variants, the 0/0 or "./." variants, those not mapped in autosomal and sexual chromosomes, variants catalogued as NOANCHOR in the VCFs from Popins, and the BNDs variants that have the same chromosome in both breakpoints in the VCFs from Lumpy. Finally, the variants were divided by SVTYPE into different files.

#### 8.2 <u>Merging all VCFs per-sample and variant type</u>

For each sample, we merged the VCF outputs from the variant calling as follows. For SNVs and indels, we merged variants by (i) chromosome and position at base-pair resolution and (ii) same REF/ALT alleles. For SVs, we merged variants by (i) variant type, (ii) chromosome, (iii) overlapping window breakpoint-error of the variant (section 4.1.3) and (iv) reciprocal overlap ≥ 80% between callers. Using these merged datasets, we applied the LRM (section 5.2) by variant type and sample. Based on the predicted LRM results, for each variant, we considered as TP if the LRM prediction was ≥ 0.5. Otherwise, we considered the variant as FP. For SNVs, we considered a variant as TP if it was detected by at least two callers, not using the LRM. On the other hand, we reported the maximum breakpoint-error, the number of callers, the number of strategies that detected the variant, the breakpoint position, genotype and length of the variant, following the strategy described in sections 5.1.2, 5.2.2 and 5.2.3 (The scripts to merge variant caller outputs per sample are deposited at https://github.com/gcatbiobank/GCAT_panel/tree/main/2_merge_callers).

#### 8.3 <u>Combining all samples in one VCF</u>

For each variant type, we combined the quality-controlled GCAT samples (785) as follows. For SNVs and indels, we merged individuals by (i) chromosome and position at base-pair resolution and (ii) REF/ALT alleles. For SVs, we merged individuals by (i) variant type, (ii) chromosome, (iii) position with a maximum breakpoint-error of the merged variant and (iv) reciprocal overlap ≥ 80% between individuals. Using these merged datasets, we calculated for each variant the proportion of TP determined by LRM. **This proportion is referred to as a <u>quality score</u> of the merged variant**. We considered a variant PASS if the quality score is ≥ 0.5. Then, we reported the length and position of each SV as the median length and median position of all the samples that have the SV. Additionally, we added in the VCF file the following population information: allelic count (AC), minor allelic count (MAC), allele frequency (AF), minor allele frequency (MAF), population variation (POPVAR) (POPVAR is calculated according to the minor allele); and the following variant-specific information: breakpoint-error (ERRBKP) and SVTYPE (The scripts to merge samples and construct the multi-sample VCF file are deposited at https://github.com/gcatbiobank/GCAT_panel/tree/main/3_merge_samples).

#### 8.4 <u>Variant Quality Control</u>

For each variant type, we applied a standard QC. We considered the PASS variants obtained from LRM (quality score > 0.5), and we discarded monomorphic variants, variants out of Hardy-Weinberg Equilibrium (Bonferroni correction p-value < $5x10^{-8}$), and those with ≥ 10% of missingness (The GCAT catalogue with all variants accepted after filtering step is deposited at http://cg.bsc.es/GCAT_BSC_iberianpanel/) (The GCAT catalogue with genotype information is deposited at https://ega-archive.org/studies/EGAS00001003018).

### 9 Validation of variants

#### 9.1 Public datasets comparative

##### 9.1.1 SNVs and Indels

We compared the final set of SNVs and indels (section 8.4) with the NCBI dbSNP Build 153 (dataset downloaded from https://ftp.ncbi.nlm.nih.gov/) as a reference sample. We merged the dbSNP dataset and GCAT variant set by (i) chromosome and position at base-pair resolution and (ii) REF/ALT alleles. We determined the number of variants shared in both datasets and the number of unique variants in the GCAT variant set (Supplementary Fig. 9a, Supplementary Fig. 9b).

##### 9.1.2 Structural Variants

We compared the final set of SVs with (i) The Genome Aggregation Database (gnomAD.v.2) (https://gnomad.broadinstitute.org/downloads), (ii) Database of Genomic Variants (DGV) (http://dgv.tcag.ca/dgv/app/downloads?ref=GRCh37/hg19), (iii) Human Genome Structural Variation Consortium set (HGSVC) http://ftp.1000genomes.ebi.ac.uk/vol1/ftp/data_collections/hgsv_sv_discovery/working/20181025_EEE_SV-Pop_1/VariantCalls_EEE_SV-Pop_1/) (iv) Hall-lab dataset (https://github.com/hall-lab/sv_paper_042020), (v) 1000 Genomes (Phase3) (ftp://ftp.1000genomes. ebi.ac.uk/vol1/ftp/phase3/) and (vi) GoNL (release 6.2) as gold standards. For each database, we merged each database and GCAT variant set by (i) variant type, (ii) chromosome, (iii) ±1000 bp breakpoint-error and (iv) reciprocal overlap > 80% between datasets. We determined the number of variants shared in at least one dataset, and the number of unique variants in the GCAT variant set (Supplementary Fig. 9c, Supplementary Fig. 9d).

#### 9.2 Experimental validation

##### 9.2.1 SNV and Indel validation using SNP-array genotyping

We validated the SNV and indel final sets (section 8.3) with the GCAT SNP array (https://ega-archive.org/datasets/EGAD00010001664) as follows. First, we selected 570 GCAT samples that have WGS and SNP array data. Then, we merged both datasets by (i) chromosome, (ii) position at base-pair resolution and (iii) same REF/ALT alleles. Finally, we calculated the recall and genotype concordance for each sample based on the 732,978 SNVs and 1,168 indels included in the SNP array (Supplementary Fig. 10).

###### 9.2.2 Inversions validation using a verified dataset

We used a golden set of inversions experimentally validated from the InvFEST project to check the accuracy of the GCAT inversion dataset. Each validated variant provides the genomic coordinates (GRCh37), length and allele frequencies in the genotyped populations from 3 main continental groups: Africa (YRI and LWK), Europe (CEU and TSI) and East-Asia (CHB and JPT). Due to the limitations of short reads to detect variants in repetitive regions, we considered 59 inversions mediated by Non-Homologous (NH) mechanisms that do not have inverted repeats at their breakpoints. Then, we merged this golden set and the GCAT final inversion set (section 8.4) by overlapping a window of ±1000 bp. Finally, allele frequency (using European frequencies) and length concordances of inversions from the GCAT and InvFEST datasets (Supplementary Fig. 11a, Supplementary Fig. 11b).

On the other hand, to evaluate the accuracy of inversion genotyping, we compared GCAT predictions for the 785 sequenced participants and imputation calls using a reference panel of experimentally validated inversion genotypes provided by InvFEST[24]. NH inversions were imputed in GCAT samples with IMPUTE v2.3.2. Inversion genotypes were called with a posterior probability higher than 0.8 and were classified as missing otherwise. If the inversion had perfect tag SNPs ($r2 = 1$) that were present in the GCAT panel, missing genotypes were recovered based on the tag SNP genotypes. Finally, genotype agreement was estimated between GCAT genotypes and InvFEST imputation calls (Supplementary Fig. 11c).

#### 10 Creation and integration set of haplotypes

##### 10.1 Benchmarking of different phasing strategies

To benchmark different phasing strategies, we evaluated the imputation accuracy of SVs by using SNP array for a subset of 95 GCAT individuals that have WGS and SNP-array data. We considered the quality of imputed variants as a validation proxy of the best phasing strategy. The phasing algorithms used were ShapeIt2 (version v2.r904), MVNcall (version 1.0), ShapeIt4 (version 4.1.3) and WhatsHap (version 0.18). We used IMPUTE2 (version 2.3.2) for imputation analysis (Supplementary Fig. 12, Supplementary Fig. 13).

###### 10.1.1 Sample pre-processing

We created a reference panel for chromosome 22 with 690 individuals using WGS data. Chromosome 22 panel contained 195,106 SNVs, 24,321 Indels and 128 large Deletions (>150bp) (section 8.4). Phasing and imputation accuracy of this panel was estimated, using as the gold standard the WGS genotypes from an independent subset of 95 individuals also characterised by SNP-Array technology and including 7,146 SNPs. We applied into Chromosome 22-panel variants a quality-control using PLINK; HWE test (--hwe 1.639629e$^{-07}$ midp (Bonferroni Correction)), filtering out variants with ≥ 10% of missing (--geno 0.1), removing singletons and doubletons (--maf 0.0026).

##### 10.1.2 Phasing strategies

###### 10.1.2.1 ShapeIt2 (with SVs)

ShapeIt2 is a tool mainly used to phase SNPs and indels and is used in similar projects such as 1000G and GoNL to create their haplotype set. For this reason, we evaluated the capacity of Shapeit2 to phase all variants, including SVs. We executed ShapeIt2 as follows:

```
shapeit --input-vcf
snps_indels_del_to_phase_HWE_no_double_single QC01.vcf --
input-map genetic_map_chr22_combined_b37.txt --output-max
snps_indels_del_to_phase_HWE_no_double_singleQC01.haps
snps_indels_ del_to_phase_HWE_no_double_singleQC01.legend --
thread 48
```

###### 10.1.2.2 ShapeIt2 and MVNcall

MVNcall is commonly used to phase MNPs (Multiple Nucleotide Polymorphism), complex indels and SVs. MVNcall requires a haplotype scaffold of SNPs and indels that we constructed using Shapeit2 (section 10.1.2.1). This phasing strategy was used to create an SV-resolved haplotype set in the 1000G and GoNL projects [42,43]. MVNcall requires GL (Genotype Likelihood) or PL (Phred-scaled Genotype Likelihoods). As we genotyped the variants from WGS using high coverage (30X), we used PL as 0 in the genotype reported by the merge, and 255 for the remaining genotype probabilities. Variants with "./." genotypes were discarded. We executed MVNcall as follows:

```
mvncall --int 1 51304566 --sample-file SNP_indels_GCAT1.sample
--glfs
Del_final_to_phase_nodots_no_singleton_doubleton_hwe.vcf --sca
ffold-file
SNPs_indels_final_to_phase_HWE_no_double_singleQC01.haps  --
lambda= 0.1 --o Del_final_to_phase_mvncall_1_51304566_all.vcf
```

###### 10.1.2.3 ShapeIt2, MVNcall with PIRs

ShapeIt2 can use the sequencing reads to improve the phasing of rare variants for samples with high coverage. The reads that span two heterozygote sites are considered as phase informative reads (PIRs) and are considered as mini-haplotypes. We downloaded the PIR module (extractPIRs.v1.r68.x86_64.tgz) from (https://mathgen.stats.ox.ac.uk/genetics_software/ shapeit/shapeit.html), and we executed this module only for SNVs. Then, we run ShapeIt2 (section 10.1.2.1), including the flag –input-pir. Finally, MVNcall was executed as described in section 10.1.2.2.

###### 10.1.2.4 ShapeIt4

ShapeiIt4 uses the positional Burrows-Wheeler transform (pBWT) algorithm. This method stores the haplotypes at each iteration in a pBWT data structure. Before running

```
shapeit4 --input
snps_indels_del_to_phase_HWE_no_double_singleQC01. vcf.gz --
map genetic_map_chr22_combined_b37.txt --output SNP_indels_
SV_GCAT_all.vcf --pbwt-depth 8 --seed 123456 --region 22 --
thread 48 --sequencing --log SNP_indels_SV_GCAT_all_ok.log
```

ShapeIt4, we compressed with bgzip and indexed with tabix the variants. Then, we ran Shapeit4 as follows:

Finally, we executed Bcftools (convert -–haplegendsample) to convert the vcf obtained to the files .hap, .legend, and .sample, required to execute IMPUTE2.

###### 10.1.2.5 ShapeIt4 and MVNcall

We applied the same strategy described in section 10.1.2.2, changing the ShapeIt version.

###### 10.1.2.6 ShapeIt4 and WhatsHap

ShapeIt4 uses the WhatsHap tool to extract the PIRs by grouping heterozygous genotypes into phase sets when the same sequencing reads overlap them. However, this tool only can extract PIRs from SNVs and indels. WhatsHap cannot parallelise executions. Then, to increase the performance, we divided the multi-sample VCF into individual VCFs, one for each sample. We ran WhatsHap as follows:

```
whatshap phase -o
sample_snps_indels_SV_filtered_final_to_phase_QC01_
shapeti4_whatshap.vcf.gz --tag PS --reference genome.fa --
indels              --chromosome 22
snps_indels_SV_filtered_final_to_phase_QC01_shapeit4_
sample_ok.vcf.gz
/gpfs/scratch/bsc05/bsc05138/GCAT1/01_GCAT/samples_GCAT_01/CWG
S026/bam/bam/CWGS026_bam_final/bam_basequality/CWGS026.bam
```

Next, we combined all individual VCFs in a multi-sample VCF file. Finally, we executed Shapeit4, including the flag --use-PS 0.0001, and Bcftools as described in section 10.1.2.4.

##### 10.1.3 Imputation using different phasing strategies

We imputed 95 GCAT individuals using a different reference panel depending on the strategies described in the previous section. Before imputation,10.1.1 we applied a “pre-phased” to reduce the computational costs without compromising the accuracy. We used the flags --input-ref and -H for Shapeit2 and Shapeit4, respectively. Then, we run IMPUTE2 for each reference panel as follows:

```
impute2 -use_prephased_g -m
genetic_map_chr22_combined_b37.txt   -h
reference_panel.hap.gz -l all_snps_
reference_panel.legend.gz -known_haps_g
gcat_test_imputation.hap.gz -int 5 Mb batch -o
output_file.gen -o_gz
```

###### 10.1.4 Imputation accuracy

We evaluated each phasing strategy by counting the number of variants with an info score $\geq 0.7$ and by calculating the genotype concordance between imputed data and NGS. Based on our results, slight differences were obtained between Shapeit4 and Shapeit4 + WhatsHap strategies. The differences came from the imputation of rare variants, where Shapeit4+WhatsHap improved their imputation quality (Supplementary Fig. 12, Supplementary Fig. 13). For this reason, we used Shapeit4+WhatsHap to phase the GCAT samples.

##### 10.2 Pipeline to construct the GCAT haplotype panel

We followed the strategy described in section 10.1.2.6 to create the haplotype-resolved panel of GCAT using the 785 samples. We built a haplotype panel for each autosomal chromosome and X chromosome. To generate the haplotype-resolved panel of chromosome X, we separated chromosome X by pseudo-autosomal regions 1 and 2 (PAR1 and PAR2) and non-pseudo-autosomal regions (NOPAR). Then, for male samples, we coded the heterozygous genotypes in NOPAR as "./.". On the other hand, to perform the chromosome X imputation, we included two specific flags –chrX and -sample_g, and we grouped by gender the samples from the array (Supplementary Fig. 14 shows the pipeline followed) (GCAT|Panel is deposited at https://ega-archive.org/studies/EGAS00001003018).

#### 11 Imputation using the GCAT reference panel

##### 11.1 Imputation analysis using the GCAT SNP-array

Similar to section 10, we created a whole-genome pilot reference panel of 690 GCAT individuals. The variants included in this panel were obtained after filtering out variants with $\geq 10\%$ of missing and removing monomorphic variants with PLINK. Using this pilot reference panel, we evaluated the imputation accuracy of 95 independent GCAT individuals using 754,593 SNPs from the SNP-array that passed strict Quality Control (Galván-Femenía et al.). We considered the WGS variant calling of the same 95 independent GCAT individuals as a reference, and we compared the genotype concordance of the imputed variants (info score ≥ 0.7) and the WGS-called genotypes (Fig. 4a, Supplementary Fig. 15, Supplementary Fig. 16). Then, we evaluated the number of variants that are in linkage disequilibrium (LD, $r^2 > 0.2$) with each common SV type (MAF ≥ 0.05) in a window of 1 MB, to relate the imputation genotype concordance with patterns of low LD (Supplementary. 17, Supplementary Fig. 18).

Additionally, we assessed the effect of the presence of SVs in the SNV and indel imputation by exploring the info imputation by counting the number of SNVs and indels imputed without a reference panel of SVs (Supplementary Table 12).

##### 11.2 **Imputation quality using the array of 1000G**

The GCAT panel is composed of Iberian samples. To evaluate the efficiency to impute variants in populations from other ethnicities, we used the 1000G array data (ftp://ftp.1000genomes.ebi.ac.uk/vol1/ftp/release/20130502/supporting/hd_genotype_chip/ALL.chip.omni_broad_sanger_combined.20140818.snps.genotypes.vcf.gz). This array contains 2,318 samples (48.1% male) from 19 populations and 5 continental groups with 2,458,861 SNPs. Supplementary Table 13 shows the origin of 19 populations and their continents.

###### 11.2.1 Sample filtering and Quality Control of the 1000G array

The array data was cleaned up, removing samples or markers of questionable quality. We downloaded the file igsr_samples.tsv from https://www.internationalgenome.org/data-portal/sample to evaluate the sample gender. Then, we filtered out from the array 41 samples with unknown gender. We discarded 395 related samples until 2nd degree, using the file 2013606_g1k.ped downloaded from ftp://ftp.1000genomes.ebi.ac.uk/vol1/ftp/technical/working/ 20130606_sample_info/. We discarded two Masai samples due to low sample size, obtaining an array of 1,880 samples. Finally, the array was divided by the 19 populations included in the data. PLINK was used to check variants for missings (--geno 0.1), discarded A-T, CG sites, applied HWE test (--hwe), discarded related samples (--rel-cutoff 0.05), individuals with an excess of heterozygosity ± 2sd (--het) and with missing call rate ≥0.1. For chromosome X, we separated males and females and divided the PAR and NONPAR regions.

###### 11.2.2 Imputation study of GCAT panel with non-Iberian samples

Using the GCAT reference panel (section 10.2), we imputed the 19 populations from 1000 Genomes separately (section 10.1.3). To evaluate the imputation of SVs in non-Iberian samples, we used as a reference the SV characterisation performed by Audano using long reads. We downloaded the file EEE_SV-Pop_1.ALL.sites.20181204.vcf from ftp://ftp.1000genomes.ebi.ac.uk/vol1/ftp/data_collections/hgsv_sv_discovery/working/20181025_EEE_SVPop_1/VariantCalls_EEE_SV-Pop_1/. This file contains 15 samples sequenced with PacBio (GRCh38 aligned) and SVs called with SMRT-SV tool. The samples HG00514, HG00733, NA19240, NA19434, HG01352, HG02059, NA12878, HG02106, HG00268 were used to evaluate the precision and recall of the imputation of 1000 Genomes as follows. We evaluated the concordance of SVTYPE and variant length between Audano's characterisation and GCAT dataset. Additionally, we discarded the variants with genotypes 0/0 and "./.". Then, we considered a TP if an imputed variant (info score ≥ 0.7) in a window of ±50 bp overlapped with an SV in Audano's set (Supplementary Fig. 19).

To calculate the genotype concordance, we used as a reference the genotype reported by Hickey project. This project genotyped three samples from Audano's file (HG00514, HG00733, and NA19240) using short reads and applying a variation graph, implemented in the vg toolkit. We downloaded the files svpop-vg-HG00514.vcf.gz, svpop-vg-HG00733.vcf.gz, svpop-vg-NA19240.vcf.gz from https://s3-us-west-2.amazonaws.com/human-pangenomics/index.html? prefix=vgsv2019/vcfs/. All variants with "./." were discarded. Also, the genotypes from TRA, TRP, INS, DUP of imputation were compared with all INS reported by Hickey. In both Audano's and Hickey's files, we converted the coordinates of the SV calls from GRCh38 to GRCh37 using the liftOverPLink.py tool (Supplementary Fig. 19b, Supplementary Fig. 20).

#### 11.3 Preliminary discoveries using the GCAT|Panel

##### 11.3.1 Genome-Wide Association Analysis

Association analysis was performed for 70 multiples independent GWAS in selected phenotypes. Phenome analysis included only chronic conditions defined by linked Electronic Health Records from the cohort registry (2012-2017). Chronic conditions were selected according to the indicator of the chronic condition and grouping the resulting conditions by similarity, considering ICD-9 codes and descriptions. Conditions with more than 50 cases were retained for the GWAS analysis (i.e. 70). Each genomic wide association tests were performed as independent logistic regression for each cohort under the assumption of an additive model for allelic effects, with adjustments made for age, sex, and the first five principal components. Gender-specific conditions were analysed only for a specific gender. Analysis was performed using PLINK2.0 for autosomal chromosomes. Locus Zoom was derived for specific regions, and suggestive tower profiles were analysed, based on LD patterns and gene centred impact. Bonferroni correction was considered for 10 independent body systems (Fig. 5a, Supplementary Fig. 28, Supplementary Fig.29).

###### 11.3.2 Experimental validation of the Alu element

PCR amplicon analysis was designed using Primer 3.0 software using the hg19_dna range=chr3:49,492,813-49,496,062 sequence, including the Alu element. Sequence primers are for F-primer, (5'CATTGACTCATTCAGCAAGCA 3') and for R-primer (5'AAATTAAGCCCCACCCTAG3'). Using standard conditions (35X, Tm=60ºC) in a Veriti™ 96-Well Thermal Cycler (Thermo Fisher Scientific), we obtain a 515 bp fragment corresponding to the control-allele and a 848 bp for the Alu- allele. Fragments were resolved by e-agarose gel, in a TapeStation (Agilent). Further, the amplicon of a non-ALU allele carrier was analysed by Sanger Sequence Method to verify the insertion point (i.e. at hg19 Chr3:49,494,280) and the ALU sequence insertion (324 bp) (Fig. 5b).

#### 12 Benchmarking different panels of genetic variability

We imputed 4,988 individuals and 756,773 SNVs from the GCAT array (section 10.1.1) using different population reference panels: 1000G phase3, GoNL-SV, UK10K and HRC. Multiple reference panel imputation was conducted using GUIDANCE. First, using PLINK, we double-checked for missingness by sample and chromosome, and we discarded three samples with missing call-rate ≥ 0.10. Next, we filtered out samples that were present in the array and the GCAT reference panel. We considered imputed variants with MAF > 0.001 and info imputation score ≥ 0.7, in order to compare the number of imputed variants of each reference panel. To be consistent with other projects, the SVs were all variants with a length ≥ 50 bp.

We compared the number of unique variants imputed by the different panels as follows. For SNPs and indels, we considered the same variant between panels if the position and ALT allele coincided. For SVs, we considered the same variant between panels if the SV type overlapped in a window of ±1,000 bp (Fig. 4b, Fig. 4c). To assess the quality of imputation by allele frequency, we filtered out the variants with MAF < 0.001, and we calculated the average of the info score ($r^2$) by rare (MAF < 0.01), low frequency (0.01 ≤ MAF < 0.05) and common (MAF ≥ 0.05) variants (Supplementary Fig. 21).

#### 13 Biological impact of Structural variants

##### 13.1 Structural Variant distribution in worldwide populations

We evaluated the allele frequency, type of variant distribution and quality of imputed SVs in different human populations. We imputed each 1000 Genomes population separately with the Iberian reference panel (section 11.2). We used Shapeit4 to pre-phase and IMPUTE2 to impute the variants.

We used the concordance table metrics provided by IMPUTE2 (.summary file) to assess the genotype concordance for each population. We considered the values with the most confident imputed genotypes (max prob ≥ 0.9) and depicted the median of the genotype concordance of all imputed batches (1 MB) per population (Supplementary Fig. 22).

The SV distribution among different populations was determined using the .gen_infofile provided by IMPUTE2. This file contains information on frequency (exp_freq_a1) and certainty (info) of imputed variants. We evaluated SVs with info score ≥ 0.7 and the length ≥ 50 bp (Fig.6, Supplementary Fig. 23).

#### 13.2 Functional implications of Structural Variant in humans

##### 13.2.1 Structural variant annotation using AnnotSV

We used AnnotSV to provide functional, regulatory and clinical annotations of SVs. This tool uses multiple datasets to evaluate the effect of SVs in humans. We ran AnnotSV by using default parameters in the multi-sample VCF by chromosome and with a reciprocal overlap parameter of 80% (-overlap 80) to be consistent with our merge algorithm (Section 8) (The catalogue of SVs annotated is deposited at http://cg.bsc.es/GCAT_BSC_iberianpanel/) (The catalogue of SVs annotated with genotype information is deposited at https://ega-archive.org/studies/EGAS00001003018).

We studied the SV annotation analysing: (i) the SV distribution in human genes (Supplementary Fig. 24), (ii) level of overlap with regulatory regions, (iii) predicted loss of function intolerance (pLI) effect of SVs in human (Supplementary Fig. 25) and (iv) associated diseases (Supplementary Fig. 26, Supplementary Table 14).

##### 13.2.2 Evaluation of SVs using GWAS catalog

We used the genome-wide association study (GWAS) catalog version 1.0 (e98 r2020-03-08) to analyse reported loci in GWAS with our defined SVs set; we counted several variants tagged by SNPs. For the analysis, we considered autosomal variants, with MAF > 0.01 in the GCAT database and being reported associated with European ancestry. In brief, 106,906 GWAS associations corresponding to 68,323 unique SNPs were evaluated from the GWAS catalog. Intersection with defined breakpoints positions in our SVs set, evidenced 4,733 SNP GWAS associations (2,669 unique SNPs) having at least one structural variant in strong linkage disequilibrium ($r^2 > 0.80$ threshold in a 1 MB window) (Supplementary Fig. 27).

### 14 Supplementary Tables and Figures

#### 14. 1 Supplementary Tables

| Features of sequencing | Parameters |
|---|---|
| **Sequencing system** | HiSeq 2000 |
| **Read length** | 100 bp |
| **Insert size** | 500 bp |
| **Inner mate distance** | 300 bp |
| **Simulation** | Paired-end |
| **Coverage per allele** | 15 X |

**Supplementary Table 1. Sequencing description of *in-silico* sample**. We sequenced each haplotype separately using ART tool; for this reason, the coverage of sequencing (15X) is half of the total coverage in the *in-silico* (30X). *bp = Base Pair*

| FLAG | VARIANT TYPE | Description | TOTAL EVENTS |
|---|---|---|---|
| *Germline* | SNV | Germline variants from 1000G | 4.871.660 |
| *Germline* | Insertion/deletion | Germline variants from 1000G | 454.995 |
| *PointMutation* | SNV | Point Mutations variants from PanCancer | 723 |
| *indel* | Insertion/deletion | Indels from PanCancer | 3.052 |
| *no_indel* | Deletion | Deletions > 100bp from PanCancer | 269 |
| *consensus* | Deletion | Deletions 101>9000bp from PanCancer | 26 |
| *big_del* | Deletion | 9 largest Deletions (>10kbp) in in-silico obtained from PanCancer | 9 |
| *random_transDel_IDtranslocation* | Deletion | Deletions associated with a non-reciprocal translocation randomly generated(complex event) | 674 |
| *random_SVtype_flank* | Deletion | Small deletions flanking with another SV randomly generated | 8 |
| *random_ins_new* | Insertion | Insertions randomly generated | 440 |
| *random_SVtype_shard* | Insertion | Genomic shards inserted between breakpoints in variants >200bp randomly generated | 9 |
| *consensus_inv* | Inversion | Inversions from PanCancer | 14 |
| *random_inv* | Inversion | Inversions randomly generated | 241 |
| *random_dup* | Duplication | Duplications randomly generated near to the original sequence (tandem dup) | 458 |
| *random_dup_inv* | Duplication | Duplications randomly generated and inverted near to the original sequence | 17 |
| *random_dup_(inv)_tandemnum* | Duplication | Tandem duplications event that occurs as many times as the number that accompanies the flag randomly generated | 62 |
| *consensus_tran* | Duplication | Duplications obtained from PanCancer | 14 |
| *random_trans_num* | Translocation | Non-reciprocal translocations randomly generated associated with a deletion (*idtranslocation*) | 610 |
| *translocation_chrA_chrB, translocation_chrB_chrA,* | Translocation | Reciprocal translocation randomly generated | 2 |
| *virus_intron, virus_intron_complete, virus_random* | Virus | Retroviruses obtained from NCBI affecting intronic regions with sequence truncated, complete sequence or randomly inserted | 89 |
| *random_GEN* | Pseudogene | Randomly generated Pseudogenes ( GEN = gene-name related) | 9 |
| *random_retrotransposon_nametransposon* | Transposon | Transposons obtained from Repbase database | 100 |
| *random_SVtype_Homozygotic* | All SV Variants | Homozygotic variants generated randomly | 1.150 |
| *Variant_in_chr12_dup* | All Variants | Chr12 was duplicated, all the variants duplicated have this flag | 56 |

**Supplementary Table 2. Description of variant types inserted in the *in-silico* sample**. All variants which are obtained from 1000G or PanCaner project have the flag consensus, Germline, PointMutation, indel or no_indel. Python scripts generated the variants with flag random. The transposons and viruses are obtained from NCBI or Repbase databases and are inserted randomly in the genome.

| Variant Caller | Strategy | Variant type |
|---|---|---|
| Haplotype Caller | SR, AS | SNVs, Indels, Mid DEL |
| Deepvariant | ML | SNVs, Indels, Mid DEL |
| Strelka2 | AS | SNVs, Indels, Mid DEL |
| Delly2 | SR, DR, RD | Mid DEL, DEL, DUP, INS, INV, TRA |
| Manta | AS | Mid DEL, DEL, DUP, INS, INV, TRA |
| Pindel | SR, DR | Mid DEL, DEL, DUP, INS, INV, TRA |
| Lumpy | SR, DR, RD | Mid DEL, DEL, DUP, INV, TRA |
| Whamg | SR, DR, ML | Mid DEL, DEL, DUP, INS, INV |
| SvABA | AS, SR, DR | Mid DEL, NO SV TYPE |
| CNVnator | RD | DEL, DUP |
| Popins | AS | INS |
| MELT | DR | TRP |
| Platypus | AS | SNVs, Indels, Mid DEL |
| VarScan2 | SR | SNVs, Indels |
| Genome Strip | SR, DR, RD | DEL, DUP, mCNV |
| Pamir | SR, DR, OEA | INS |
| AsmVar | AS | DEL, DUP, INS, INV, TRA |

**Supplementary Table 3. Variant callers evaluated to analyse the genome variability.** To perform an accurate variant detection, we evaluated different variant callers based on their strategy and variant detection, covering all genome variability. SR= Split read; DR= Discordant Read; RD= Read-depth; AS= de novo Assembly; ML= Machine Learning; OEA; One End Anchored. The variant callers coloured in red were discarded for variant detection in GCAT samples.

Mid DEL = Deletions size ranges 30-150 bp; DEL = Deletions >150; INS = Insertions ≥ 30 bp; DUP = Duplications ≥ 50 bp; INV = Inversions ≥ 50 bp; TRP = Transposons; TRA = Translocations; mCNV = Multiple Copy Number Variation

| Callers | Strategy | Breakpoint Resolution (31-150 bp) Mid DEL | Breakpoint Resolution (150 bp ≥) DEL | Breakpoint Resolution (50 bp ≥) DUP | Breakpoint Resolution (50 bp ≥) INV | Breakpoint Resolution (50 bp ≥) INS | Breakpoint Resolution (50 bp ≥) TRA |
|---|---|---|---|---|---|---|---|
| Delly2 | SR+DR+RD | ±10 | ±100 | ±10 | ±10 | ±10 | ±300 |
| SvABA | AS+SR+DR | ±10 | ±100 | ±10 | ±10 | ±10 | ±10 |
| Manta | AS | ±10 | ±50 | ±20 | ±10 | ±10 | ±200 |
| CNVnator | RD | None | ±300 | ±100 | None | None | None |
| Whamg | SR+DR+ML | ±10 | ±10 | ±10 | ±10 | ±10 | None |
| Lumpy | SR+DR+RD | ±10 | ±100 | ±50 | ±10 | None | ±200 |
| Popins | AS | None | None | None | None | ±10 | None |
| Pindel | SR+DR | ±10 | ±10 | ±10 | ±10 | ±10 | ±10 |

**Supplementary Table 4. Selected Breakpoint-error of each variant caller by SV type.** SV detections do not have base-pair resolution. To combine different variant callers, we first determined the accuracy to correctly report the SV position. The breakpoint-error allowed us to determine if a SV detected by different tools was the same since the breakpoint-errors overlapped.

None: The software does not detect this variant type.

| Mid-size Deletion | DEL | INS |
|---|---|---|
| Deepvariant, Haplotype Caller, Strelka2, Manta, Whamg, Delly2, Lumpy, Pindel, SvABA, Num of callers detected, Reciprocal overlap ≥ 0.8 | Manta, Whamg, Delly2, Lumpy, Pindel, SvABA, CNVnator, Variant size, Strategy | Manta, Whamg, Delly2, Pindel, SvABA, Popins, Haplotype caller, Strelka2, Num of callers detected |
| **INV** | **DUP** | **TRA** |
| Manta, Whamg, Delly2, Pindel, SvABA, Lumpy, Variant size | Manta, Whamg, Delly2, Lumpy, Pindel, SvABA, CNVnator, Variant size, Reciprocal overlap ≥ 0.8 | Manta, Delly2, Pindel, SvABA, Lumpy |

**Supplementary Table 5. Independent predictor variables used to train the Logistic Regression Model for each SV type.** Variant caller variables are the genotypes indicating the presence or the absence of the variant. The **variant size** is divided into different ranges, (i-e. 500-1000, 1000-2000 bp, 2000-3000, > 3000), each range is a predicted variable. The **number of detection callers** indicates the number of different algorithms detecting each variant (i.e. 2, 3, 4-5). Each number is a predicted variable. **The reciprocal overlap ≥0.8**, is a numerical variable indicating the size of the overlapping between two different algorithms. The **strategy** is the number of different strategies used by callers to detect a variant.

| Structural Variant Type | Order |
|---|---|
| Mid-Deletion/Deletion | Pindel > Whamg > Delly2 > Manta > Median of remaining callers |
| Insertion | Pindel > Delly2 > Strelka2 > SvABA > Manta > Median of remaining callers |
| Duplication | SvABA > Pindel > Delly2 > Whamg > Median of remaining callers |
| Inversion | Lumpy > Pindel > Delly2 > Median of remaining callers |
| Translocation | Manta > Median of remaining callers |

**Supplementary Table 6. Order to report the breakpoint by Logistic Regression Model.** The most precise algorithm was used to report the breakpoint by the merge algorithm. When the breakpoint-error was large, we reported the median.

| Study purpose | Fraction sample | Vacutainer tube | Volume mL | Transport T°C | Time to PMPPC | Aliquots n (T°C) | Control assay |
|---|---|---|---|---|---|---|---|
| **Genomic/ epigenomic** | Buffy coat | EDTA | 10 | 4 | max 24 hours | 2 (−80) | SNP array, qPCR, PCR, STR |
| | Highly concentrate d buffy coat | Blood bag | 480 | 18 | max 48 hours | 2 (−80) | SNP array, qPCR, PCR, STR |

**Supplementary Table 7. Description of sampling for Genomic analysis in the GCAT project.** This table is adapted from Obón-Santacana et al. PMPPC, Program of Predictive and Personalised Medicine of Cancer. STR, short tandem repeat

| Study purpose | Number of participants | Fraction sample | Platform | Machine | Analysed |
|---|---|---|---|---|---|
| **Genotype** | 5,459 | Buffy coat | Infinium Multi-Ethnic Global (MEGAEX2) array | HiScan confocal scanner (Illumina) | $2\times10^6$ SNPs, InDels |
| **Whole-genome Sequencing (WGS)** | 808 | Buffy coat | Illumina TruSeq PCR free/Illumina paired-end SBS | HiSeq 4000 sequencer (Illumina) | 30X coverage Read length 150 bp Insert size 600 |

**Supplementary Table 8. Summary of Genomic data of the GCAT project.** This table is adapted from Obón-Santacana et al.

| **QC metrics and Inclusion criteria** |
|---|
| Fraction purified reads > 0.90 |
| Fraction reads aligned in pairs > 0.95 |
| 0.495 < Strand Balance < 0.505 |
| 250 bp < Mean insert size < 350 bp |
| Standard deviation of insert size < 50 bp |
| *Fraction of duplicated reads < 0.1 |
| 27X < Mean Coverage < 37X |
| + Fraction of paired reads mapped in the same chromosome > 0.88 |

**Supplementary Table 9. Alignment Quality control metrics.**

* Biobambam2 tool; + Alfred tool

| Sample discarded | BAM low quality | Non-Iberian representative | Familiar relatedness |
|---|---|---|---|
| EGA_07970 | | X | |
| EGA_05253 | | X | |
| EGA_06951 | | X | |
| EGA_07438 | | X | |
| EGA_03681 | | X | |
| EGA_02098 | | | X |
| EGA_07185 | | X | |
| EGA_18629 | | X | |
| EGA_00980 | | | X |
| EGA_07947 | | X | |
| EGA_04194 | | X | |
| EGA_02397 | | X | |
| EGA_16770 | | X | |
| EGA_16000 | | X | |
| EGA_02096 | | X | |
| EGA_13093 | | X | |
| EGA_03662 | | X | |
| EGA_06463 | | X | |
| EGA_13871 | | X | |
| EGA_16988 | X | | |
| EGA_06827 | | X | |
| EGA_03692 | | X | |

**Supplementary Table 10. Samples discarded from the GCAT cohort after sample filtering.**

| Variant Caller | Total Time (hour) | CPU/h | % |
|---|---|---|---|
| CNVnator | 977.03 | 11,724.34 | 0.34 |
| Whamg +SVTyper | 2,122.87 | 24,384.13 | 0.71 |
| SvABA | 1,867.2 | 29,875.18 | 0.87 |
| Strelka2 | 999.05 | 49,414.24 | 1.45 |
| Popins | 4,201.68 | 69,094.31 | 2.02 |
| Manta | 3,194.05 | 153,314.4 | 4.48 |
| Deepvariant | 13,184.25 | 336,301.2 | 9.84 |
| Haplotype Caller | 72,152.83 | 609,258.2 | 17.82 |
| Lumpy | 416,645.28 | 627,228.5 | 18.35 |
| Pindel | 107,925.77 | 647,554.6 | 18.94 |
| Delly2 | 143,393.36 | 860,375.3 | 25.17 |
| **Total computational time consumed** | **766,663.37** | **3,418,524** | **100** |

**Supplementary Table 11. The Computational time to perform the variant calling of 785 GCAT samples.** The computational resources are a bottleneck of variant calling. For example, Delly2, Lumpy, Haplotype caller and Deepvariant are executed using all samples together, investing a high amount of time in their executions. The electricity costs needed to perform this variant calling is around 820,329 €. Lumpy, Delly, Haplotype caller and Deepvariant invested most of the overall CPU time, due to re-genotype all variants using all samples together.

| Variant type | Iberian-GCAT panel | Genotype concordance (imputation vs calling) | Variants imputed with info score ≥ 0.7 |
|---|---|---|---|
| INDEL | Panel with SV | 93.97% | 44.50% |
| INDEL | Panel without SVs | 93.96% | 44.50% |
| SNV | Panel with SVs | 97.75% | 39.13% |
| SNV | Panel without SVs | 97.75% | 39.13% |

**Supplementary Table 12**. **Impact of structural variants on imputation.** The imputation quality of SNVs and indels was not influenced by the inclusion of SVs in the Iberian-GCAT reference panel neither in terms of genotype concordance.

| Population Code | Population Description | Super Population |
|---|---|---|
| ACB | African Caribbeans in Barbados | AFR |
| ASW | Americans of African Ancestry in SW USA | AFR |
| CDX | Chinese Dai in Xishuangbanna, China | EAS |
| CEU | Utah Residents (CEPH) with Northern and Western European Ancestry | EUR |
| CHB | Han Chinese in Beijing, China | EAS |
| CHS | Southern Han Chinese | EAS |
| CLM | Colombians from Medellin, Colombia | AMR |
| FIN | Finnish in Finland | EUR |
| GBR | British in England and Scotland | EUR |
| GIH | Gujarati Indian from Houston, Texas | SAS |
| IBS | Iberian Population in Spain | EUR |
| JPT | Japanese in Tokyo, Japan | EAS |
| KHV | Kinh in Ho Chi Minh City, Vietnam | EAS |
| LWK | Luhya in Webuye, Kenya | AFR |
| MXL | Mexican Ancestry from Los Angeles, USA | AMR |
| PEL | Peruvians from Lima, Peru | AMR |
| PUR | Puerto Ricans from Puerto Rico | AMR |
| TSI | Toscani in Italia | EUR |
| YRI | Yoruba in Ibadan, Nigeria | AFR |

**Supplementary Table 13. Populations used in imputation study from 1000G.**

| Phenotypes | num SVs | % of SVs associated with diseases |
|---|---|---|
| Cardiomyopathy, dilated, 3B, 302045 (3)/ Becker muscular dystrophy, 300376 (3)/ Duchenne muscular dystrophy, 310200 (3) | 26 | 4.48 |
| Epileptic encephalopathy, early infantile, 12, 613722 (3) | 25 | 4.30 |
| Mental retardation, AD 33, 616311 (3)/ (Ventricular fibrillation, paroxysmal familial, 2), 612956 (3) | 24 | 4.13 |
| Acrodysostosis 2, with or without hormone resistance, 614613 (3) | 20 | 3.44 |
| Mental retardation, XL 21/34, 300143 (3) | 15 | 2.58 |
| Pitt-Hopkins-like syndrome 2, 614325 (3)/ (Schizophrenia, susceptibility to, 17), 614332 (3) | 15 | 2.58 |
| Mental retardation, AD 39, 616521 (3) | 13 | 2.24 |
| Mental retardation, AR, 6, 611092 (3) | 12 | 2.07 |
| Cerebellar ataxia, nonprogressive, with mental retardation, 614756 (3) | 10 | 1.72 |
| Koolen-De Vries syndrome, 610443 (3) | 10 | 1.72 |

**Supplementary Table 14. Top 10 diseases related to SVs using the OMIM database.** The SVs evaluated for this analysis where 581 SVs associated with genes with high pLI and Haploinsufficiency. Mental and muscular diseases were the top 10 diseases related to deleterious SVs.

SV type associated with diseases are in Supplementary Figure 19.

**a**

| Population Frequency | Exons | Introns | Total |
|---|---|---|---|
| Common | 293 | 5,119 | 5,412 |
| Low frequency | 271 | 3,451 | 3,722 |
| Rare | 4,151 | 27,779 | 31,930 |

**b**

| Population Frequency | Exons | Introns |
|---|---|---|
| Common | 0.054 | 0.946 |
| Low frequency | 0.073 | 0.927 |
| Rare | 0.130 | 0.870 |

**Supplementary Table 15**. **a,** Number of SVs overlapping coding regions (Exons) and non-overlapping coding regions (introns) by common (MAF ≥ 0.05), low frequency (0.01 ≤ MAF < 0.05) and rare (MAF < 0.01) population frequencies. **b,** Proportion of SVs overlapping coding regions (exons) and non overlapping coding regions (introns) for common (MAF ≥ 0.05), low frequency (0.01 ≤ MAF<0.05) and rare (MAF < 0.01) population frequencies.

#### 14.2 Supplementary Figures

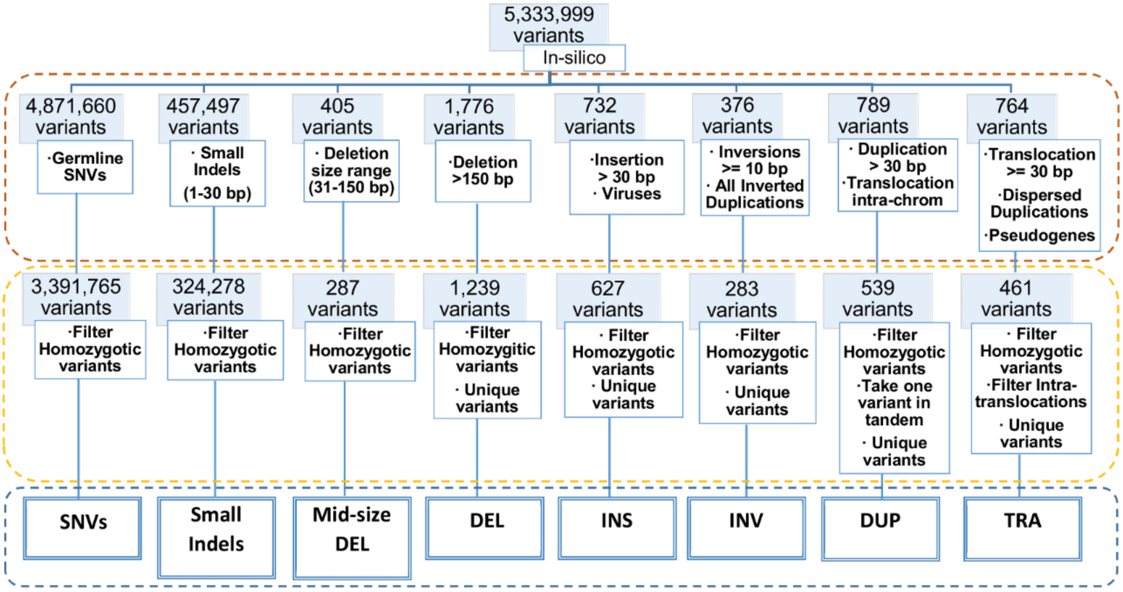

**Supplementary Fig. 1. Classification and filtering of the *in-silico* variants by type of variant**. Each **dashed line** colour ( - - - ; - - - ; - - - ) indicates the grouping and filtering steps, and different variant type respectively. The **white boxes** of grouping and filtering, indicate the criteria used to obtain the unique variants of each variant type. The **blue boxes** shows the number of variants after grouping and filtering step. In the grouping step, we also included those variants where the callers detect the breakpoint, but fail the variant type. The variants obtained after filtering were used to benchmark and generate the Logistic Regression Model, one for each SV type and indels (except SNVs).

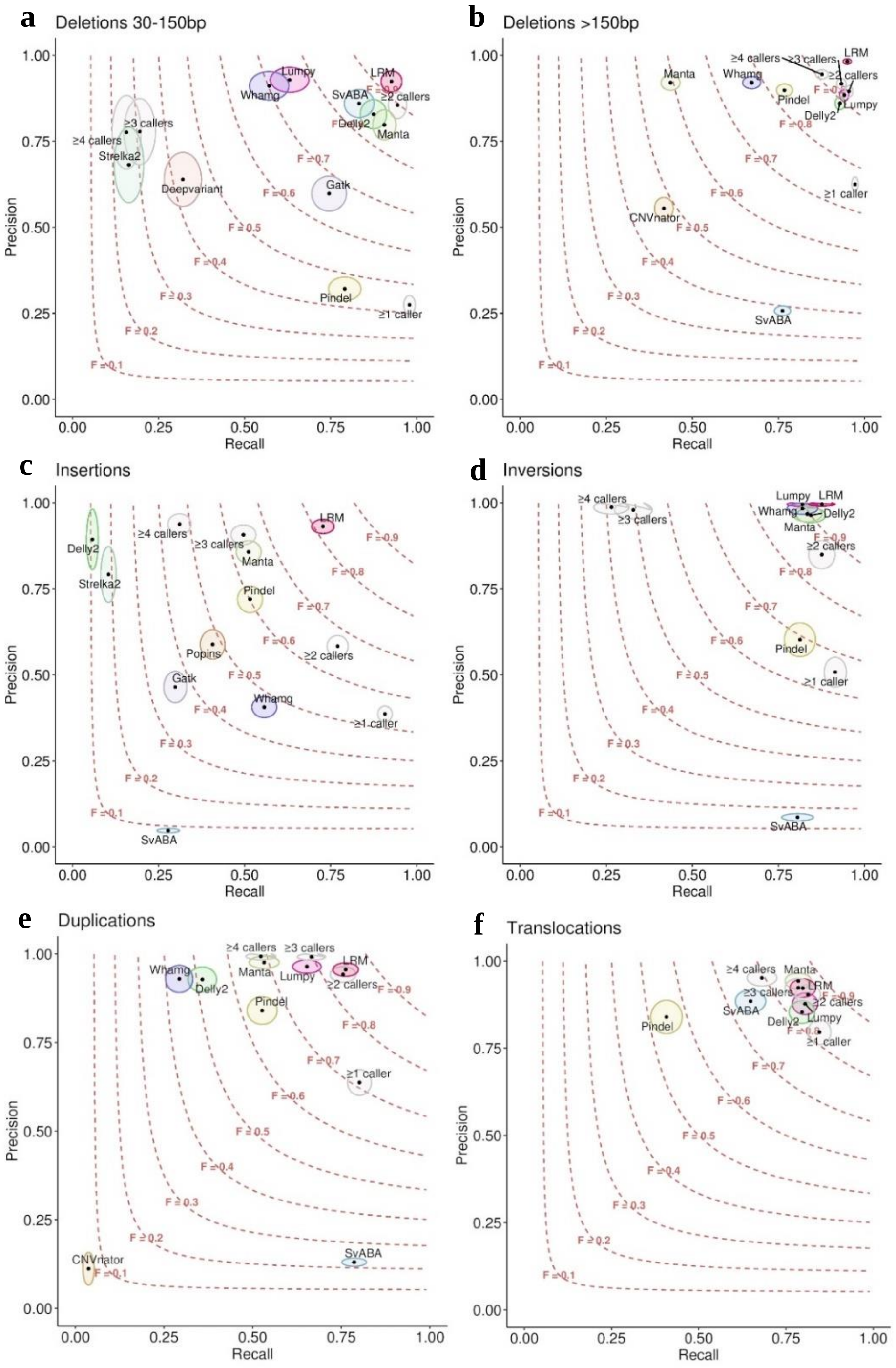

**Supplementary Fig. 2 | Variant calling benchmarking analysis classified by variant type.** Precision and recall distribution for each variant caller, logic rule (≥) and Logistic Regression Model (LRM), categorised by SV type. Dashed line indicate the F-score threshold divided by chuncks of 0.1. The Confidence intervals are represented as colored areas for each algorithm. **a, Deletions (30-150 bp).** The LRM outperformed the variant discovery (obtaining an F-score 0.93 and precision > 0.90) based on logical rules or variant callers individually. **b**, **Large Deletions (> 150 bp).** The LRM outperformed the variant discovery with an F-score > 0.95 and precision > 0.98. **c**, **Insertions.** The LRM is the most accurate strategy to detect insertions, obtaining an F-score 0.82 with high precision 0.93. **d, Inversions.** The LRM shows similar discovery performance compared with callers individually with 0.93 of F-score. However, the LRM shows larger F-score than logical rules. **e, Duplications.** No difference is noticeable between LRM and ≥2 callers' strategies. However, the LRM is 0.013 more precise than ≥2 callers. **f**, **Translocations.** No difference is observed between the most accurate variant caller and LRM.

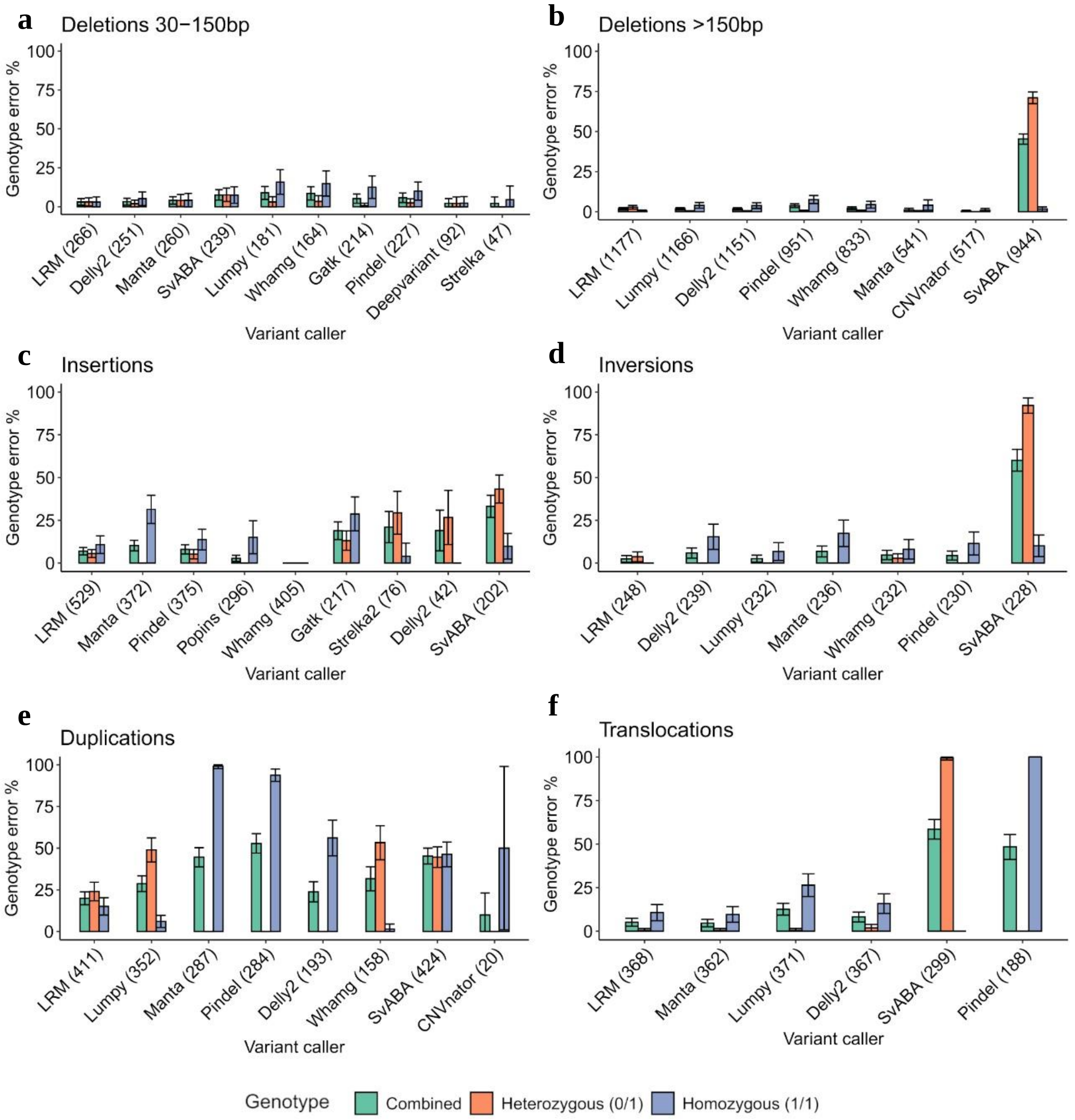

**Supplementary Fig. 3 | Comparative analysis of the genotype error associated with variant callers and Logistic Regression Model (LRM).** The orange bar indicates the genotyping error for variants that are heterozygous in the *in-silico* sample (0/1). The blue bar indicates the genotyping error for variants that are homozygous in *in-silico* sample (1/1). The green bar indicates the overall error for all detected variants. For all the structural variant types, the LRM proportionally showed a lower genotyping error than the different callers independently. The highest genotype error was reported in duplications (e), with 20% of genotypes.

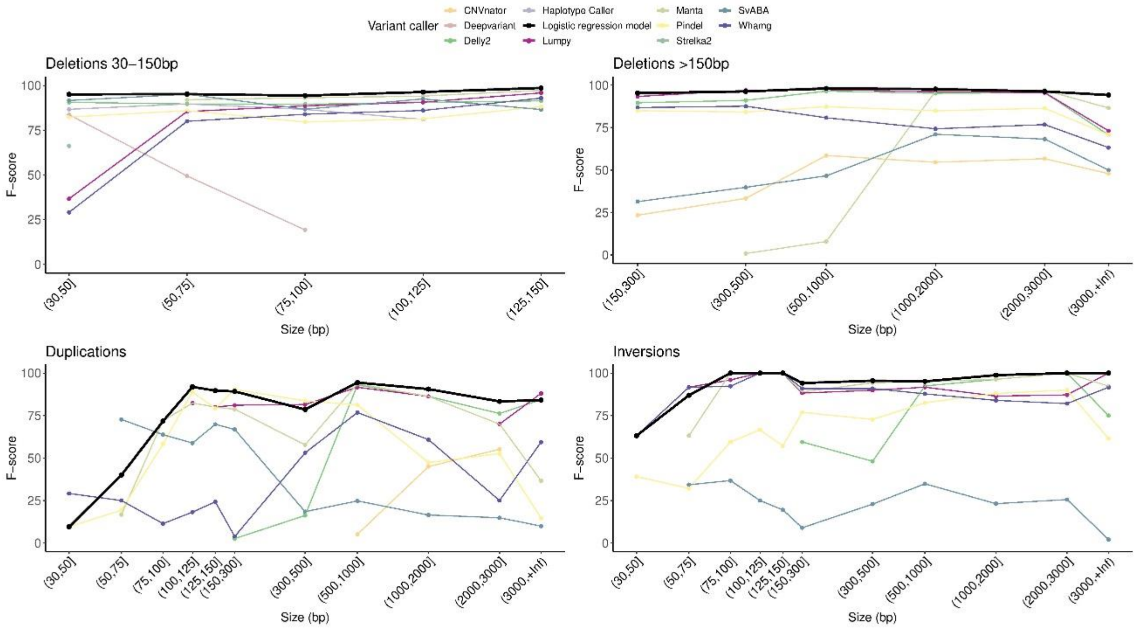

**Supplementary Fig. 4. Variant discovery grouped by SV type and classified by SV size.** The F-score of variant callers and Logistic Regression Model (LRM) fluctuated across SV sizes, demonstrating different accuracy in specific size ranges. For deletions, the F-score of the Logistic regression model is between 97-99%, depending on the SV size, demonstrating that the variable size improved SV discoveries' accuracy in specific size ranges. For Duplications and Inversions, the F-score of variant callers fluctuated across SV sizes. For this reason, including the SV size in Logistic Regressio Model (LRM), could improve the SV discovery of variant callers individually, increasing the discovery performance.

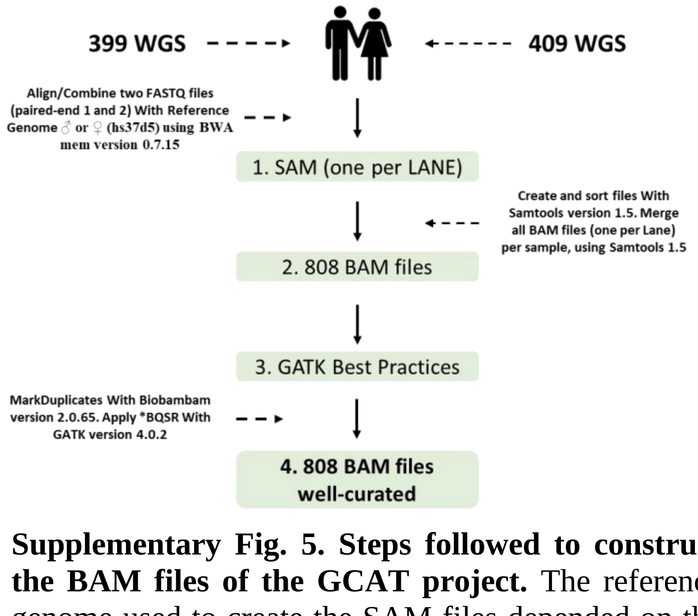

**Supplementary Fig. 5. Steps followed to construct the BAM files of the GCAT project.** The reference genome used to create the SAM files depended on the sample gender. We cleaned the BAM files following the Best Practices of GATK. Overall ~100 Terabytes were necessary to Store all BAM files.

* BQSR: Base Quality Score Recalibration

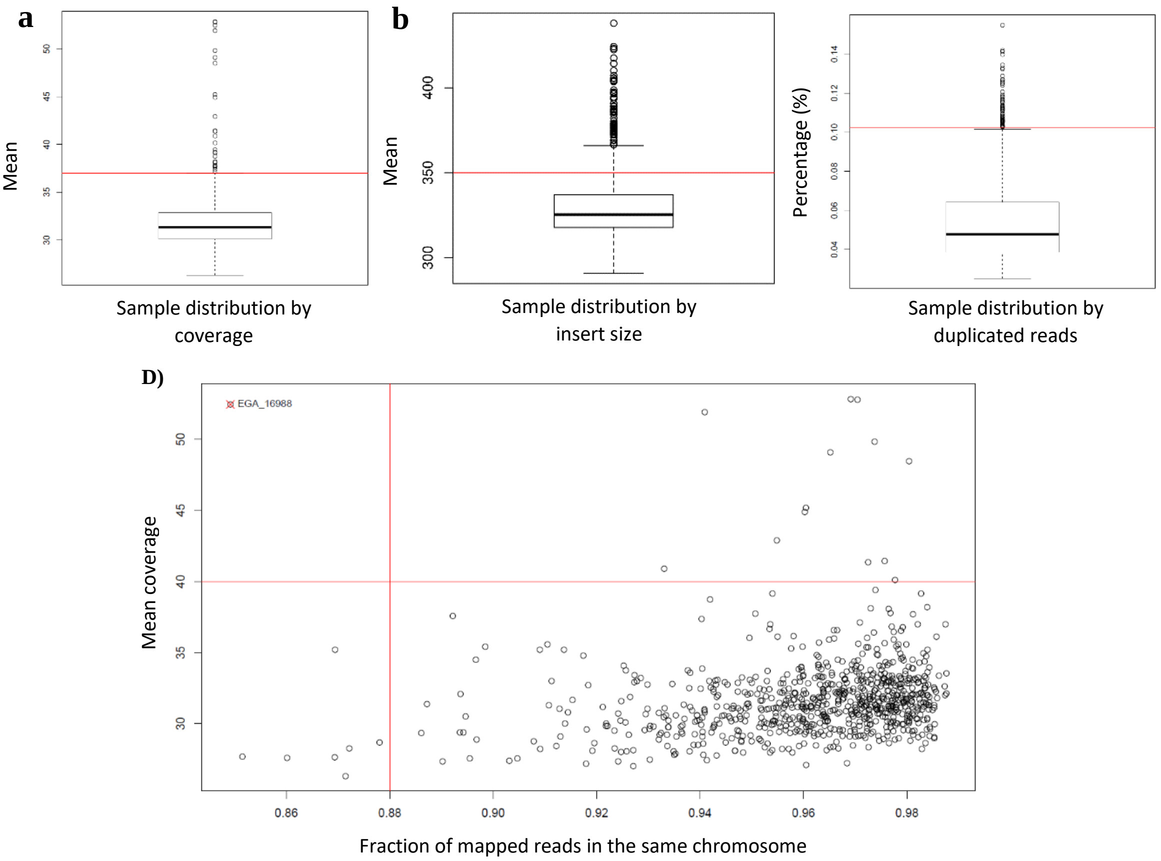

**Supplementary Fig. 6**. **Sample distribution based on QC metrics. a,** Mean coverage. 29 samples exceeded the threshold. **b,** Insert size distribution. 112 samples exceeded the threshold. **c,** Fraction of duplicated reads. 58 samples exceeded the threshold. **d,** Fraction of mapped reads in the same chromosome. 7 samples exceeded the threshold. One sample exceeded the threshold of coverage mean > 40 and the fraction of mapped reads in the same chromosome. This sample was discarded for variant calling irregularities. Horizontal and vertical lines crossing the panel indicate quality thresholds noted in Supplementary Table 9.

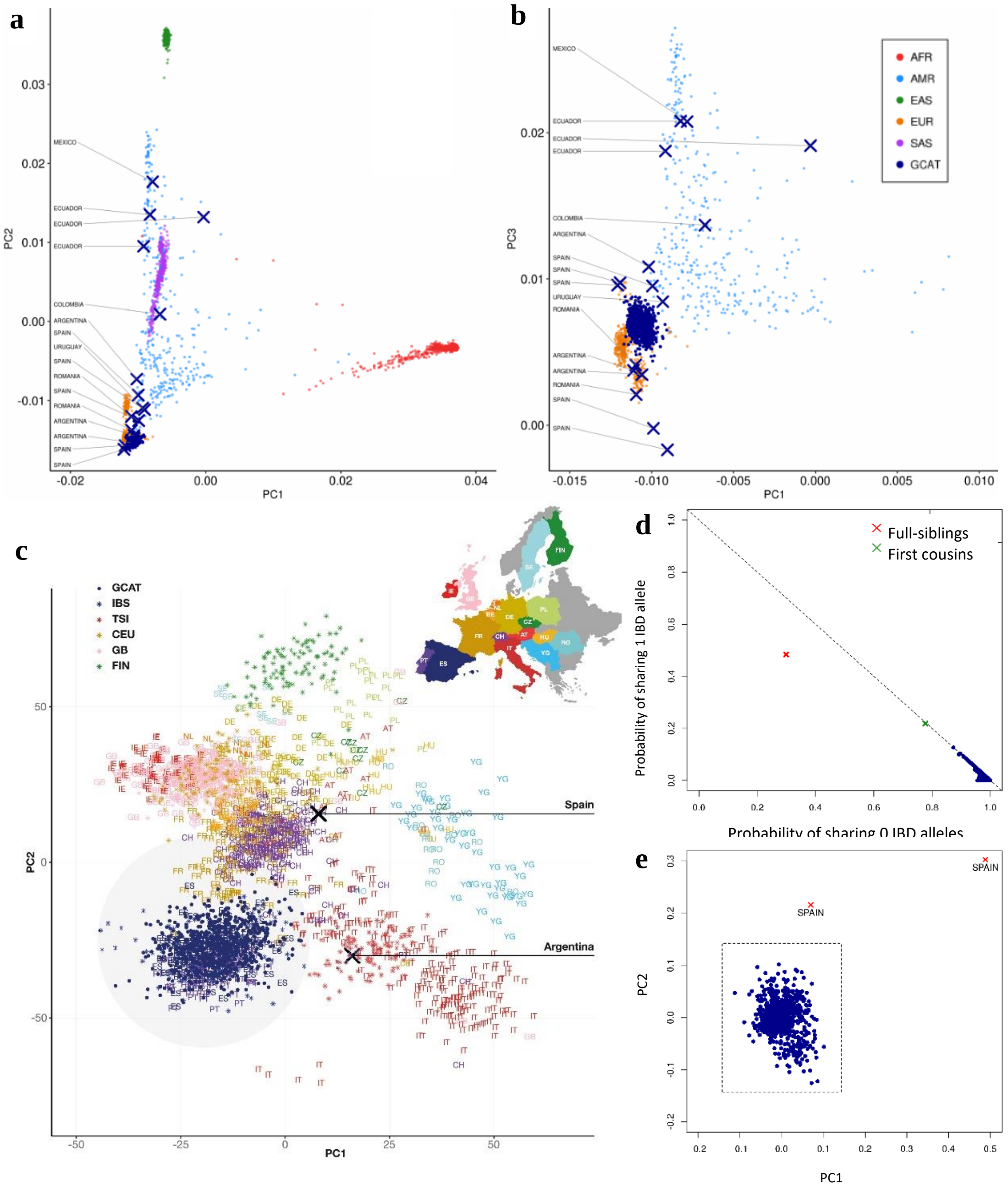

**Supplementary Fig. 7. Evaluation of the homogeneity in the GCAT cohort. a,** Distribution of the GCAT samples among 1000 Genomes, with PC1 and PC2. **b,** Distribution of the GCAT samples among 1000 Genomes, with PC1 and PC3. **c,** GCAT samples distribution among 1000 Genomes European samples and PROPES project using PC1 and PC2. **d,** Identity by Descent test (IBD). **e,** GCAT homogeneity, with PC1 and PC2.

X: Discarded Sample; ⁎: 1000G European samples; PC: Principal Component

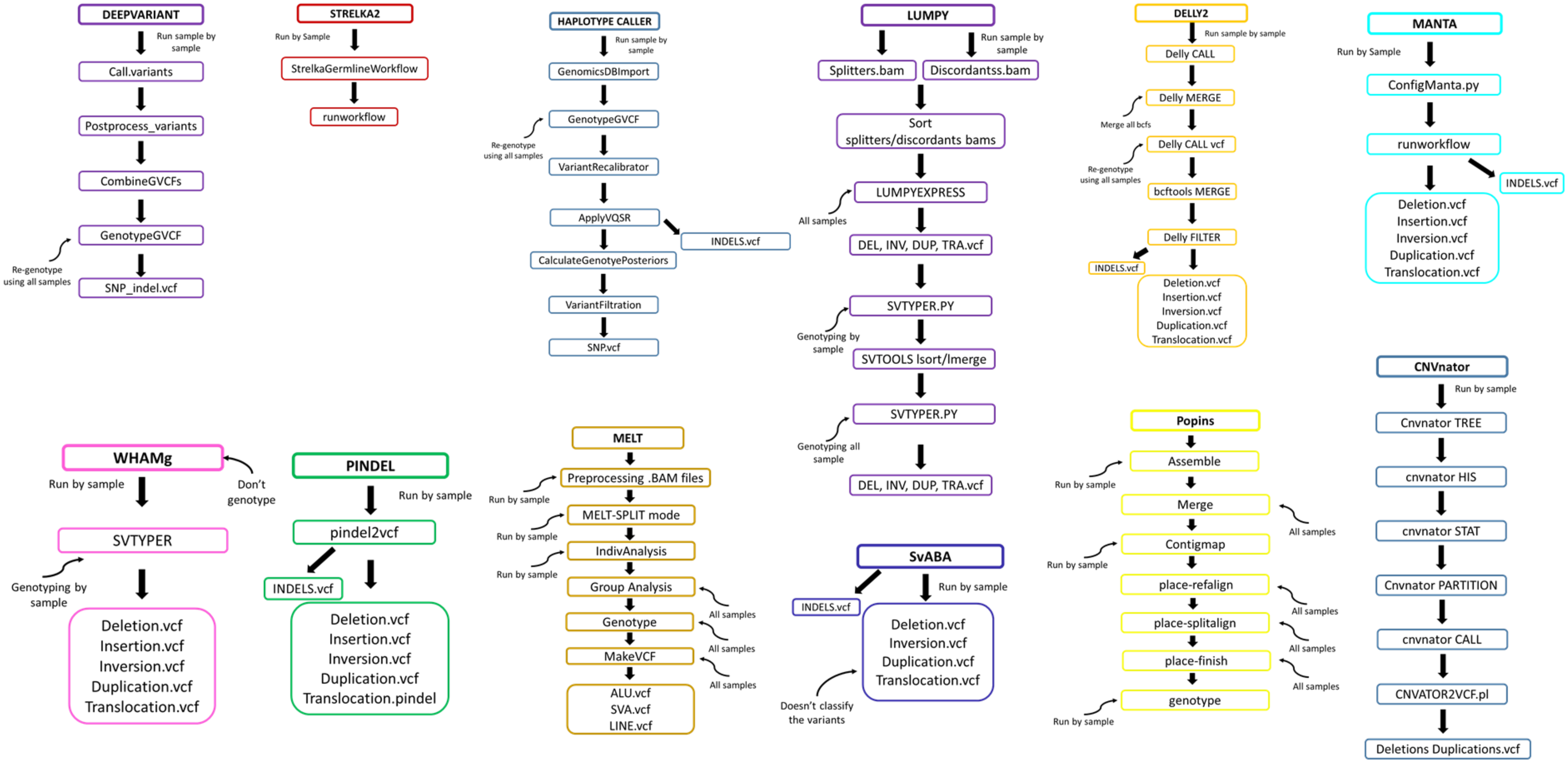

**Supplementary Fig. 8. Overview to execute each variant caller in GCAT biobank samples.**

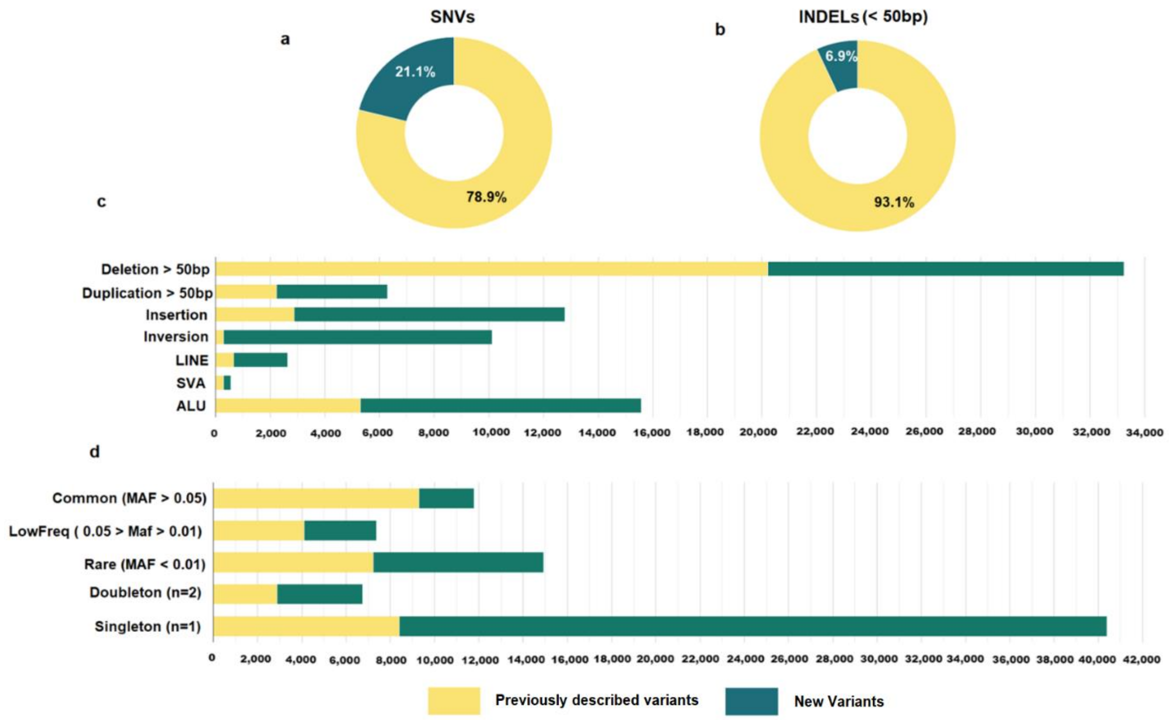

**Supplementary Fig. 9 | GCAT variant contribution in comparison to popular datasets. a,** Proportion of new and described SNVs in dbSNP (v.153). Variants were matched considering both genomic coordinates and alleles. **b,** Proportion of new and described indels (<50 bp) in dbSNP (v.153). Variants were matched considering genomic coordinates but not ref/alt states to avoid normalization errors. **c,** Proportion of new and previously described SVs against popular datasets distributed by SV type. Variants were matched by genomic coordinates considering a breakpoint error of ±1000 bp, structural variant type and if the size was available, 80% of reciprocal overlap (see Methods). **d,** Proportion of new and previously described SVs compared to popular datasets distributed by variant frequency.

a

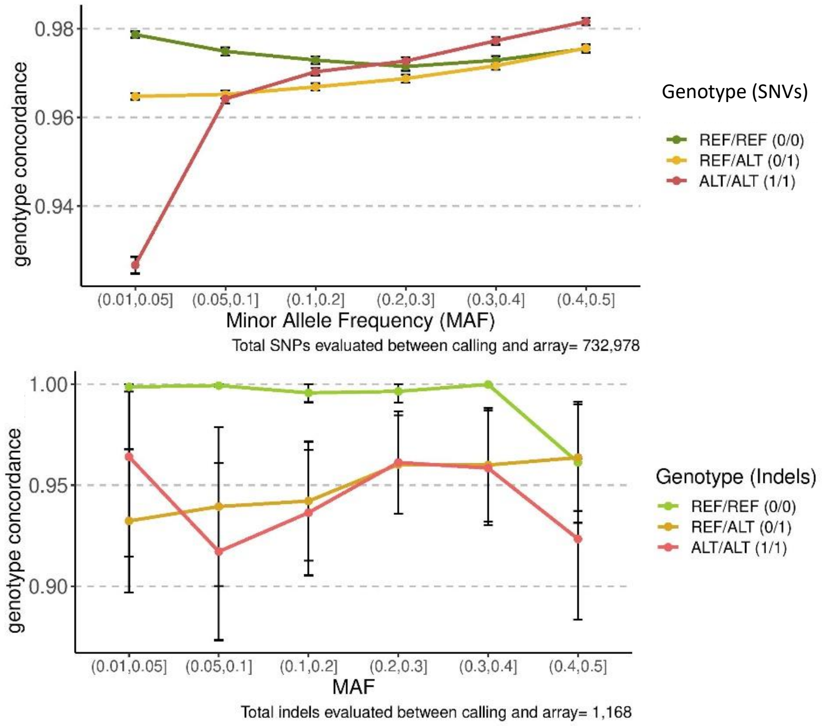

**Supplementary Fig. 10. Genotype concordance between WGS variant calling and GCAT-genotyping array for SNVs and indels.** We compared the genotypes obtained from our variant calling pipline with those from SNP-genotyping array, in order to estimate the genotype accuracy of variant callers. **a,** Genotype concordance of SNVs between variant calling and GCAT SNP-array. 96% of SNVs from SNP-genotyping array matched with WGS variant calling. Few discrepancies were observed between genotypes, being concordant 97% of all the genotype types and frequencies, with the exception of low frequency (0.01, 0.05] homozygous alternative SNVs, where the concordance decreased to < 94% **b,** Genotype concordance of indels between variant calling and GCAT SNP-array. 87% of 1,168 indels (≤ 30bp with MAF > 0.01) from SNP-genotyping array matched with WGS variant calling. 96% of alleles reported by variant calling were concordant with array genotypes, independently of allele frequency.

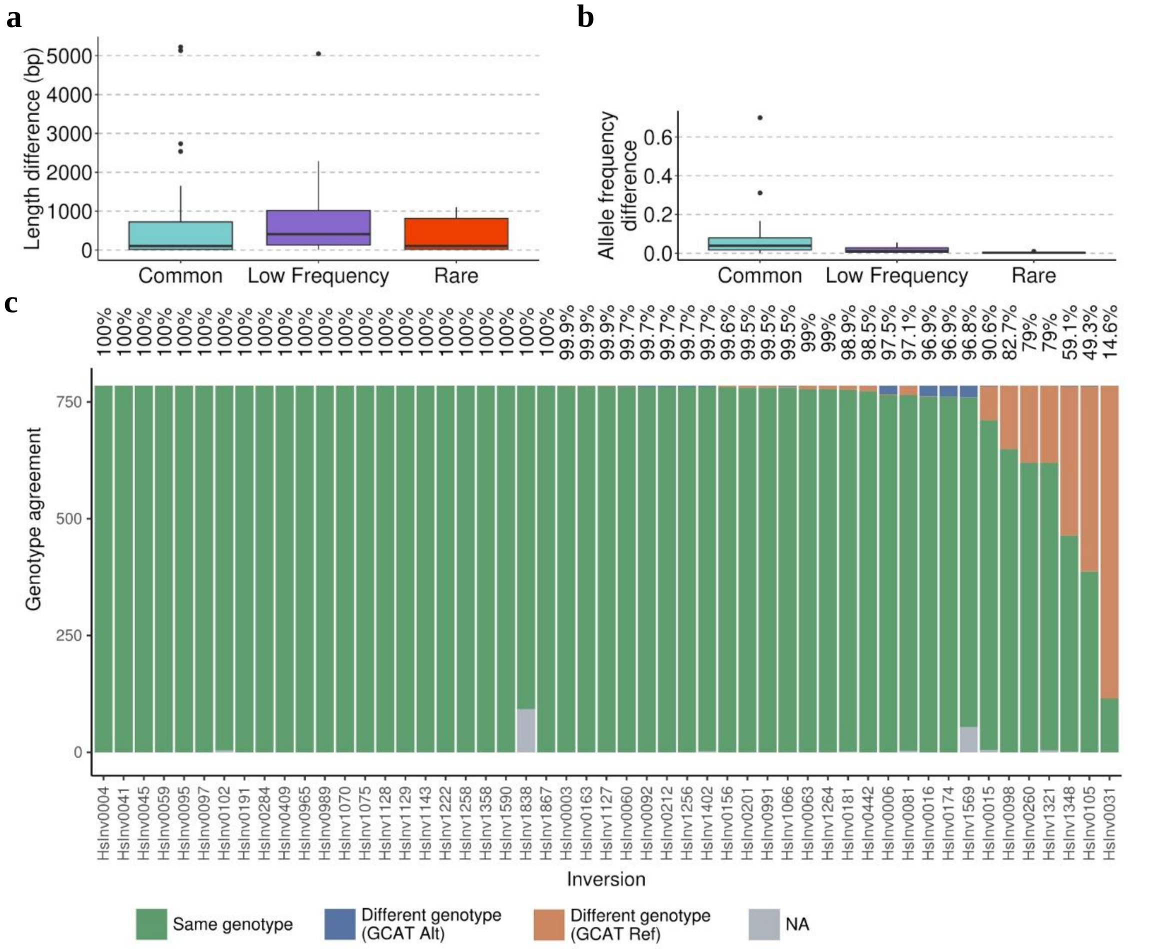

**Supplementary Fig. 11 | Validation of inversion calls using the InvFEST dataset. a,** Length discordance between the InvFEST and GCAT inversions. **b,** Allele frequency differences between 51 (MAF > 0.01) InvFEST and GCAT inversions. **c,** Genotype accuracy between GCAT variant calling and imputation calls with the InvFEST experimental panel with green bars showing agreement between both (with percentage indicated at the top). Individuals that had extra alleles with the same orientation as the reference genome in the GCAT calls were labelled as GCAT Ref (orange) whereas extra alternative orientation alleles in GCAT are indicated as GCAT Alt (blue). NA, individuals with unknown imputation call from the InvFEST panel (not considered in the comparison).

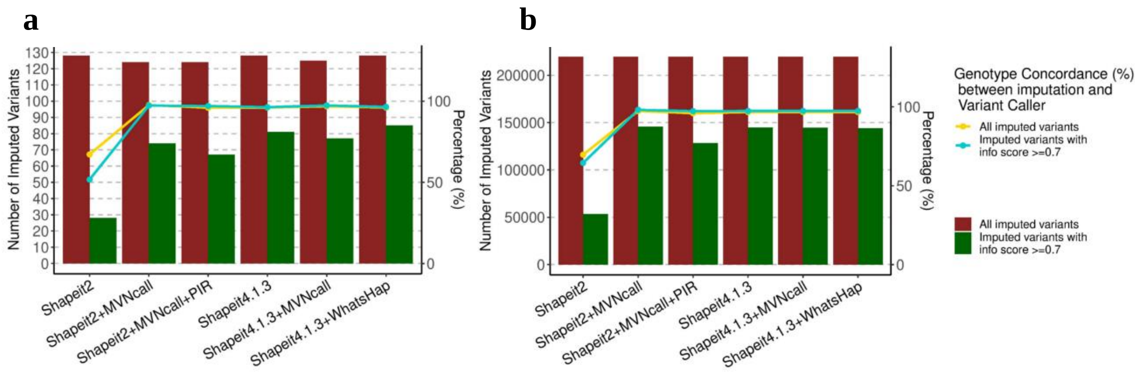

**Supplementary Fig. 12. Number of variants imputed using different phasing strategies.** The left Y axis showed the number of imputed variants, the right Y axis showed the percentage of concordant genotypes between WGS and imputation. **a,** Large deletions (≥ 50 bp) on chromosome 22 imputed with an info score ≥0.7, obtained from pilot reference panels built with different phasing strategies. **b,** SNVs and indels on chromosome 22 imputed with an info score ≥ 0.7, obtained from pilot reference panels built with different phasing strategies.

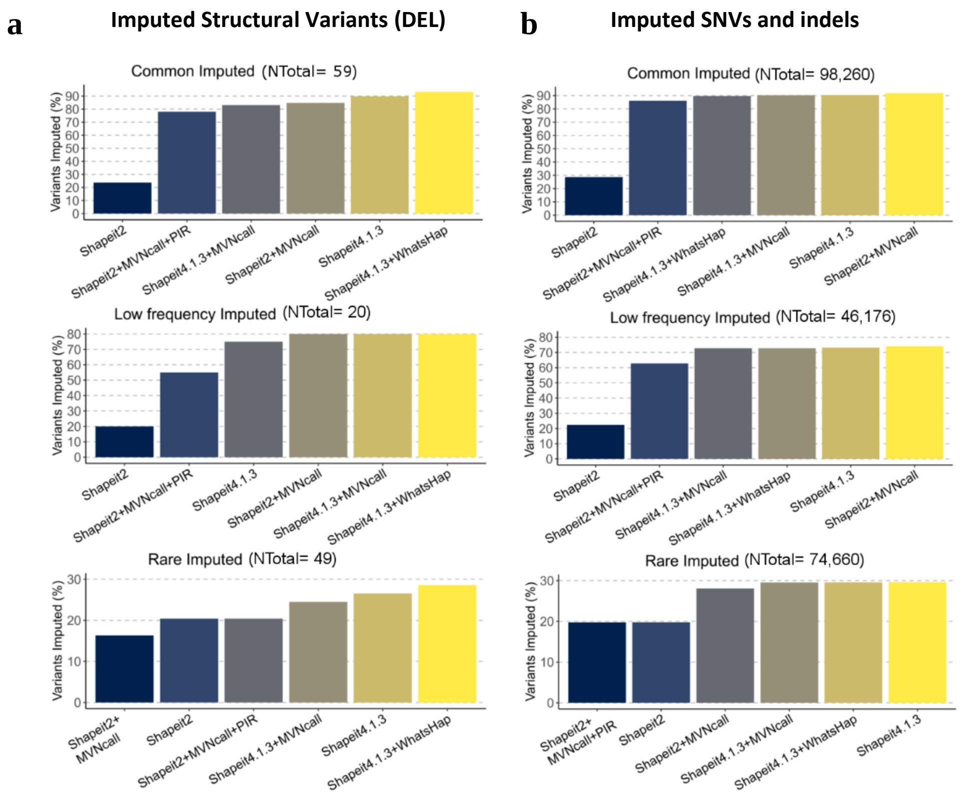

**Supplementary Fig. 13 | Benchmarking of phasing strategies in chromosome 22.** Each bar shows the phasing strategies evaluated, and sorted by number of variants imputed with info score ≥ 0.7. On the top of each graph shows the total of imputable variants in chromosome 22 panel. **a,** Shapeit4+WhatsHap improved SV imputation performance, mainly in rare. **b,** There are no remarkable differences between Shapeit2+MVNcall and all Shapeit4 combinations.

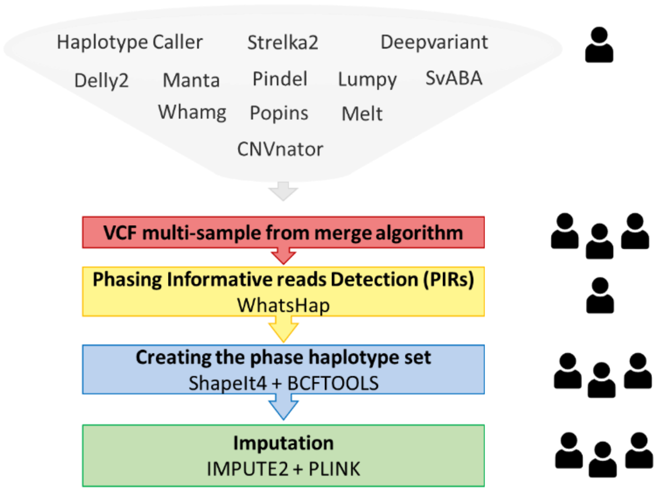

**Supplementary Fig. 14. Pipeline followed to construct the GCAT|Panel.**

Step executed sample by sample.

Step executed with all the 785 GCAT samples together.

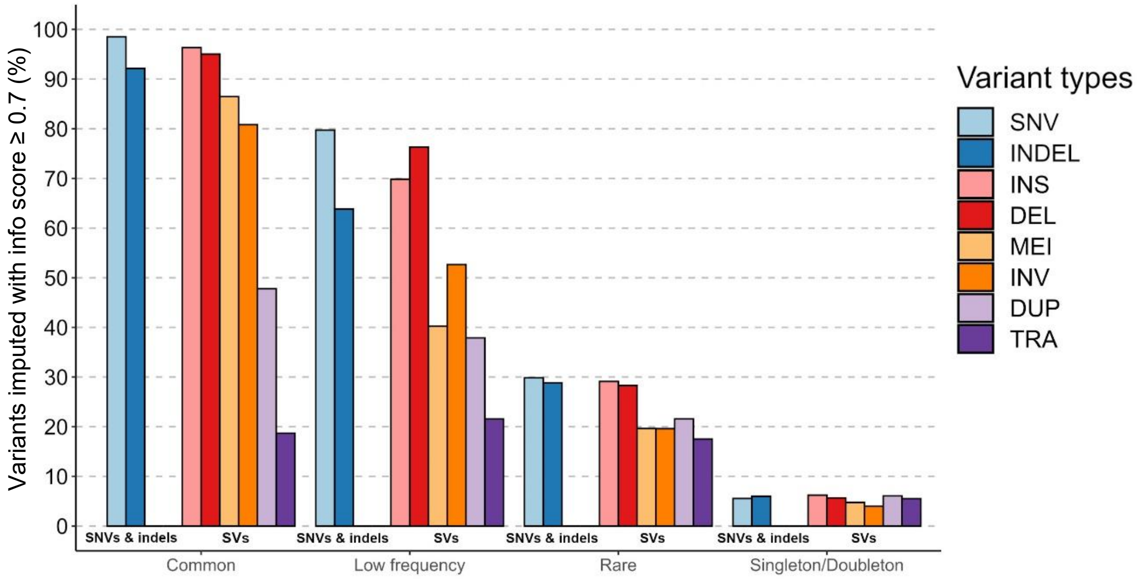

**Supplementary Fig. 15 | Imputation performance of GCAT|Panel by variant type.** The barplot shows the variants imputed with an info score ≥ 0.7 categorised by allele frequency. More than 90% of SNVs and indels and around 90% of common Structural variants (MAF ≥ 0.05) were imputed with an info score ≥0.7. The proportion of variants imputed decreased according to the allele frequency.

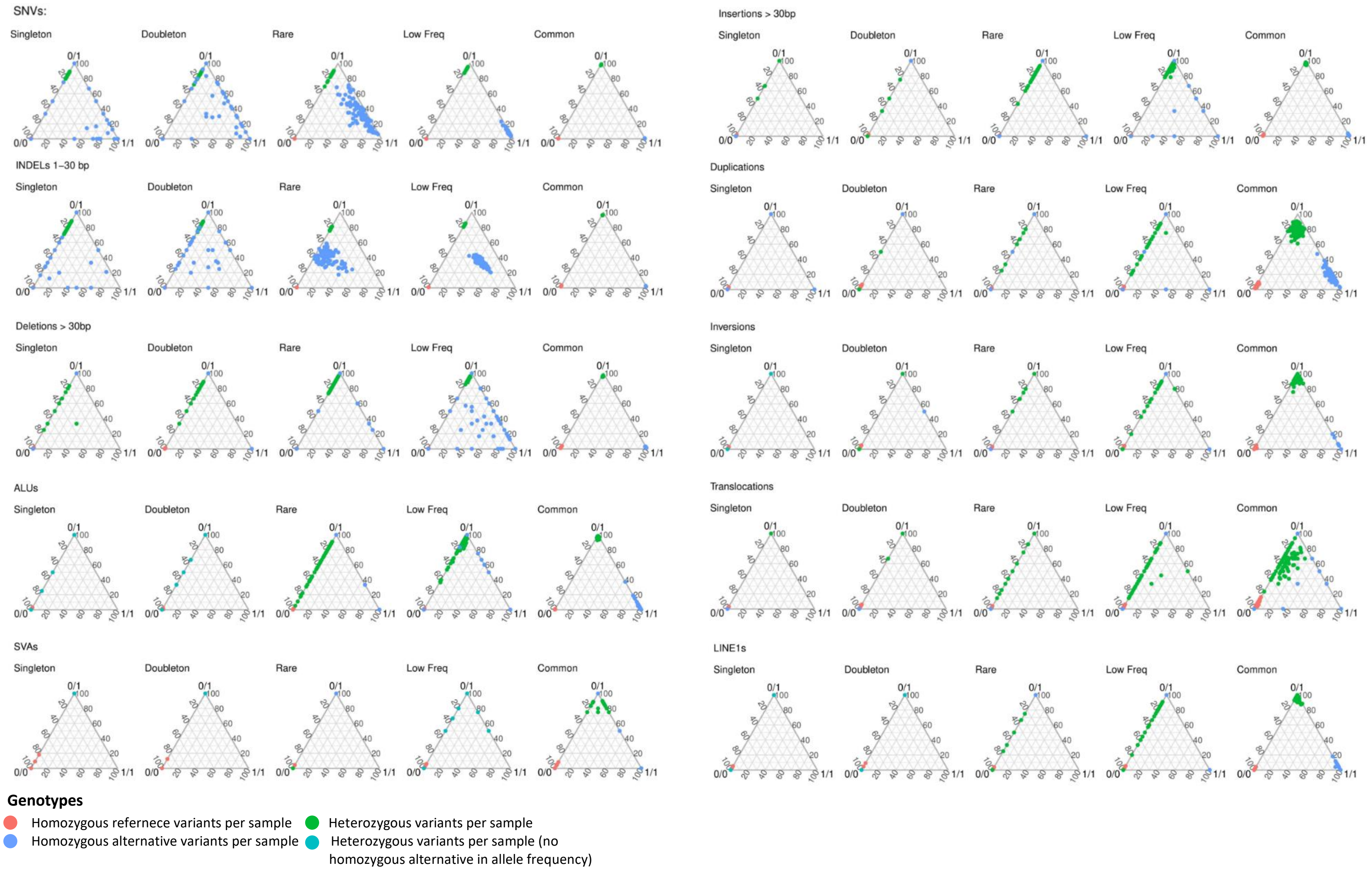

**Supplementary Fig 16. Imputation performance of the Iberian-GCAT panel for different variant types.** Genotype concordance for 95 samples, grouped by variant type and allele frequency. Each dot is the percentage of genotype concordance between genotype imputation and genotype reported by Iberian-GCAT dataset, considering all variants per sample. The genotype concordance was calculated for each genotype state independently. When a dot is in a vertice, it means high genotype concordance. For common SNVs, Indels, Deletions, Insertions, Inversions, Alus, LINE1 and SVA the genotype concordance between variant calling and imputation was 98%, except Duplications (84%) and Translocations (73%). Genotype concordance decreased with the allele frequency. For low-frequency variants ($0.01 \leq MAF < 0.05$), the homozygous alternative genotypes showed a concordance of ~40-60%, with the exception of SNVs, where concordance was ~90% for both heterozygous and homozygous alternative genotypes. Finally, for rare variants ($MAF < 0.01$), concordance for heterozygous genotypes was ~60-86%, and dropped to ~35% for homozygous alternative genotypes

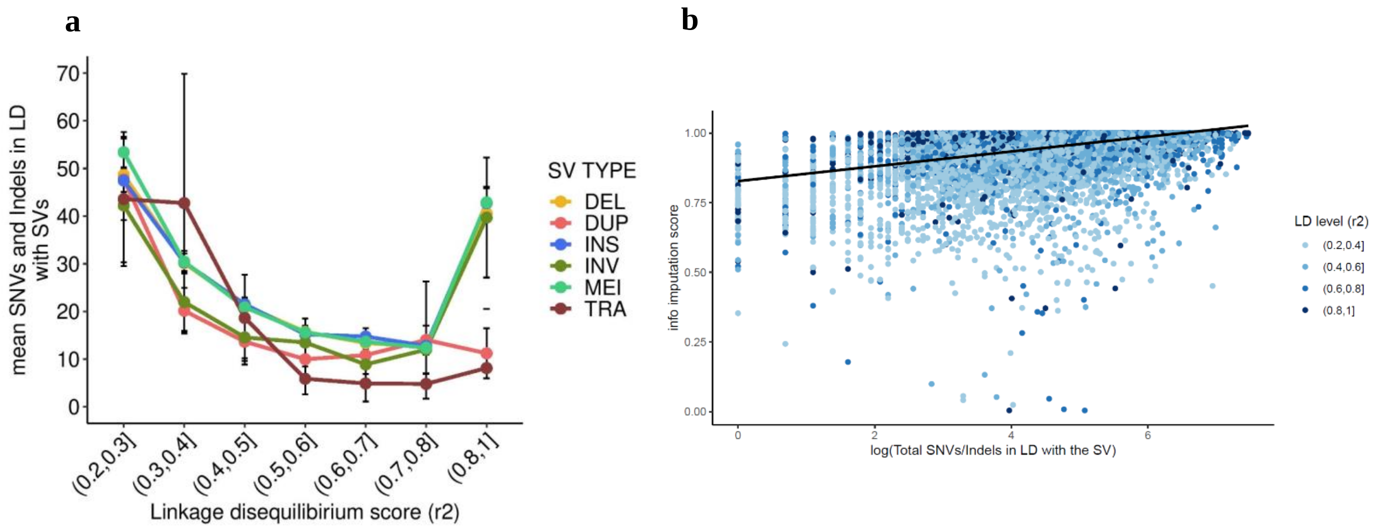

**Supplementary Fig. 17 | Imputation performance of common SVs (MAF ≥ 0.05) correlated with SNVs and indels in Linkage Disequilibrium (LD). a,** Relation between linkage disequilibrium score and number of SNVs and indels in LD by SV type. On average 40 SNVs and indels have high LD ($r^2 > 0.8$) with deletions (DEL), inversions (INV), mobile element insertions (MEI) and *de novo* insertions (INS). **b,** Scatterplot of the number of SNVs/Indels in LD (in log scale) and the info imputation score per SV. The median of LD level of all the SNVs/Indels is represented in blue colour by bins of (0.2,0.4], (0.4,0.6], (0.6,0.8] and (0.8,1] LD ($r^2$). The fitted regression line is represented by a black line showing a positive Pearson correlation coefficient, r = 0.38 (95%CI = [0.37,0.40], p-value < $2x10^{-16}$).

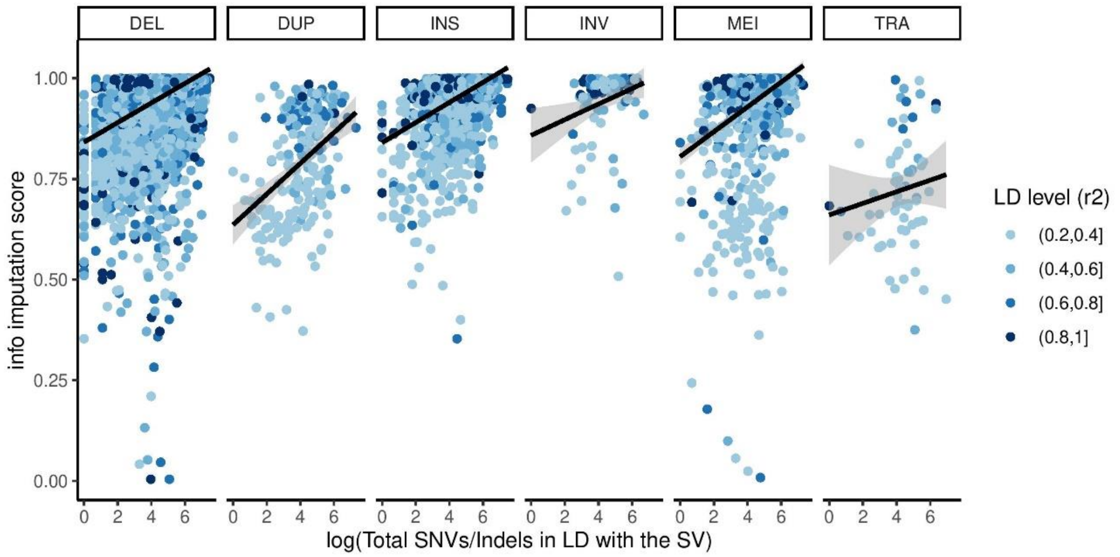

**Supplementary Fig. 18. Scatterplot of the number of SNVs/Indels in LD with the SV (in log scale) and the info imputation score of the SV for each SV type.** The median of LD of all the SNVs/Indels is represented in blue color by bins of (0.2,0.4], (0.4,0.6], (0.6,0.8] and (0.8,1] LD levels (r2). The fitted regression line is represented by a black line and confidence intervals in gray color for each SV type independently. Deletions (Pearson's r = 0.39, df = 9746, 95% CI = [0.38,0.41], p-value < $2x10^{-16}$), Insertions (Pearson's r = 0.44, df = 2511, 95% CI = [0.41,0.47], p-value < $2x10^{-16}$), Duplications (Pearson's r = 0.41, df = 217, 95% CI = [0.30,0.52], p-value = $1.2x10^{-8}$), Inversions (Pearson's r = 0.26, df = 98, 95% CI = [0.06,0.43], p-value = 0.008) and Translocations (Pearson's r = 0.13, df = 61, 95% CI = [-0.12,0.36], p-value = 0.31), with Translocations the unique type without significat correlation.

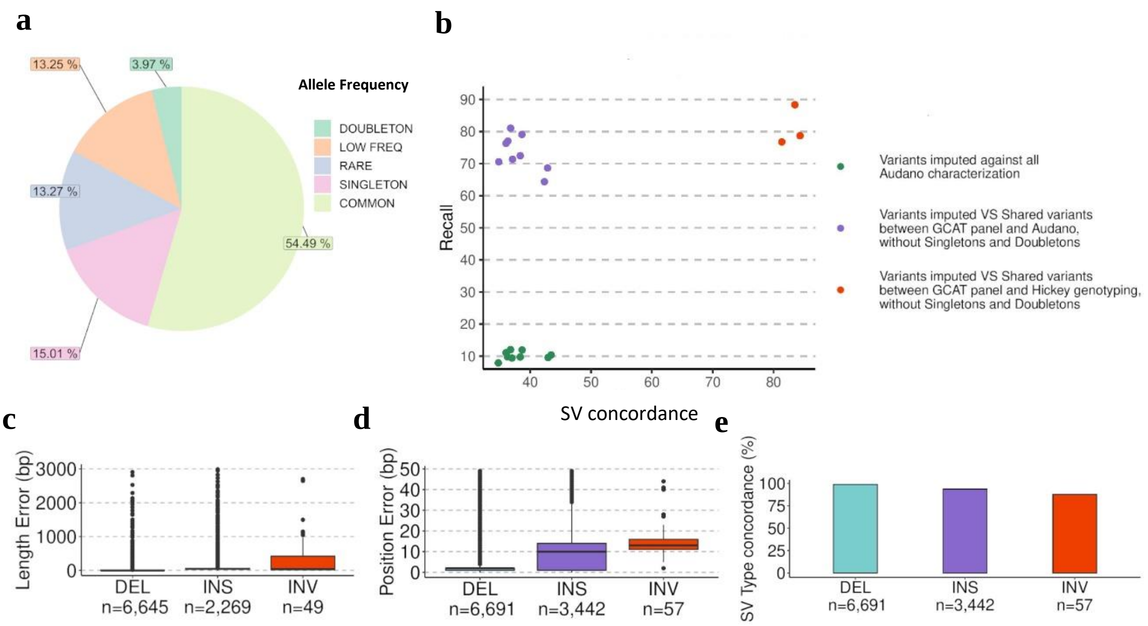

**Supplementary Fig. 19. Imputation performance of the GCAT|Panel. a,** Allele frequency distribution of the 16,704 SVs shared between the Audano et al and GCAT SV dataset. The majority of which were common (54.5%), followed by singletons+doubletons (18.9%), rare (13.3%), and low-frequency (13.3%). **b,** Comparison of imputation results using the SVs detected by Audano et al.[15] and genotypes from Hickey et al[32]. Considering all SVs detected by Audano et al.[15], 10% of them were recovered by GCAT|Panel (green dots). From this 73% were imputed using 1000G array (purple dots). Hickey et al[32]. genotyped three of nine samples using short-reads, increasing both SV concordance (83%) and recall (81%). **c,d,e,** Length error, breakpoint resolution and variant type reported by the GCAT SV catalogue and Audano et al. was consistent in all categories.

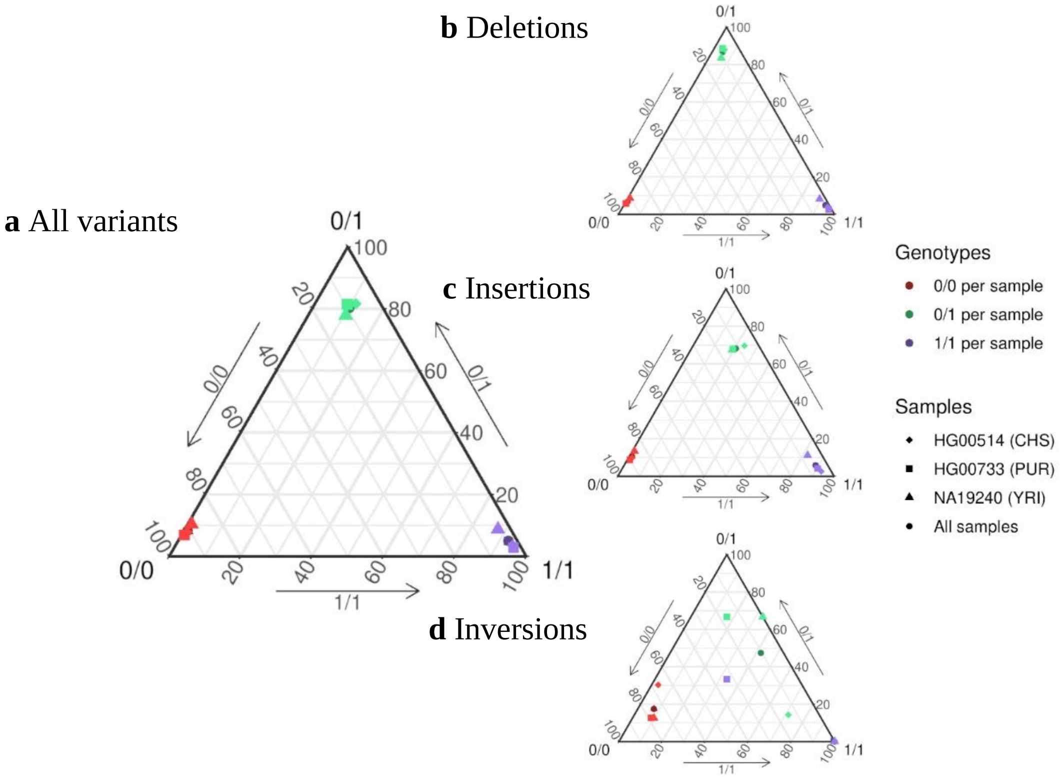

**Supplementary Fig. 20. Genotype concordance of structural variants between imputation and Hickey et al genotyping.** Genotype concordance was calculated using all variants for each sample, obtaining proportion of concordance between SV imputation results and Hickey et al. **a,** Ternary diagram of genotype imputation accuracy using three genotyped samples from Hickey et al[32]. The samples with high genotype concordances between both approaches are close to the ternary diagram vertices. **b,** Genotype concordance of Deletions (n=17,205). Deletions were the SVs imputed with highest concordance (~90%). **c,** Genotype concordance of Insertions (n=8,903). The insertion genotypes obtained from imputation with the Iberian-GCAT reference panel, comprising TRAs, DUPs, TRPs and INSs**,** were ~70% concordant in heterozygous and ~90% in homozygous with those reported by Hickey et al., showing a good genotype concordance. **d,** Genotype concordance of Inversions (n=141). Small number of inversions were shared between the two projects, hampering the possibility of obtaining deeper insights into this poorer concordance.

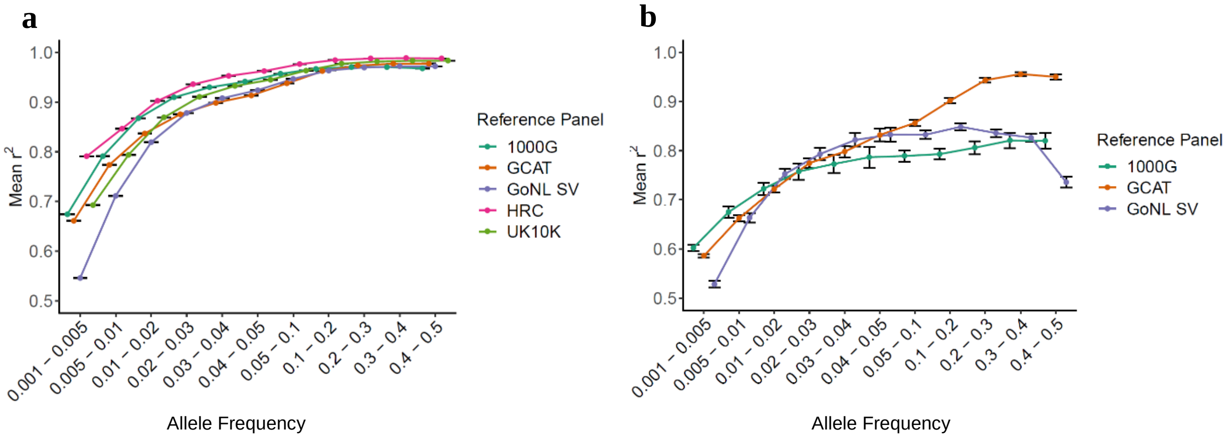

**Supplementary Fig. 21. Quality of variants imputed (info score) using different reference panels. a,** Imputation accuracy of SNVs and indels at different allele frequencies. HRC obtained the best results due to the panel sample size, allowing to impute also rare variants with high quality. **b,** The imputation quality among different panels showed slightly differencies for low allele frequencies (MAF < 0.05). However, the GCAT|Panel showed a higher imputation quality for common SVs in comparison to other panels.

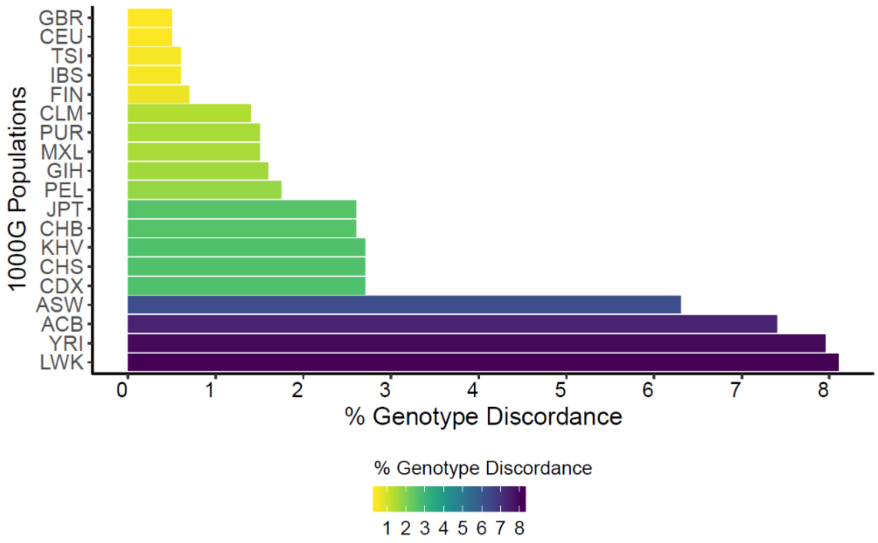

**Supplementary Fig. 22. Imputation quality grouped by populations.** Imputation quality was overall high in all populations, suggesting that the GCAT|Panel could be used in all populations for imputation analysis, even in African populations, where the imputation quality is lower. As expected, imputation quality was strongly correlated with the genetic similarity between populations and Iberians. Indeed, European ancestries showed a genotype discordance ≤ 1% in European ancestries, followed by Latin American ancestries (and Indian population (GIH)) (< 2% discordance), Asian ancestries (< 3% discordance) and African ancestries, which genotype discordance increased to > 6%.

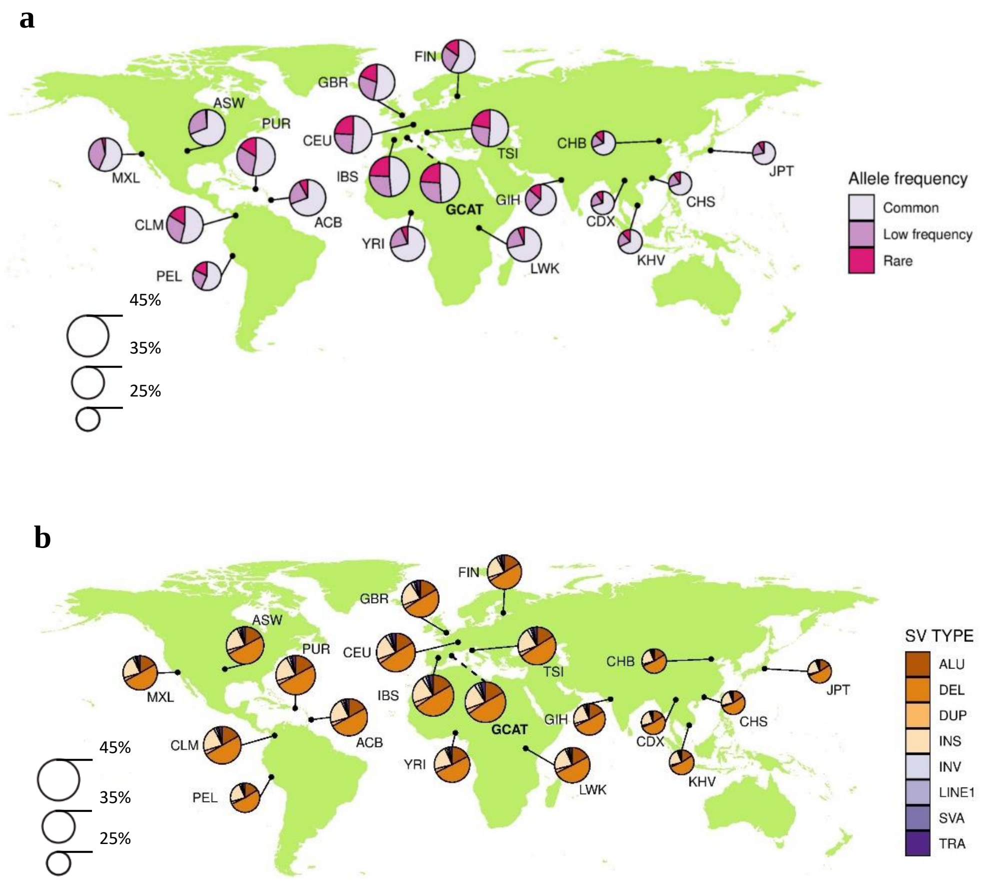

**Supplementary Fig. 23. Structural variant distribution by Population.** A total of 49,724 SVs were recovered with an info score ≥ 0.7 after impute the 1000G array for each population independently (19 populations). **a,** Allele Frequency Distribution by population. For Asian populations (with the exception of the Indian ones (GIH)), we were able to recover fewer SVs than other continents (~25%). Between 35 and 40% of all the imputed SVs were instead present in the African and Latin American populations. The European populations carried (with the exception of Finnish (FIN)) > 40% of the imputed SVs. Additionally, the majority of imputed variants were common (MAF ≥ 0.05), with ~70% in Asian and African populations, and ~50% in Europeans and Latin Americans. **b,** SV type recovered and distributed by population. Deletions were the most imputed SVs across all populations (~50%), followed by insertions (~22%) and Alus (~17%). The other SVs represented ~10% of all imputed SVs per population. These results indicate that imputation performance is correlated with ancestries.

Further description of 19 populations in Supplementary Table 14.

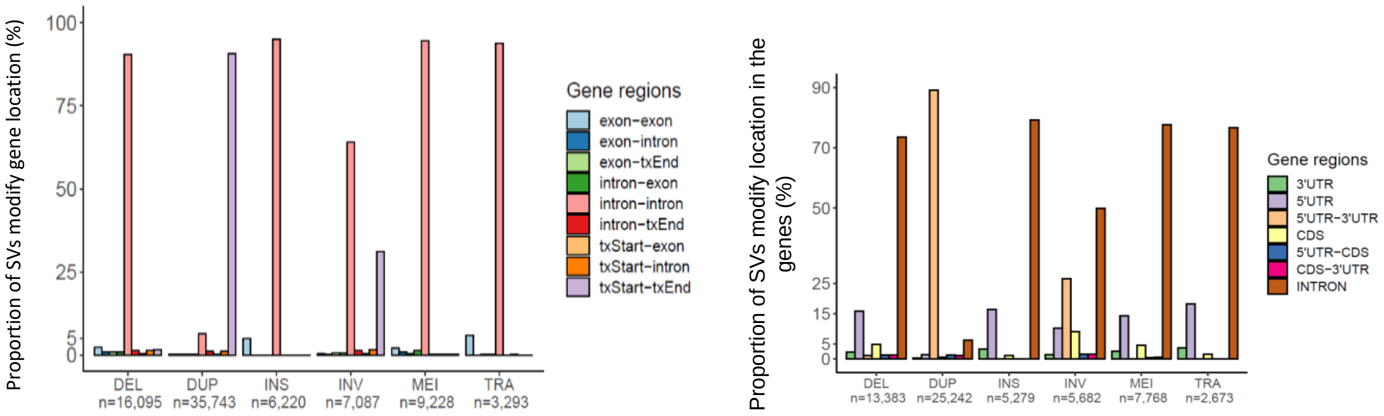

**Supplementary Fig. 24. Structural Variant annotation by type in different gene regions. a,** The barplot shows the SV distribution in different gene regions using AnnotSV. SVs overlapped a gene 77,666 times, with 88% of SVs matching in intronic regions. Duplications and inversions, due to their sizes, modified a large fraction of transcripts (Tx). **b,** Structural Variant overlapping in protein-coding genes. Distribution of SVs in protein-coding genes regions. Duplications modified more genes, and the majority of SVs overlapped intronic regions (> 70%). Besides, 7,695 SVs affected UTR regions, with an enrichment of 10-18% in 5'UTR, except for duplications. In the GCAT project, 3,135 SVs modified just CDS regions, with deletions as the predominant (33%), followed by duplications (27%), inversions (23%), MEIs (13%), and a small fraction of the other SV types.

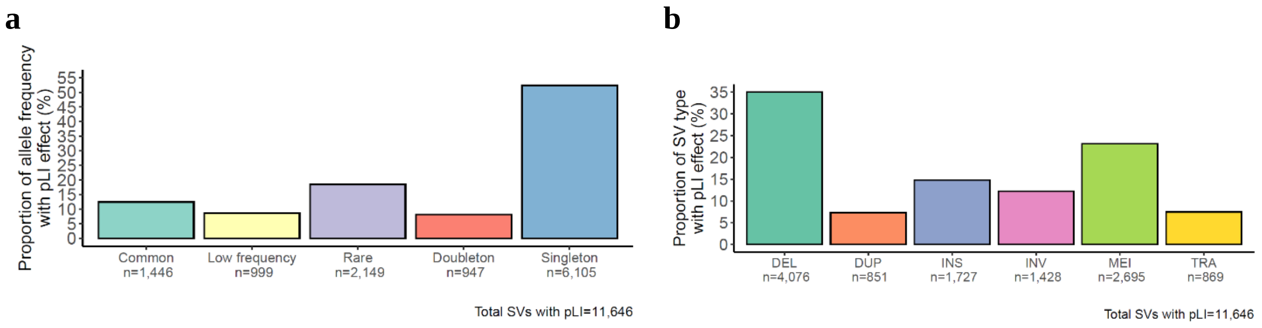

**Supplementary Fig. 25. Deleterious effects of SVs in GCAT cohort. a,** Prediction of loss of function intolerance (pLI) categorised by allele frequency. SVs with MAFs < 0.01 represented 79% of all pLI genes, suggesting a selective pressure to variants with high gene function impact. However, 21% of SV were MAF ≥ 0.01, indicating different pathogenic levels. **b,** Prediction loss of function intolerance, categorised by SV type. Deletions and Mobile Element Insertions (MEIs) were more deleterious than duplications and translocations.

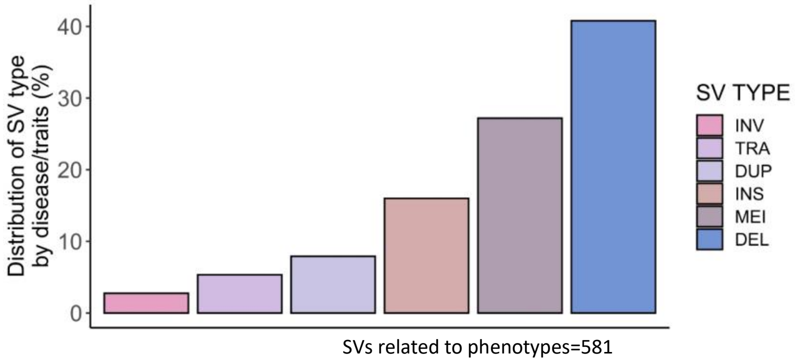

**Supplementary Fig. 26. Structural variants prediction effect on diseases using OMIM database.** Structural variants related with a diseases in OMIM database. The SVs selected were those which overlapped in genes with pLI ≥ 0.9 and Haploinsufficiency effect. Deletions were the most associated with diseases (41%).

Diseases associated with SV types are in Supplementary Table 15.

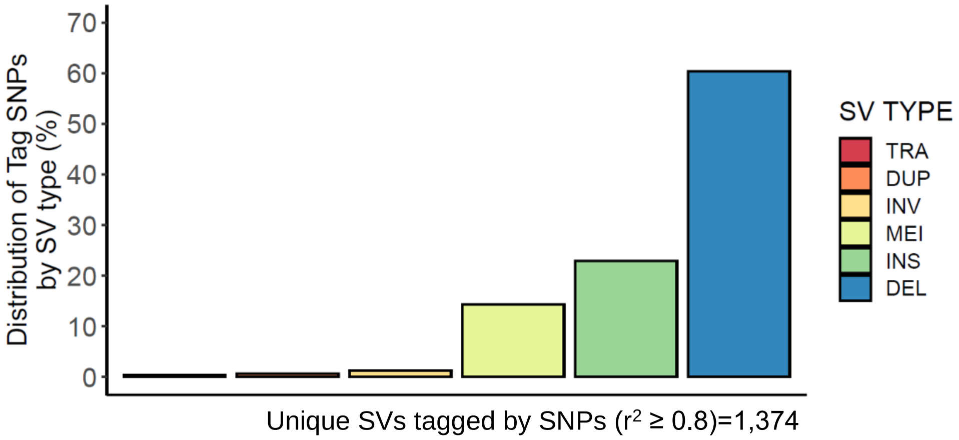

**Supplementary Fig. 27. Structural variants tagged by SNPs from GWAS catalog with high Linkage disequilibrium (LD ≥ 0.8)**. The SNPs from the GWAS catalog tagged with high LD were 3.72% of the SV in the GCAT dataset (36,887 SVs (MAF ≥ 0.01)). The predominant type variants were deletions, insertions, and MEIs.

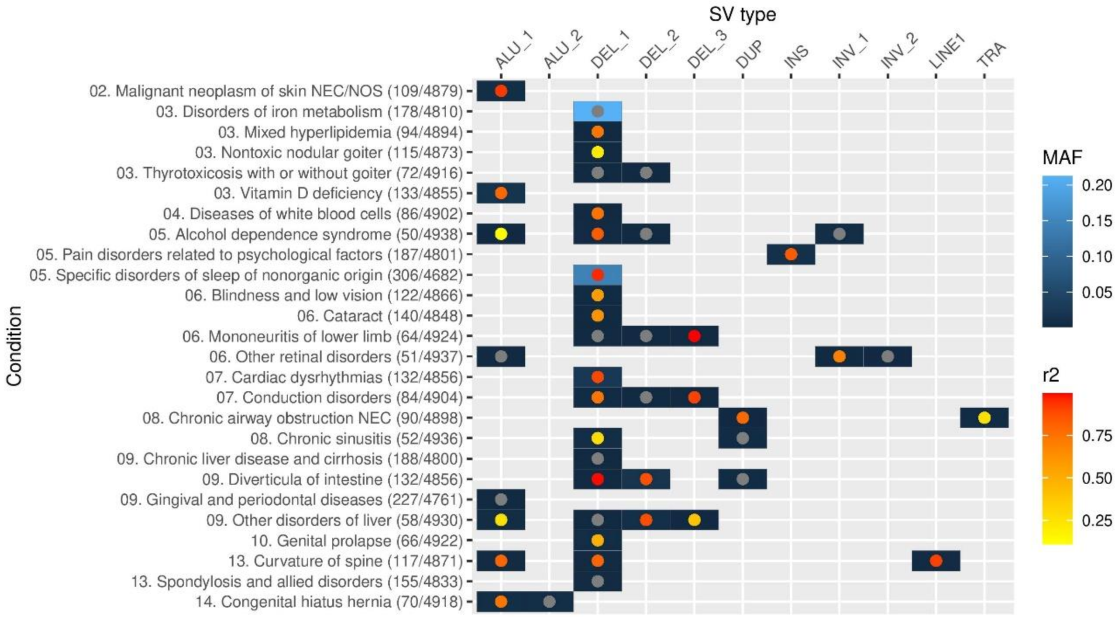

**Supplementary Fig. 28 | Genome Wide Association analysis.** GWAS results of suggestive loci (p-value < 1 x$10^{-6}$) harboring a structural variant by condition and type. Each blue square represents the SV identified for a particular condition. When more than one SV of a particular type is identified for a condition, additional squares are represented in the X axis with successive numbers. ICD-9 Chapter and its description with the number of cases and controls is represented in the Y axis. The intensity of the blue square colour represents the MAF of the variant. The colour intensity of the points represents the linkage disequilibrium ($r^2$) between the structural variant and the leading variant in a 100 Kb window. Those SV being the leading variant are marked in grey.

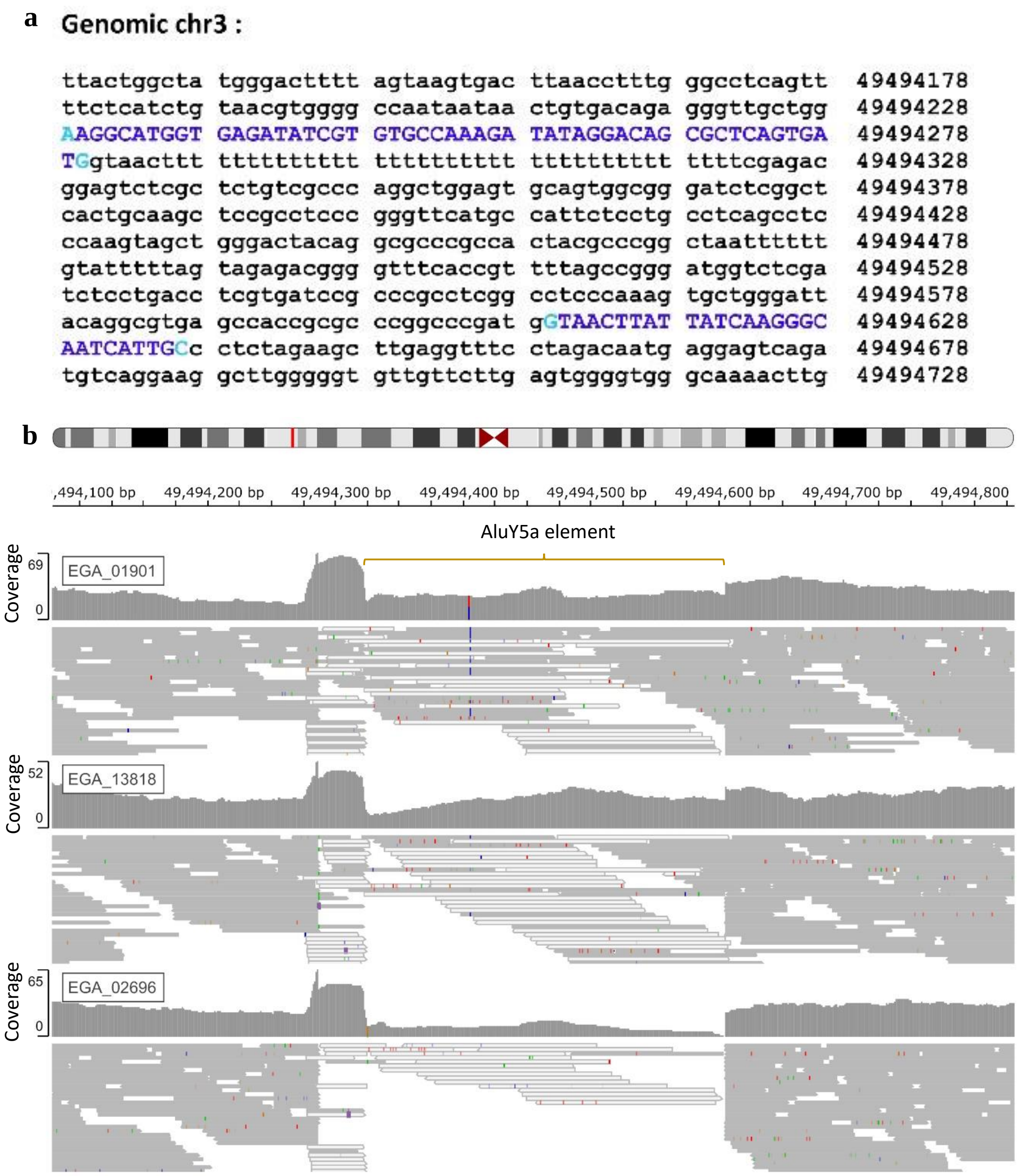

**Supplementary Fig. 29 | Experimental validation of the AluY5a element. a**, Insertion point of the Alu-element. BLAT analysis of the control sequence in a non-Alu-allele individual indicates the insertion point at Chr3:49,494,280. In blue are the sanger sequences of the control individual. **c**, IGV of the Alu region in two carriers and in one control. Samples EGA_01901 and EGA_13818 are heterozygous for the Alu insertion, and EGA_02696 is homozygous for the control allele. Sample EGA_01901 shows in the IGV was genotyped and imputed and confirmed experimentally by PCR and Sanger sequence. Reads with map quality of 0 are represented in white.
